# Projecting genetic associations through gene expression patterns highlights disease etiology and drug mechanisms

**DOI:** 10.1101/2021.07.05.450786

**Authors:** Milton Pividori, Sumei Lu, Binglan Li, Chun Su, Matthew E. Johnson, Wei-Qi Wei, Qiping Feng, Bahram Namjou, Krzysztof Kiryluk, Iftikhar Kullo, Yuan Luo, Blair D. Sullivan, Benjamin F. Voight, Carsten Skarke, Marylyn D. Ritchie, Struan F.A. Grant, Casey S. Greene

## Abstract

Genes act in concert with each other in specific contexts to perform their functions. Determining how these genes influence complex traits requires a mechanistic understanding of expression regulation across different conditions. It has been shown that this insight is critical for developing new therapies. In this regard, the role of individual genes in disease-relevant mechanisms can be hypothesized with transcriptome-wide association studies (TWAS), which have represented a significant step forward in testing the mediating role of gene expression in GWAS associations. However, modern models of the architecture of complex traits predict that gene-gene interactions play a crucial role in disease origin and progression. Here we introduce PhenoPLIER, a computational approach that maps gene-trait associations and pharmacological perturbation data into a common latent representation for a joint analysis. This representation is based on modules of genes with similar expression patterns across the same conditions. We observed that diseases were significantly associated with gene modules expressed in relevant cell types, and our approach was accurate in predicting known drug-disease pairs and inferring mechanisms of action. Furthermore, using a CRISPR screen to analyze lipid regulation, we found that functionally important players lacked TWAS associations but were prioritized in trait-associated modules by PhenoPLIER. By incorporating groups of co-expressed genes, PhenoPLIER can contextualize genetic associations and reveal potential targets missed by single-gene strategies.

## Introduction

Genes work together in context-specific networks to carry out different functions [1,2]. Variations in these genes can change their functional role and, at a higher level, affect disease-relevant biological processes [3]. In this context, determining how genes influence complex traits requires mechanistically understanding expression regulation across different cell types [4,5,6], which in turn should lead to improved treatments [7,8]. Previous studies have described different regulatory DNA elements [5,9,10,11,12] including genetic effects on gene expression across different tissues [4]. Integrating functional genomics data and GWAS data [13,13,14,15] has improved the identification of these transcriptional mechanisms that, when dysregulated, commonly result in tissue- and cell lineage-specific pathology [16,17,18].

Given the availability of gene expression data across several tissues [4,19,20,21], an effective approach to identify these biological processes is the transcription-wide association study (TWAS), which integrates expression quantitative trait loci (eQTLs) data to provide a mechanistic interpretation for GWAS findings. TWAS relies on testing whether perturbations in gene regulatory mechanisms mediate the association between genetic variants and human diseases [22,23,24,25], and these approaches have been highly successful not only in understanding disease etiology at the transcriptome level [26,27,28] but also in disease-risk prediction (polygenic scores) [29] and drug repurposing [30] tasks. However, TWAS works at the individual gene level, which does not capture more complex interactions at the network level.

These gene-gene interactions play a crucial role in current theories of the architecture of complex traits, such as the omnigenic model [31], which suggests that methods need to incorporate this complexity to disentangle disease-relevant mechanisms. Widespread gene pleiotropy, for instance, reveals the highly interconnected nature of transcriptional networks [32,33], where potentially all genes expressed in disease-relevant cell types have a non-zero effect on the trait [31,34]. One way to learn these gene-gene interactions is using the concept of gene module: a group of genes with similar expression profiles across different conditions [2,35,36]. In this context, several unsupervised approaches have been proposed to infer these gene-gene connections by extracting gene modules from co-expression patterns [37,38,39]. Matrix factorization techniques like independent or principal component analysis (ICA/PCA) have shown superior performance in this task [40] since they capture local expression effects from a subset of samples and can handle modules overlap effectively. Therefore, integrating genetic studies with gene modules extracted using unsupervised learning could further improve our understanding of disease origin [36] and progression [41].

Here we propose PhenoPLIER, an omnigenic approach that provides a gene module perspective to genetic studies. The flexibility of our method allows integrating different data modalities into the same representation for a joint analysis. In this work, we show that this module perspective can infer how groups of functionally-related genes influence complex traits, detect shared and distinct transcriptomic properties among traits, and predict how pharmacological perturbations affect genes’ activity to exert their effects. PhenoPLIER maps gene-trait associations and drug-induced transcriptional responses into a common latent representation. For this, we integrated thousands of gene-trait associations (using TWAS from PhenomeXcan [42]) and transcriptional profiles of drugs (from LINCS L1000 [43]) into a low-dimensional space learned from public gene expression data on tens of thousands of RNA-seq samples (recount2 [19,44]). We used a latent representation defined by a matrix factorization approach [44,45] that extracts gene modules with certain sparsity constraints and preferences for those that align with prior knowledge (pathways). When mapping gene-trait associations to this reduced expression space, we observed that diseases were significantly associated with gene modules expressed in relevant cell types: such as hypothyroidism with T cells, corneal endothelial cells with keratometry measurements, hematological assays on specific blood cell types, plasma lipids with adipose tissue, and neuropsychiatric disorders with different brain cell types. Moreover, since PhenoPLIER can use models derived from large and heterogeneous RNA-seq datasets, we could also identify modules associated with cell types under specific stimuli or disease states. We observed that significant module-trait associations in PhenomeXcan (our discovery cohort) replicated in the Electronic Medical Records and Genomics (eMERGE) network phase III [27,46] (our replication cohort). Furthermore, we performed a CRISPR screen to analyze lipid regulation in HepG2 cells. We observed more robust trait associations with modules than with individual genes, even when single genes known to be involved in lipid metabolism did not reach genome-wide significance. Compared to a single-gene approach, our module-based method also better predicted FDA-approved drug-disease links by capturing tissue-specific pathophysiological mechanisms linked with the mechanism of action of drugs (e.g., niacin with cardiovascular traits via a known immune mechanism). This improved drug-disease prediction suggested that modules may provide a better means to examine drug-disease relationships than individual genes. Finally, exploring the phenotype-module space revealed stable trait clusters associated with relevant tissues, including a complex branch involving lipids with cardiovascular, autoimmune, and neuropsychiatric disorders. In summary, instead of considering single genes associated with different complex traits, PhenoPLIER incorporates groups of genes that act together to carry out different functions in specific cell types. This approach improves robustness in detecting and interpreting genetic associations, and here we show how it can prioritize alternative and potentially more promising candidate targets even when known single gene associations are not detected. The approach represents a conceptual shift in the interpretation of genetic studies. It has the potential to extract mechanistic insight from statistical associations to enhance the understanding of complex diseases and their therapeutic modalities.

## Results

### PhenoPLIER: an integration framework based on gene co-expression patterns

PhenoPLIER is a flexible computational framework that combines gene-trait and gene-drug associations with gene modules expressed in specific contexts (Figure 1a). The approach uses a latent representation (with latent variables or LVs representing gene modules) derived from a large gene expression compendium (Figure 1b, top) to integrate TWAS with drug-induced transcriptional responses (Figure 1b, bottom) for a joint analysis. The approach consists in three main components (Figure 1b, middle, see Methods): 1) an LV-based regression model to compute an association between an LV and a trait, 2) a clustering framework to learn groups of traits with shared transcriptomic properties, and 3) an LV-based drug repurposing approach that links diseases to potential treatments. We performed extensive simulations for our regression model (Supplementary Note 1) and clustering framework (Supplementary Note 2) to ensure proper calibration and expected results under a model of no association.

**Figure 1:**
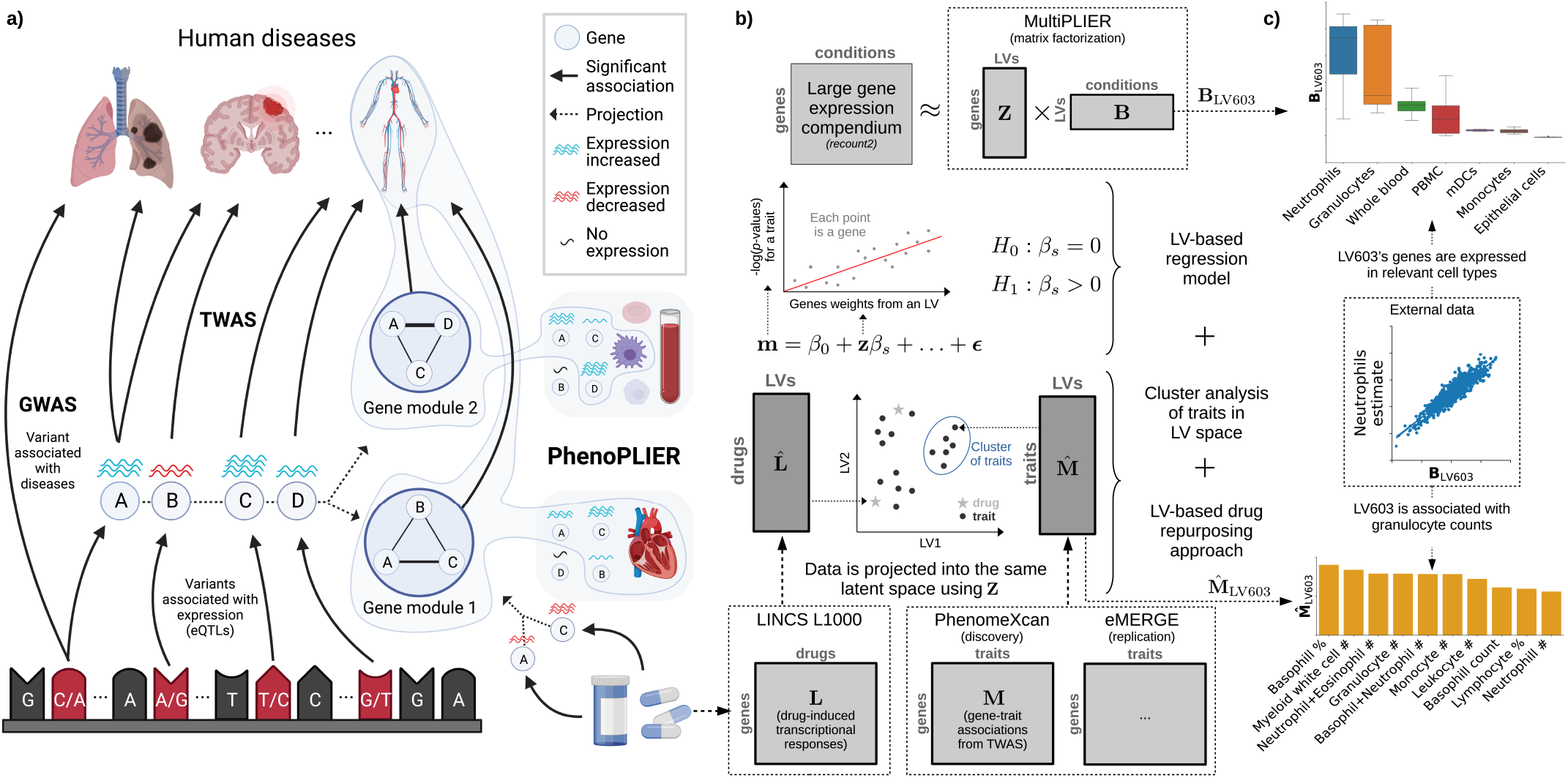
Schematic of the PhenoPLIER framework. **a)** High-level schematic of PhenoPLIER (a gene module-based method) in the context of TWAS (single-gene) and GWAS (genetic variants). PhenoPLIER integrates groups of genes co-expressed in specific cell types (gene modules) with gene-trait and gene-drug associations. **b)** The integration consists of projecting gene-trait/gene-drug associations from PhenomeXcan/LINCS L1000 (bottom) to a latent space based on gene modules (represented by latent variables/LVs) from MultiPLIER (top). This process generates matrices 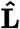 and 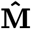, where LVs now describe each drug/trait. In the middle, we show the three main computational components provided by PhenoPLIER to perform this integration: 1) an LV-based regression model, 2) a clustering framework to learn groups of traits, and 3) an LV-based drug repurposing approach. **c)** LV603, termed as a neutrophil signature in the original MultiPLIER study, was associated in PhenoPLIER with neutrophil counts and other white blood cells (bottom, showing the top 10 traits for LV603 after projecting gene-trait associations in PhenomeXcan). Genes that are part of LV603 were expressed in relevant cell types (top). PBMC: peripheral blood mononuclear cells; mDCs: myeloid dendritic cells.

We used TWAS results from PhenomeXcan [42] and the eMERGE network [27] as discovery and replication cohorts, respectively (Methods). PhenomeXcan provides gene-trait associations for 4,091 different diseases and traits from the UK Biobank [47] and other studies, whereas the analyses on eMERGE were performed across 309 phecodes. TWAS results were derived using two statistical methods (see Methods): 1) Summary-MultiXcan (S-MultiXcan) associations were used for the regression and clustering components, and 2) Summary-PrediXcan (S-PrediXcan) associations were used for the drug repurposing component. In addition, we also used colocalization results, which provide a probability of overlap between the GWAS and eQTL signals. For the drug-repurposing approach, we used transcriptional responses to small molecule perturbations from LINCS L1000 [43] comprising 1,170 compounds.

The latent gene expression representation was obtained from the MultiPLIER models [44], which were derived by applying a matrix factorization method (the pathway-level information extractor or PLIER [45]) to recount2 [19] – a uniformly-curated collection of transcript-level gene expression quantified by RNA-seq in a large, diverse set of samples collected across a range of disease states, cell types differentiation stages, and various stimuli (see Methods). The MultiPLIER models extracted 987 LVs by optimizing data reconstruction but also the alignment of LVs with prior knowledge/pathways.

Each LV or gene module represents a group of weighted genes expressed together in the same tissues and cell types as a functional unit. Since LVs might represent a functional set of genes regulated by the same transcriptional program [48,49], we conjecture that the projection of TWAS and pharmacologic perturbations data into this latent space could provide a better mechanistic understanding. For this projection of different data modalities into the same space, PhenoPLIER converts gene associations to an LV score: all genes’ standardized effect sizes for a trait (from TWAS) or differential expression values for a drug (from pharmacologic perturbation data) are multiplied by the LV genes’ weights and summed, producing a single value. Instead of looking at individual genes, this process links different traits and drugs to functionally-related groups of genes or LVs. PhenoPLIER uses LVs annotations about the specific conditions where the group of genes is expressed, such as cell types and tissues, even at specific developmental stages, disease stages or under distinct stimuli. Although this is not strictly necessary for PhenoPLIER to work, these annotations can dramatically improve the interpretability of results. MultiPLIER’s models provide this information by linking LVs to samples, which may be annotated for experimental conditions (represented by matrix **B** at the top of Figure 1b) in which genes in an LV are expressed. An example of this is shown in Figure 1c. In the original MultiPLIER study, the authors reported that one of the latent variables, identified as LV603, was associated with a known neutrophil pathway and highly correlated with neutrophil count estimates from whole blood RNA-seq profiles [50]. We analyzed LV603 using PhenoPLIER and found that 1) neutrophil counts and other white blood cell traits were ranked among the top 10 traits out of 4,091 (Figure 1c, bottom), and basophils count and percentage were significantly associated with this LV when using our regression method (Supplementary Table 5), and 2) LV603’s genes were expressed in highly relevant cell types (Figure 1c, top). These initial results suggested that groups of functionally related and co-expressed genes tend to correspond to groups of trait-associated genes, and the approach can link transcriptional mechanisms from large and diverse dataset collections to complex traits.

Therefore, PhenoPLIER allows the user to address specific questions, namely: do disease-associated genes belong to modules expressed in specific tissues and cell types? Are these cell type-specific modules associated with *different* diseases, thus potentially representing a “network pleiotropy” example from an omnigenic point of view [31]? Is there a subset of module’s genes that is closer to the definition of “core” genes (i.e., directly affecting the trait with no mediated regulation of other genes [34]) and thus represents alternative and potentially better candidate targets? Are drugs perturbing these transcriptional mechanisms, and can they suggest potential mechanisms of action?

### LVs link genes that alter lipid accumulation with relevant traits and tissues

Our first experiment attempted to answer whether genes in a disease-relevant LV could represent potential therapeutic targets. For this, the first step was to obtain a set of genes strongly associated with a phenotype of interest. Therefore, we performed a fluorescence-based CRISPR-Cas9 in the HepG2 cell line and identified 462 genes associated with lipid regulation (Methods). From these, we selected two high-confidence gene sets that either caused a decrease or increase of lipids: a lipids-decreasing gene-set with eight genes: *BLCAP*, *FBXW7*, *INSIG2*, *PCYT2*, *PTEN*, *SOX9*, *TCF7L2*, *UBE2J2*; and a lipids-increasing gene-set with six genes: *ACACA*, *DGAT2*, *HILPDA*, *MBTPS1*, *SCAP*, *SRPR* (Supplementary File 2).

Next, we analyzed all 987 LVs using Fast Gene Set Enrichment Analysis (FGSEA) [51], and found 15 LVs nominally enriched (unadjusted *P* < 0.01) with these lipid-altering gene-sets (Supplementary Tables 2 and 3). Among those with reliable sample metadata, LV246, the top LV associated with the lipids-increasing gene-set, contained genes mainly co-expressed in adipose tissue (Figure 2a), which plays a key role in coordinating and regulating lipid metabolism. Using our regression framework across all traits in PhenomeXcan, we found that gene weights for this LV were predictive of gene associations for plasma lipids, high cholesterol, and Alzheimer’s disease (Supplementary Table 8, FDR < 1e-23). These lipids-related associations also replicated across the 309 traits in eMERGE (Supplementary Table 9), where LV246 was significantly associated with hypercholesterolemia (phecode: 272.11, FDR < 4e-9), hyperlipidemia (phecode: 272.1, FDR < 4e-7) and disorders of lipoid metabolism (phecode: 272, FDR < 4e-7).

**Figure 2:**
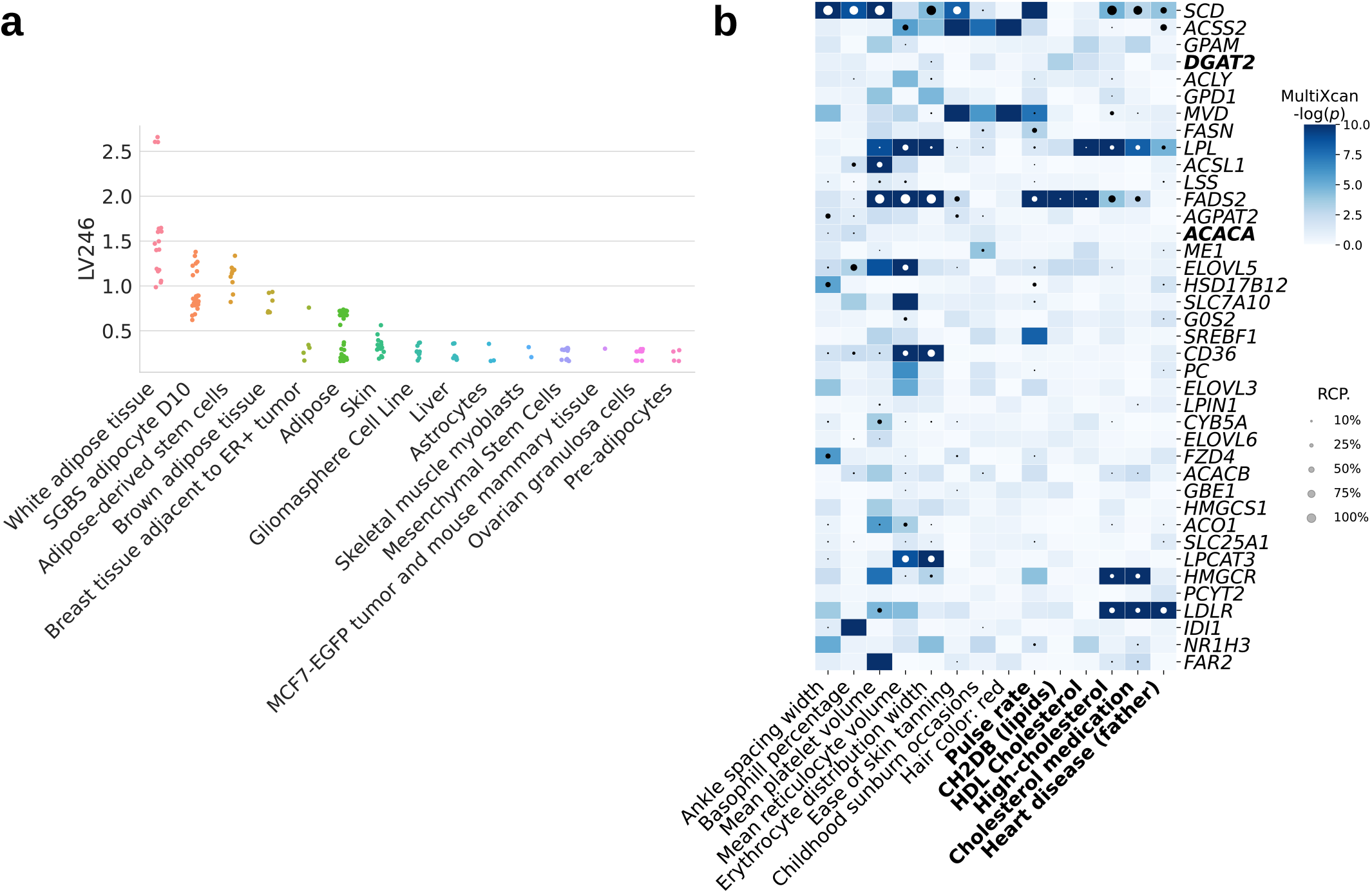
Tissues and traits associated with a gene module related to lipid metabolism (LV246). **a)** Top cell types/tissues in which LV246’s genes are expressed. Values in the *y*-axis come from matrix **B** in the MultiPLIER models (Figure 1b, see Methods). In the ***x***-axis, cell types/tissues are sorted by the maximum sample value. **b)** Gene-trait associations (S-MultiXcan; threshold at -log(*p*=10) and colocalization probability (fastENLOC) for the top traits in LV246. The top 40 genes in LV246 are shown, sorted by their LV weight (matrix **Z**), from largest (the top gene *SCD*) to smallest (*FAR2*); *DGAT2* and *ACACA*, in boldface, are two of the six high-confidence genes in the lipids-increasing gene set from the CRISPR screen. Cardiovascular-related traits are in boldface. SGBS: Simpson Golabi Behmel Syndrome; CH2DB: CH_2_ groups to double bonds ratio; HDL: high-density lipoprotein; RCP: locus regional colocalization probability.

Two high-confidence genes from our CRISPR screening, *DGAT2* and *ACACA*, are responsible for encoding enzymes for triglycerides and fatty acid synthesis and were among the highest-weighted genes of LV246 (Figure 2b, in boldface). However, in contrast to other members of LV246, *DGAT2* and *ACACA* were not associated nor colocalized with any of the cardiovascular-related traits and thus would not have been prioritized by TWAS alone; instead, other members of LV246, such as *SCD*, *LPL*, *FADS2*, *HMGCR*, and *LDLR*, were significantly associated and colocalized with lipid-related traits. This lack of association of two high-confidence genes from our CRISPR screen might be explained from an omnigenic point of view [34]. Assuming that the TWAS models for *DGAT2* and *ACACA* capture all common *cis*-eQTLs (the only genetic component of gene expression that TWAS can capture) and there are no rare *cis*-eQTLs, these two genes might represent “core” genes (i.e., they directly affect the trait with no mediated regulation of other genes), and many of the rest in the LV are “peripheral” genes that *trans*-regulate them.

### LVs predict drug-disease pairs better than single genes

We next determined how substituting LVs for individual genes predicted known treatment-disease relationships. For this, we used the transcriptional responses to small molecule perturbations profiled in LINCS L1000 [43], which were further processed and mapped to DrugBank IDs [52,53,54]. Based on an established drug repurposing strategy that matches reversed transcriptome patterns between genes and drug-induced perturbations [55,56], we adopted a previously described framework that uses imputed transcriptomes from TWAS to prioritize drug candidates [30]. For this, we computed a drug-disease score by calculating the negative dot product between the *z*-scores for a disease (from TWAS) and the *z*-scores for a drug (from LINCS) across sets of genes of different sizes (see Methods). Therefore, a large score for a drug-disease pair indicated that higher (lower) predicted expression values of disease-associated genes are down (up)-regulated by the drug, thus predicting a potential treatment. Similarly, for the LV-based approach, we estimated how pharmacological perturbations affected the gene module activity by projecting expression profiles of drugs into our latent representation (Figure 1b). We used a manually-curated gold standard set of drug-disease medical indications [53,57] for 322 drugs across 53 diseases to evaluate the prediction performance.

It is important to note that the gene-trait associations and drug-induced expression profiles projected into the latent space represent a compressed version of the entire set of results. Despite this information loss, the LV-based method outperformed the gene-based one with an area under the curve of 0.632 and an average precision of 0.858 (Figure 3). The prediction results suggested that this low-dimensional space captures biologically meaningful patterns that can link pathophysiological processes with the mechanism of action of drugs.

**Figure 3:**
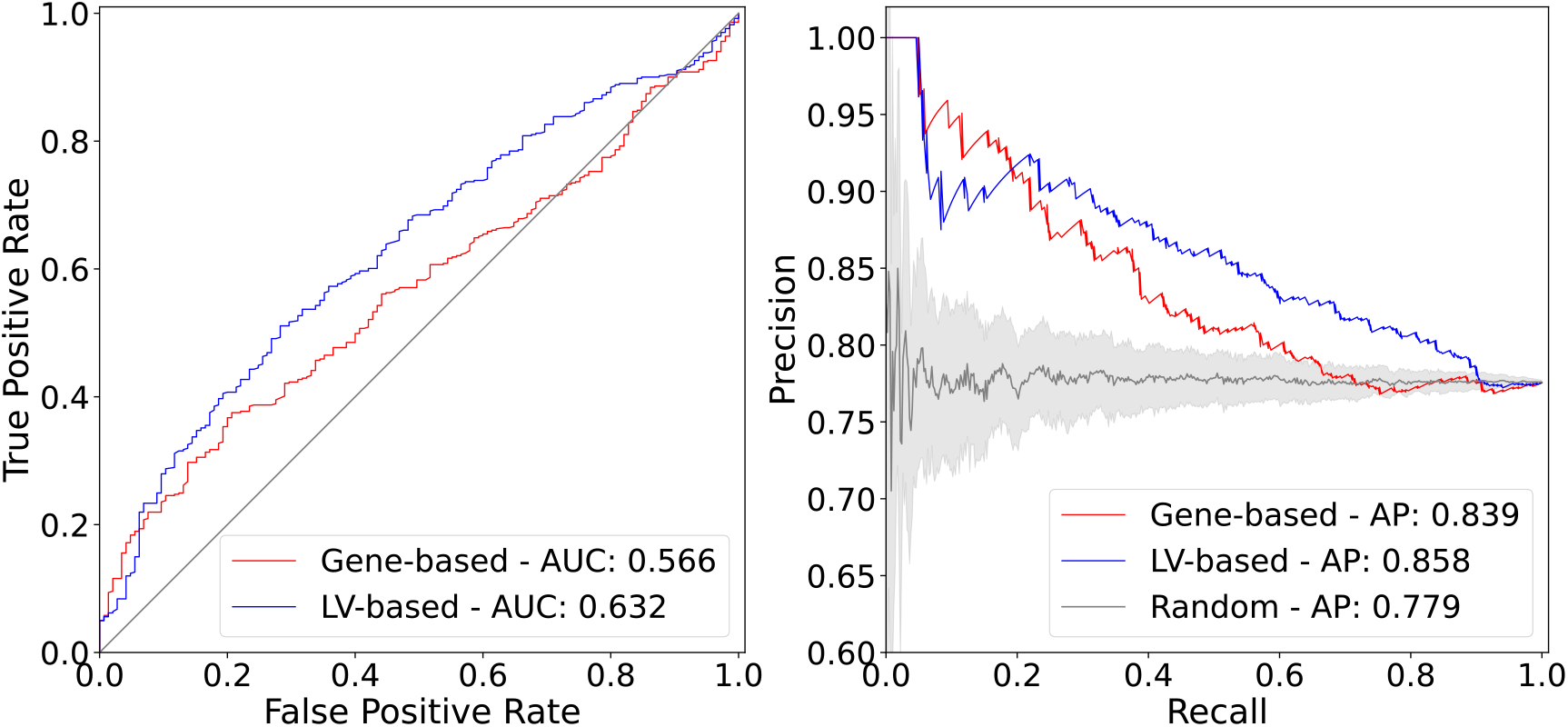
Drug-disease prediction performance for gene-based and LV-based approaches. The receiver operating characteristic (ROC) (left) and the precision-recall curves (right) for a gene-based and LV-based approach. AUC: area under the curve; AP: average precision.

We examined a specific drug-disease pair to determine whether the LVs driving the prediction were biologically plausible. Nicotinic acid (niacin) is a B vitamin widely used clinically to treat lipid disorders, although there is controversy on its clinical utility in preventing cardiovascular disease [58,59,60]. Niacin exerts its effects on multiple tissues, although its mechanisms are not well understood [61,62,63,64]. This compound can increase high-density lipoprotein (HDL) by inhibiting an HDL catabolism receptor in the liver. Niacin also inhibits diacylglycerol acyltransferase–2 (DGAT2), which decreases the production of low-density lipoproteins (LDL) either by modulating triglyceride synthesis in hepatocytes or by inhibiting adipocyte triglyceride lipolysis [61]. Niacin was one of the drugs in the gold standard set indicated for atherosclerosis (AT) and coronary artery disease (CAD). We observed that this compound was predicted by the gene-based and LV-based approach as a medical indication for coronary artery disease (CAD), with scores above the mean (0.51 and 0.96, respectively). For AT, the LV-based approach predicted niacin as a therapeutic drug with a score of 0.52, whereas the gene-based method assigned a negative score of −0.01 (below the mean). Since LVs represent interpretable features associated with specific cell types, we analyzed which LVs positively contributed to these predictions (i.e., with an opposite direction between niacin and the disease). Notably, LV246 (Figure 2), expressed in adipose tissue and liver and associated with plasma lipids and high cholesterol (Supplementary Table 8), was the 16th most important module in the prediction of niacin as a therapeutic drug for AT. Besides the gold standard set, LV246 was among the top modules for other cardiovascular diseases, such as ischaemic heart disease (wide definition, 15th module) and high cholesterol (7th module).

The analysis of other top niacin-contributing LVs across different cardiovascular diseases revealed additional mechanisms of action. For example, *GPR109A/HCAR2* encodes a G protein-coupled high-affinity niacin receptor in adipocytes and immune cells, including monocytes, macrophages, neutrophils and dendritic cells [65,66]. It was initially thought that the antiatherogenic effects of niacin were solely due to the inhibition of lipolysis in adipose tissue. However, it has been shown that nicotinic acid can reduce atherosclerosis progression independently of its antidyslipidemic activity by activating *GPR109A* in immune cells [67], thus boosting anti-inflammatory processes [68]. In addition, flushing, a common adverse effect of niacin, is also produced by the activation of GPR109A in Langerhans cells (macrophages of the skin). This alternative mechanism for niacin could have been hypothesized by examining the cell types where the top-contributing modules are expressed: for instance, LV116 and LV931 (Figure 4, Supplementary Figure 17, and Supplementary Tables 10 and 11) were the top two modules for AT, with a strong signature in monocytes, macrophages, neutrophils, dendritic cells, among others. In Figure 4, it can be seen that LV116’s genes are expressed as an immune response when these cell types are under different stimuli, such as diarrhea caused by different pathogens [69], samples from multiple sclerosis or systemic lupus erythematosus [70,71], or infected with different viruses (such as herpes simplex [72], West Nile virus [73], *Salmonella typhimurium* [74], among others). These three LVs (LV246, LV116 and LV931) were among the top 20 modules contributing to the niacin prediction across different cardiovascular traits (Table 1).

**Figure 4:**
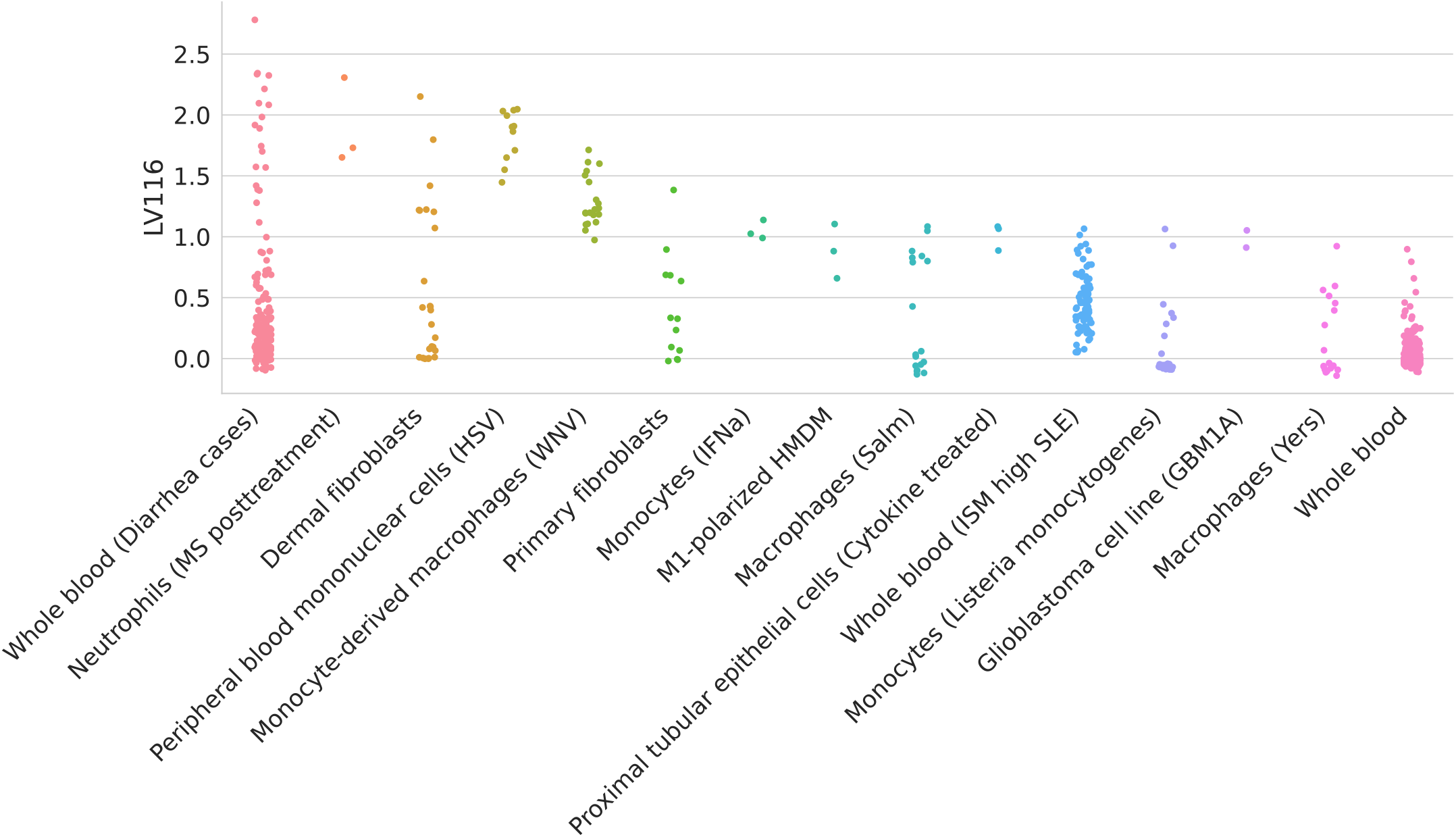
Top cell types/tissues where LV116 s genes are expressed. Values in the ***y*** axis come from matrix **B** in the MultiPLIER models (Figure 1b). In the ***x***-axis, cell types/tissues are sorted by the maximum sample value. The figure shows a clear immune response with cell types under different stimuli. MS: multiple sclerosis; HSV: treated with herpes simplex virus; WNV: infected with West Nile virus; IFNa: treated with interferon-alpha; HMDM: human peripheral blood mononuclear cell-derived macrophages; Salm: infected with *Salmonella typhimurium*; Yers: infected with *Yersinia pseudotuberculosis*; ISM: Interferon Signature Metric; SLE: Systemic lupus erythematosus.

**Table 1:**
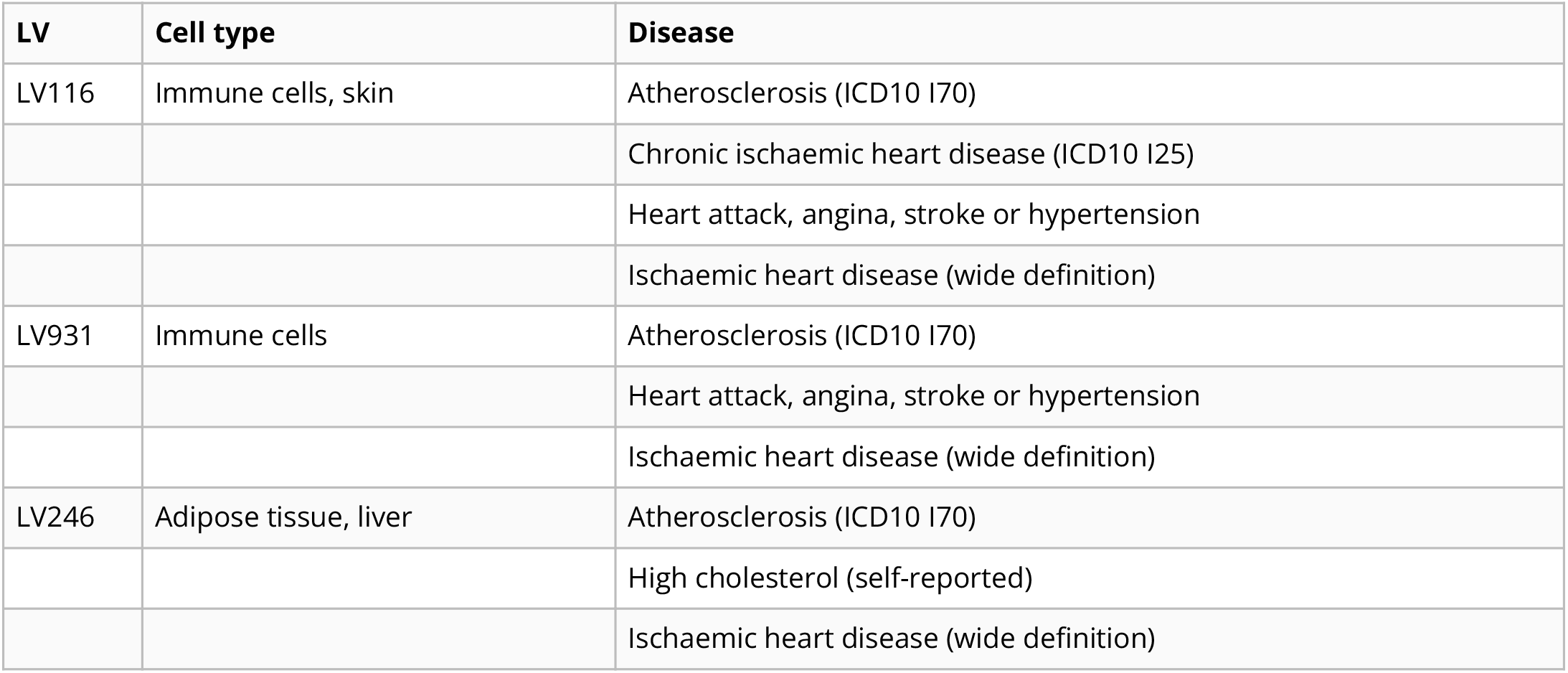
LVs among the top 20 contributors to the prediction of niacin for five cardiovascular diseases. “Heart attack, angina, stroke or hypertension” refers to the UK Biobank data-field 6150. GWAS sample size: Atherosclerosis (361,194 in total and 566 cases), Chronic ischaemic heart disease (361,194 in total and 12,769 cases), Heart attack, angina, stroke or hypertension (360,420 in total and 253,565 cases), Ischaemic heart disease/wide definition (361,194 in total and 20,857 cases), High cholesterol/self-reported (361,141 in total and 43,957 cases).

**Table 2:** Gene modules (LVs) nominally enriched for the lipids-increasing gene-set from the CRISPR-screen (*P* < 0.01). LVs significantly aligned with pathways (FDR < 0.05) from the MultiPLIER models are shown in boldface.

| Gene module | Lipids gene-set | Leading edge | p-value |
| --- | --- | --- | --- |
| **LV246** | increase | *DGAT2, ACACA* | 0.0035 |
| LV702 | increase | *ACACA, DGAT2* | 0.0046 |
| **LV607** | increase | *ACACA, DGAT2* | 0.0058 |
| LV890 | increase | *ACACA, DGAT2* | 0.0067 |
| **LV74** | increase | *MBTPS1, DGAT2* | 0.0078 |
| **LV865** | increase | *ACACA, DGAT2* | 0.0092 |
| LV841 | increase | *ACACA, DGAT2* | 0.0096 |

**Table 3:** Gene modules (LVs) nominally enriched for the lipids-decreasing gene-set from the CRISPR-screen (*P* < 0.01). LVs significantly aligned with pathways (FDR < 0.05) from the MultiPLIER models are shown in boldface.

| Gene module | Lipids gene-set | Leading edge | p-value |
| --- | --- | --- | --- |
| LV520 | decrease | *FBXW7, TCF7L2* | 0.0006 |
| LV801 | decrease | *UBE2J2, TCF7L2* | 0.0022 |
| LV512 | decrease | *FBXW7, TCF7L2* | 0.0025 |
| **LV612** | decrease | *PTEN, FBXW7* | 0.0036 |
| LV41 | decrease | *PCYT2, TCF7L2* | 0.0041 |
| **LV838** | decrease | *UBE2J2, TCF7L2* | 0.0070 |
| LV302 | decrease | *TCF7L2, PTEN* | 0.0083 |
| LV959 | decrease | *TCF7L2, PTEN* | 0.0092 |

**Table 4:** Pathways aligned to LV603 from the MultiPLIER models.

| Pathway | AUC | FDR |
| --- | --- | --- |
| IRIS Neutrophil-Resting | 0.91 | 4.51e-35 |
| SVM Neutrophils | 0.98 | 1.43e-09 |
| PID IL8CXCR2 PATHWAY | 0.81 | 7.04e-03 |
| SIG PIP3 SIGNALING IN B LYMPHOCYTES | 0.77 | 1.95e-02 |

**Table 5:** Significant trait associations of LV603 in PhenomeXcan.

| Trait description | Sample size | Cases | FDR |
| --- | --- | --- | --- |
| Basophill percentage | 349,861 |  | 1.19e-10 |
| Basophill count | 349,856 |  | 1.89e-05 |
| Treatment/medication code: ispaghula husk | 361,141 | 327 | 1.36e-02 |

**Table 6:**
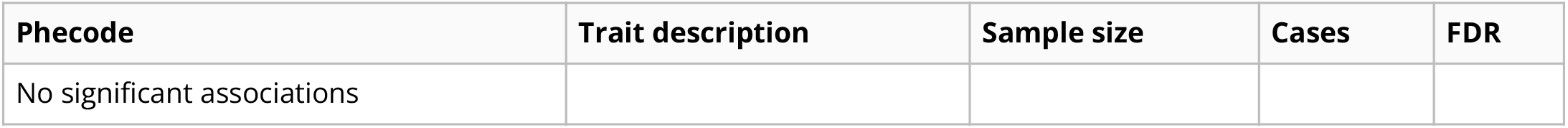
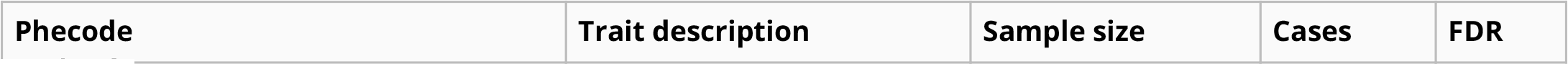
Significant trait associations of LV603 in eMERGE.

**Table 7:**
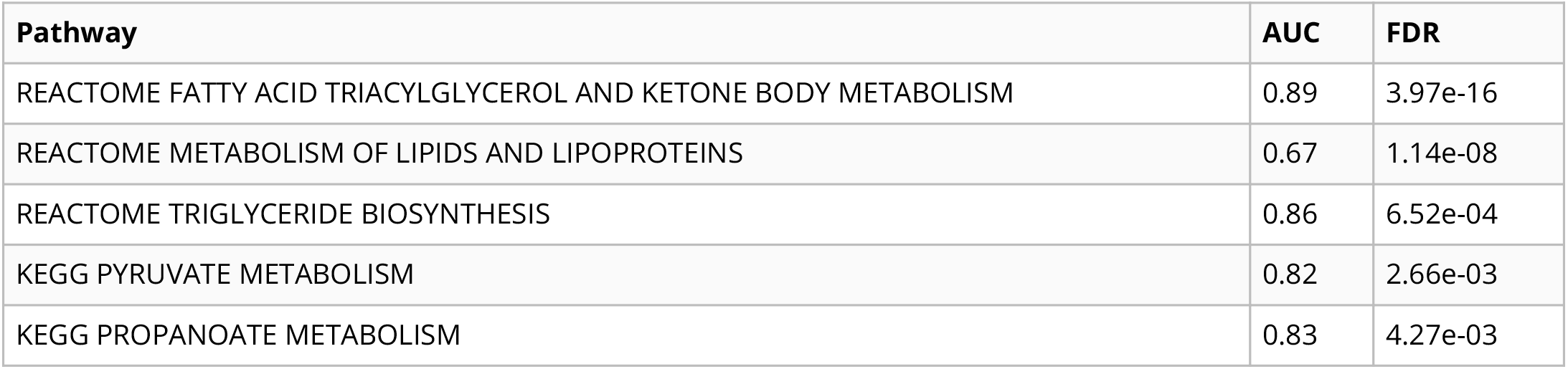
Pathways aligned to LV246 from the MultiPLIER models.

| Pathway | AUC | FDR |
| --- | --- | --- |
| REACTOME FATTY ACID TRIACYLGLYCEROL AND KETONE BODY METABOLISM | 0.89 | 3.97e-16 |
| REACTOME METABOLISM OF LIPIDS AND LIPOPROTEINS | 0.67 | 1.14e-08 |
| REACTOME TRIGLYCERIDE BIOSYNTHESIS | 0.86 | 6.52e-04 |
| KEGG PYRUVATE METABOLISM | 0.82 | 2.66e-03 |
| KEGG PROPANOATE METABOLISM | 0.83 | 4.27e-03 |

Beyond cardiovascular traits, there are other potentially interesting LVs that could extend our understanding of the mechanisms of niacin. For example, LV66, one of the top LVs affected by niacin (Supplementary Figure 18), was mainly expressed in ovarian granulosa cells. This compound has been very recently considered a potential therapeutic for ovarian diseases [75,76], as it was found to promote follicle growth and inhibit granulosa cell apoptosis in animal models.

### LVs reveal trait clusters with shared transcriptomic properties

We used the projection of gene-trait associations into the latent space to find groups of clusters linked by the same transcriptional processes. Since individual clustering algorithms have different biases (i.e., assumptions about the data structure), we designed a consensus clustering framework that combines solutions or partitions of traits generated by different methods (Methods). Consensus or ensemble approaches have been recommended to avoid several pitfalls when performing cluster analysis on biological data [77]. Since diversity in the ensemble is crucial for these methods, we generated different data versions which were processed using different methods with varying sets of parameters (Figure 5a). Then, a consensus function combines the ensemble into a consolidated solution, which has been shown to outperform any individual member of the ensemble [78,79]. Our clustering pipeline generated 15 final consensus clustering solutions (Supplementary Figure 15). The number of clusters of these partitions (between 5 to 29) was learned from the data by selecting the partitions with the largest agreement with the ensemble [78]. Instead of selecting one of these final solutions with a specific number of clusters, we used a clustering tree [80] (Figure 6) to examine stable groups of traits across multiple resolutions. To understand which latent variables differentiated the group of traits, we trained a decision tree classifier on the input data 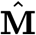 using the clusters found as labels (Figure 5b, see Methods).

**Figure 5:**
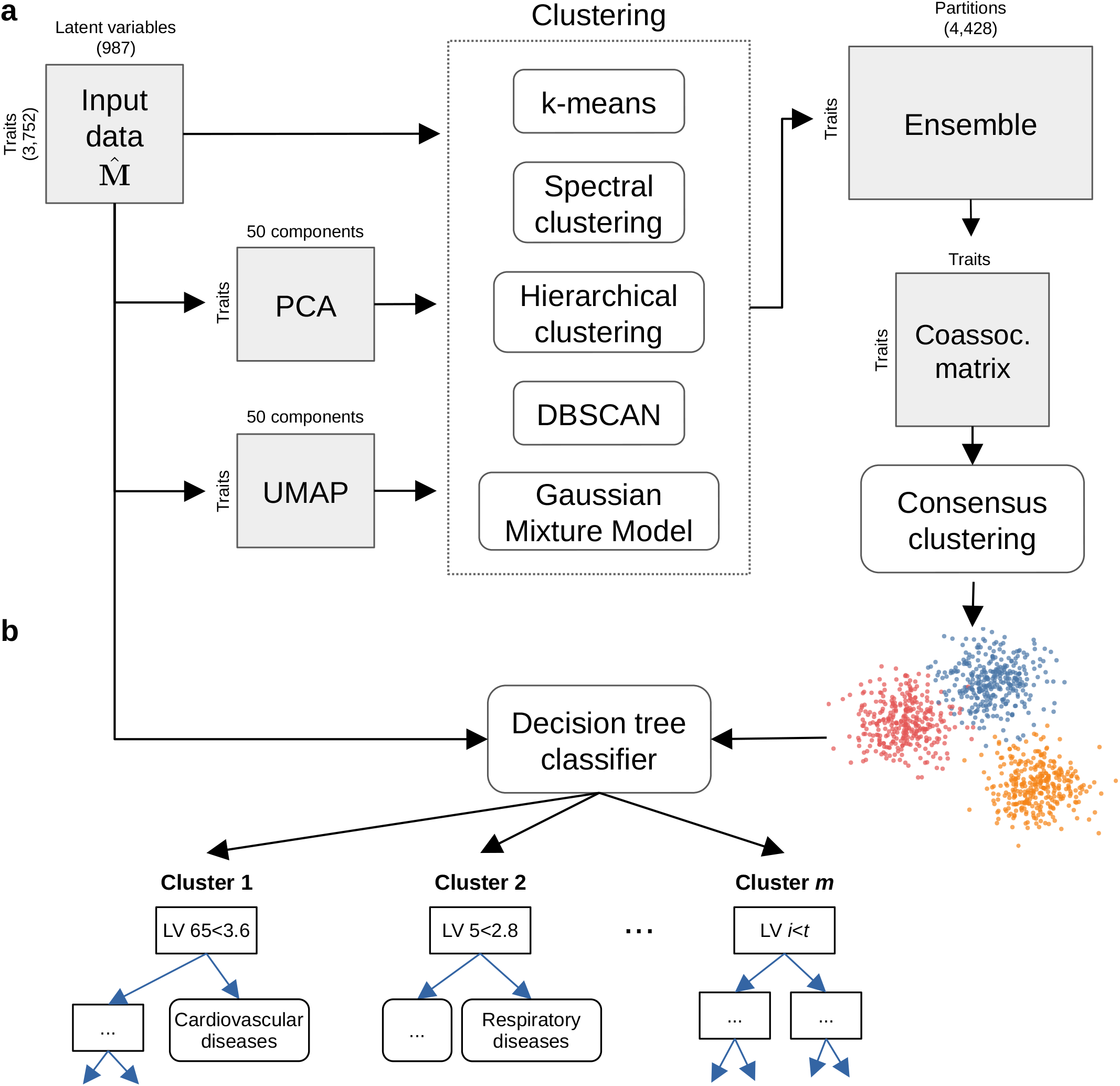
Cluster analysis on traits using the latent gene expression representation. **a)** The projection of TWAS results on 3,752 traits into the latent gene expression representation is the input data to the clustering process. A linear (PCA) and non-linear (UMAP) dimensionality reduction techniques were applied to the input data, and five different clustering algorithms processed all data versions. These algorithms derive partitions from the data using different parameters (such as the number of clusters), leading to an ensemble of 4,428 partitions. Then, a distance matrix is derived by counting how many times a pair of traits was grouped in different clusters across the ensemble. Finally, a consensus function is applied to the distance matrix to generate consolidated partitions with different numbers of clusters (from 2 to 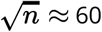). These final solutions were represented in the clustering tree (Figure 6). **b)** The clusters found by the consensus function were used as labels to train a decision tree classifier on the original input data, which detects the LVs that better differentiate groups of traits.

**Figure 6:**
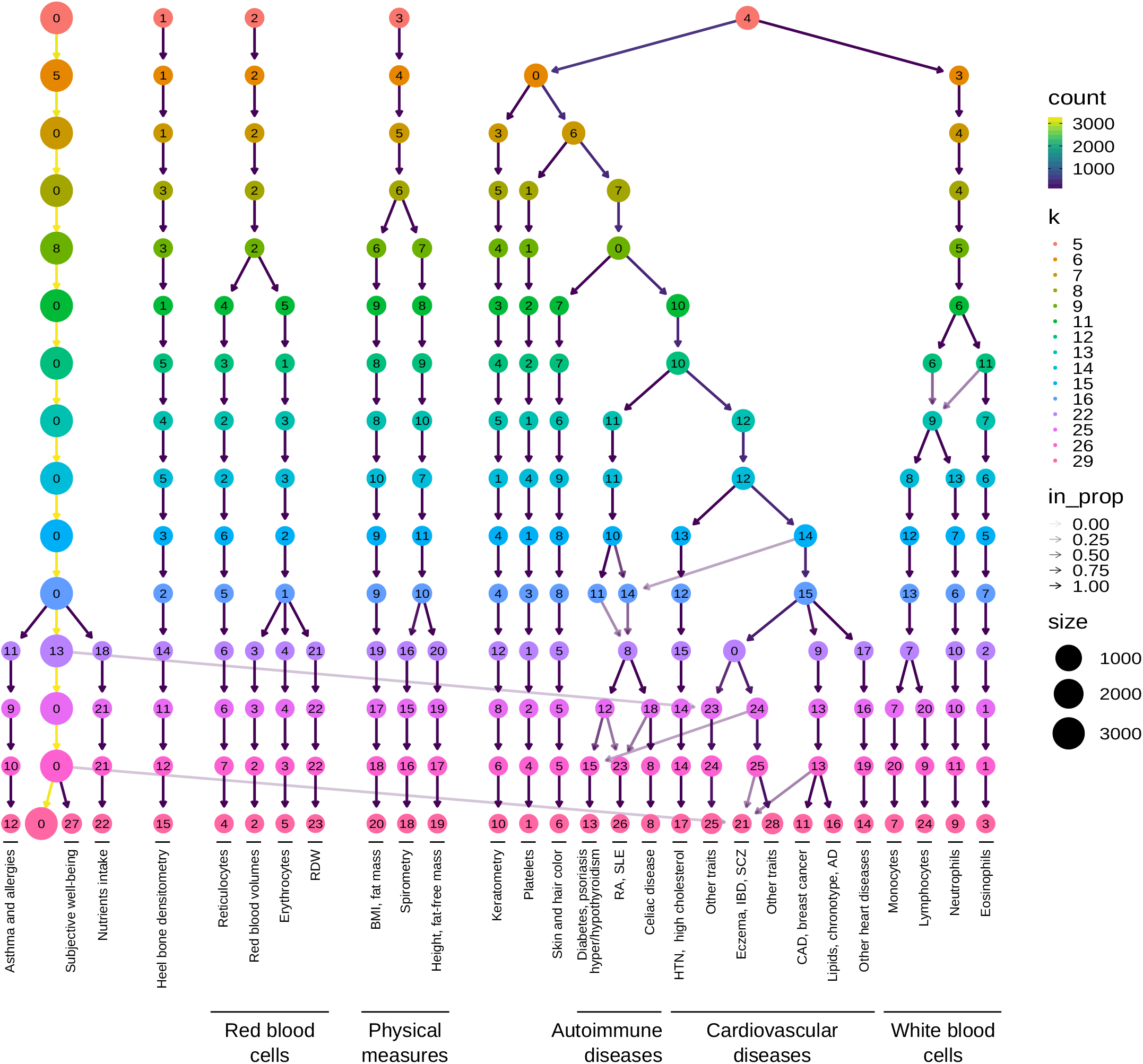
Clustering tree using multiple resolutions for clusters of traits. Each row represents a partition/grouping of the traits, and each circle is a cluster from that partition. The number of clusters goes from 5 to 29. Arrows indicate how traits in one cluster move across clusters from different partitions. Most of the clusters are preserved across different resolutions, showing highly stable solutions even with independent runs of the clustering algorithm. RDW: red cell (erythrocyte) distribution width; BMI: body mass index; WC: waist circumference; HC: hip circumference; RA: rheumatoid arthritis; SLE: systemic lupus erythematosus; HTN: Hypertension; IBD: inflammatory bowel disease; SCZ: Schizophrenia; CAD: Coronary artery disease; AD: Alzheimer’s disease; *Descriptions of traits by cluster ID (from left to right):* 12: also includes lymphocyte count and allergies such as allergic rhinitis or eczema; 4: includes reticulocyte count and percentage, immature reticulocyte fraction, and high light scatter reticulocytes count and percentage; 2: includes mean corpuscular volume, mean corpuscular hemoglobin, mean reticulocyte volume, mean sphered cell volume; 5: includes erythrocyte count, hemoglobin concentration, and hematocrit percentage; 20: also includes weight, waist and hip circumference; 18: also includes body impedance measures and ankle spacing width; 19: also includes basal metabolic rate; 1: includes platelet count, crit, mean volume, and distribution width; 13: diabetes refers to age when diabetes was first diagnosed; 25: also includes vascular problems such as angina, deep vein thrombosis (DVT), intraocular pressure, eye and mouth problems, pulse rate, hand-grip strength, several measurements of physical activity, jobs involving heavy physical work, types of transport used, intake of vitamin/mineral supplements, and various types of body pain and medications for pain relief; 21: also includes attention deficit hyperactivity disorder (ADHD), number of years of schooling completed, bone density, and intracranial volume measurement; 28: includes diabetes, gout, arthrosis, and respiratory diseases (and related medications such as ramipril, allopurinol, and lisinopril), urine assays, female-specific factors (age at menarche, menopause, first/last live birth), and several environmental/behavioral factors such as intake of a range of food/drink items including alcohol, time spent outdoors and watching TV, smoking and sleeping habits, early-life factors (breastfed as a baby, maternal smoking around birth), education attainment, psychological and mental health, and health satisfaction; 11: also includes fasting blood glucose and insulin measurement; 16: lipids include high and low-density lipoprotein (HDL and LDL) cholesterol, triglycerides, and average number of methylene groups per a double bond; 14: includes myocardial infarction, coronary atherosclerosis, ischaemic heart disease (wide definition); 7: includes monocyte count and percentage; 24: includes lymphocyte count and percentage; 9: includes neutrophil count, neutrophil+basophil count, neutrophil+eosinophil count, granulocyte count, leukocyte count, and myeloid cell count; 3: includes eosinophil count, eosinophil percentage, and eosinophil+basophil count.

We found that phenotypes were grouped into five clear branches, defined by their first node at the top of the Figure 6: 0) a “large” branch that includes most of the traits subdivided only starting at ***k***=16 (with asthma, subjective well-being traits, and nutrient intake clusters), 1) heel bone-densitometry measurements, 2) hematological assays on red blood cells, 3) physical measures, including spirometry and body impedance, and anthropometric traits with fat-free and fat mass measures in separate sub-branches, and 4) a “complex” branch including keratometry measurements, assays on white blood cells and platelets, skin and hair color traits, autoimmune disorders, and cardiovascular diseases (which also included other cardiovascular-related traits such as hand-grip strength [81], and environmental/behavioral factors such as physical activity and diet) (see Supplementary Files 3-6 for clustering results). Within these branches, results were relatively stable, with the same traits often clustered together across different resolutions. Arrows between clusters show traits moving from one group to another, and this mainly happens between clusters within the “complex” branch (4) and between clusters from the “large” branch (0) to the “complex” branch. This behavior is expected since complex diseases are usually associated with shared genetic and environmental factors and are thus hard to categorize into a single cluster.

Next, we analyzed which LVs were driving these clusters of traits. For this, we trained decision tree classifiers on the input data using each cluster at ***k***=29 (bottom of Figure 6) as labels (see Methods). This procedure yielded the top LVs that were most discriminative for each cluster. Several of these LVs were well-aligned to existing pathways (Figure 7), whereas others were not aligned to prior knowledge but still expressed in relevant tissues (Supplementary Figure 16). In Figure 7, it can be seen that some LVs were highly specific to certain traits, while others were associated with a wide range of different phenotypes, thus potentially involved in more general biological functions. We used our regression framework to determine whether these LVs were significantly associated with different traits. For example, LVs such as LV928 and LV30, which were well-aligned to early progenitors of the erythrocytes lineage [82] (Supplementary Tables 13 and 16), were predominantly expressed in early differentiation stages of erythropoiesis (Supplementary Figures 19 and 20) and strongly associated with different assays on red blood cells (FDR < 0.05; Supplementary Tables 14, 15, and 18). In contrast, other LVs were highly specific, such as LV730, which is expressed in thrombocytes from different cancer samples (Supplementary Figures 21 and Supplementary Table 19), and strongly associated with hematological assays on platelets (FDR < 0.05, Supplementary Table 20); or LV598, whose genes were expressed in corneal endothelial cells (Supplementary Figures 22 and Supplementary Table 22) and associated with keratometry measurements (Supplementary Table 23).

**Figure 7:**
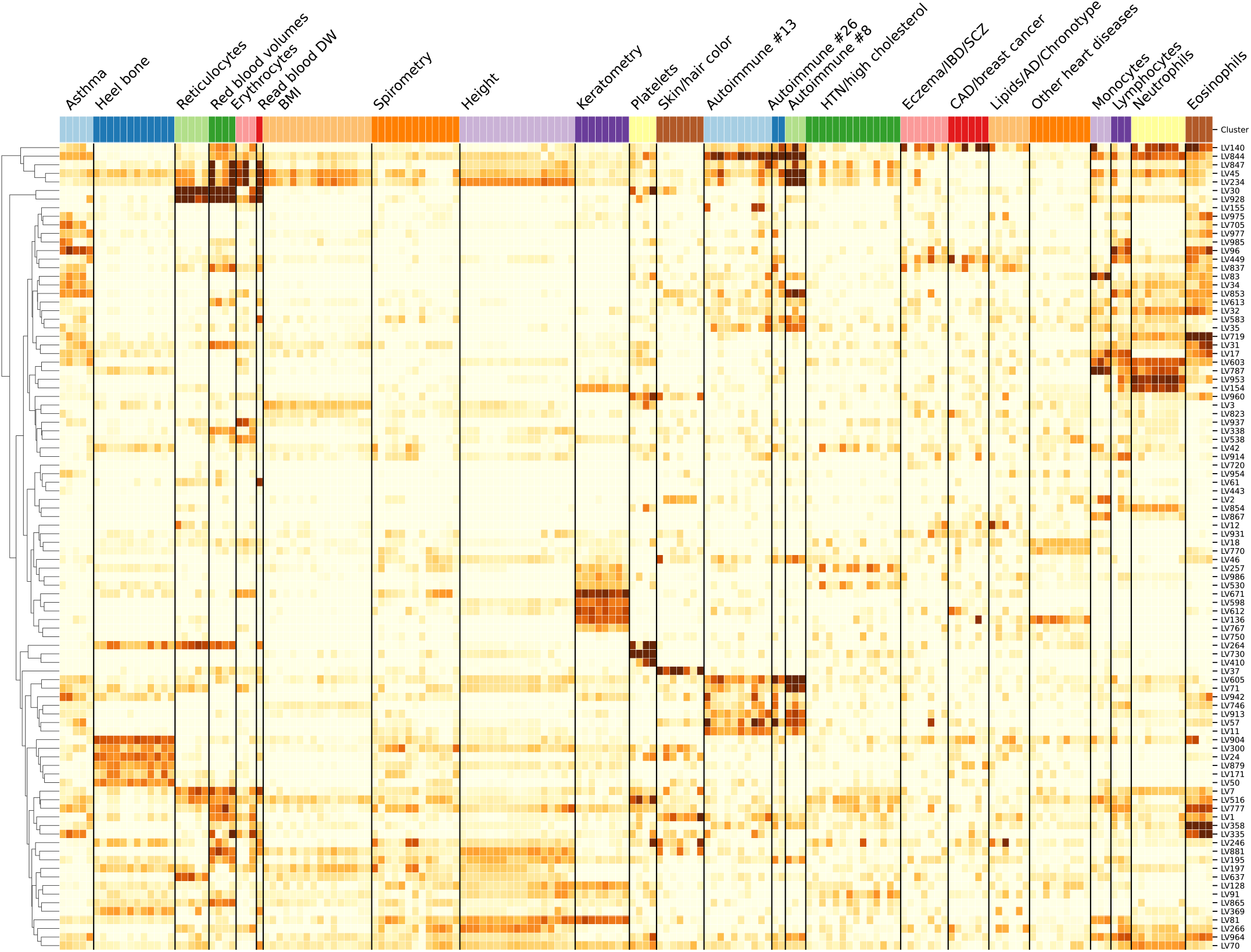
Cluster-specific and general transcriptional processes associated with different diseases. The plot shows a submatrix of 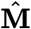 for the main trait clusters at ***k***=29, considering only LVs (rows) that are well-aligned with at least one pathway.

The sub-branches of autoimmune and cardiovascular diseases merged together at ***k*** = 10 (middle of Figure 6), so we expected to find LVs that specifically affect one or both of these types of diseases. For example, LV57, expressed in T cells (Supplementary Figure 23 and Supplementary Table 25), was the most strongly associated gene module with autoimmune disorders in PhenomeXcan (Supplementary Table 26), with significant associations with hypothyroidism that were replicated in eMERGE (27). However, this LV was also strongly associated with deep venous thrombosis in both PhenomeXcan and eMERGE. On the other hand, LV844 was more autoimmune-specific, with associations to polymyalgia rheumatica, type 1 diabetes, rheumatoid arthritis, and celiac disease in PhenomeXcan (Supplementary Table 29). However, these did not replicate in eMERGE. This LV was expressed in a wide range of cell types, including blood, breast organoids, myeloma cells, lung fibroblasts, and different cell types from the brain (Supplementary Figure 24 and Supplementary Table 28).

The cardiovascular sub-branch had 129 significant LV-trait associations in PhenomeXcan and 23 in eMERGE. LV136, aligned with known collagen formation and muscle contraction pathways (Supplementary Table 31), was associated with coronary artery disease and keratometry measurements in PhenomeXcan (Supplementary Tables 32). In eMERGE, this LV was associated with coronary atherosclerosis (phecode: 411.4) (Supplementary Table 33). LV136 was expressed in a wide range of cell types, including fibroblasts, mesenchymal stem cells, osteoblasts, pancreatic stellate cells, cardiomyocytes, and adipocytes (Supplementary Figure 25). Within the cardiovascular sub-branch, we found neuropsychiatric and neurodevelopmental disorders such as Alzheimer’s disease, schizophrenia, and attention deficit hyperactivity disorder (ADHD). These disorders were previously linked to the cardiovascular system [83,84,85,86] and share several risk factors, including hypertension, high cholesterol, obesity, smoking, among others [87,88]. However, our results grouped these diseases by potentially shared transcriptional processes expressed in specific tissues/cell types. Alzheimer’s disease (not present in eMERGE), for instance, was significantly associated with LV21 in PhenomeXcan (Supplementary Table 35). LV21, a gene module not aligned to prior pathways, was strongly expressed in a variety of soft tissue sarcomas, monocytes/macrophages (including microglia from cortex samples), and aortic valves (Supplementary Figure 26 and Supplementary Table 34). This LV was also strongly associated with lipids and high cholesterol in PhenomeXcan and hyperlipidemia (phecode: 272.1) in eMERGE (Supplementary Table 36). As discussed previously, macrophages play a key role in the reverse cholesterol transport and thus atherogenesis [89], and lipid metabolism in microglia has been recently identified as an important factor in the development of neurodegenerative diseases [90].

## Discussion

We have introduced a novel computational strategy that integrates statistical associations from TWAS with groups of genes (gene modules) that have similar expression patterns across the same cell types. Our key innovation is that we project gene-trait associations through a latent representation derived not strictly from measures of normal tissue but also from cell types under a variety of stimuli and at various developmental stages. This improves interpretation by going beyond statistical associations to infer cell type-specific features of complex phenotypes. Our approach can identify disease-relevant cell types from summary statistics, and several disease-associated gene modules were replicated in eMERGE. Using a CRISPR screen to analyze lipid regulation, we found that our gene module-based approach can prioritize causal genes even when single gene associations are not detected. We interpret these findings with an omnigenic perspective of “core” and “peripheral” genes, suggesting that the approach can identify genes that directly affect the trait with no mediated regulation of other genes and thus prioritize alternative and potentially more attractive therapeutic targets.

Using our gene module perspective, we also integrated drug-induced transcriptional profiles, which allowed us to connect diseases, drugs, and cell types. We showed that the LV-based drug-repurposing approach outperformed the gene-based one when predicting drug-disease links for 322 drugs across 53 diseases. Furthermore, and beyond statistical prediction, we focused on cardiovascular traits and a particular drug, niacin, to show that the approach connects pathophysiological processes with known mechanisms of action, including those in adipose tissue, immune cells, and ovarian granulosa cells.

Our LV-based approach could be helpful in generating novel hypotheses to evaluate potential mechanisms of action, or even adverse effects, of known or experimental drugs.

We found that the analysis of associations through latent representations provided reasonable groupings of diseases and traits affected by shared and distinct transcriptional mechanisms expressed in highly relevant tissues. Our cluster analysis approach also detected the LVs that were most discriminative for each cluster. Several of these LVs were also significantly associated with different traits. Some LVs were strongly aligned with known pathways, but others (like LV57) were not, which might represent novel disease-relevant mechanisms. In some cases, the features/LVs linked to phenotypes appear to be associated with specific cell types. Associations with such cell type marker genes may reveal potentially causal cell types for a phenotype with more precision. We observed modules expressed primarily in one tissue (such as adipose in LV246 or ovary in LV66). Others appeared to be expressed in many contexts, which may capture pathways associated with related complex diseases. For example, LV136 is associated with cardiovascular disease and measures of corneal biomechanics and is expressed in fibroblasts, osteoblasts, pancreas, liver, and cardiomyocytes, among others. Other examples include LV844, expressed in whole blood samples and associated with a range of autoimmune diseases; or LV57, which is clearly expressed in T cells and strongly associated with autoimmune and venous thromboembolism. From an omnigenic point of view, these patterns might represent cases of “network pleiotropy,” where the same cell types mediate molecularly related traits. To our knowledge, projection through a representation learned on complementary but distinct datasets is a novel approach to identifying cell type and pathway effects on complex phenotypes that is computationally simple to implement.

We also demonstrated that clustering trees, introduced initially as a means to examine developmental processes in single-cell data, provide a multi-resolution grouping of phenotypes based on latent variable associations. We employed hard-partitioning algorithms (one trait belongs exclusively to one cluster) where the distance between two traits takes into account all gene modules. However, it is also plausible for two complex diseases to share only a few biological processes instead of being similar across most of them. Another important consideration is that our TWAS results were derived from a large set of GWAS of different sample sizes and qualities. Although the potential issues derived from this data heterogeneity were addressed before performing our cluster analyses on traits, data preprocessing steps are always challenging and might not avoid bias altogether. Considering groups of related diseases was previously shown to be more powerful in detecting shared genetic etiology [91,92], and clustering trees provide a way to explore such relationships in the context of latent variables.

Finally, we developed an LV-based regression framework to detect whether gene modules are associated with a trait using TWAS ***p***-values. We used PhenomeXcan as a discovery cohort across four thousand traits, and many LV-trait associations replicated in eMERGE. In PhenomeXcan, we found 3,450 significant LV-trait associations (FDR < 0.05) with 686 LVs (out of 987) associated with at least one trait and 1,176 traits associated with at least one LV. In eMERGE, we found 196 significant LV-trait associations, with 116 LVs associated with at least one trait/phecode and 81 traits with at least one LV. We only focused on a few disease types from our trait clusters, but the complete set of associations on other disease domains is available in our Github repository for future research. As noted in Methods, one limitation of the regression approach is that the gene-gene correlations are only approximately accurate, which could lead to false positives if the correlation among the top genes in a module is not precisely captured. The regression model, however, is approximately well-calibrated, and we did not observe inflation when running the method in real data.

Our approach rests on the assumption that gene modules with coordinated expression patterns will also manifest coordinated pathological effects. Our implementation in this work integrates two complementary approaches. The first is MultiPLIER, which extracts latent variables from large expression datasets, and these LVs could represent either real transcriptional processes or technical factors (“batch effects”). We used a previously published model derived from recount2, which was designed to analyze rare disorders but might not be the optimal latent representation for the wide range of complex diseases considered here. Also, the underlying factorization method rests on linear combinations of variables, which could miss important and more complex co-expression patterns. In addition, recount2, the training dataset used, has since been surpassed in size and scale by other resources [20,93]. However, it is important to note that our models impose very few assumptions on the latent expression representation. Therefore, we should be able to easily replace MultiPLIER with other similar approaches like GenomicSuperSignature [94]. The second approach we used in this study is TWAS, where we are only considering the hypothesis that GWAS loci affect traits via changes in gene expression. Other effects, such as coding variants disrupting protein-protein interactions, are not captured. Additionally, TWAS has several limitations that can lead to false positives [95,96]. Like GWAS, which generally detects groups of associated variants in linkage disequilibrium (LD), TWAS usually identifies several genes within the same locus [25,97]. This is due to sharing of GWAS variants in gene expression models, correlated expression of nearby genes, or even correlation of their predicted expression due to eQTLs in LD, among others [95]. Our LV-based regression framework, however, accounts for these gene-gene correlations in TWAS reasonably well.

Our findings are concordant with previous studies showing that drugs with genetic support are more likely to succeed through the drug development pipeline [7,30]. In this case, projecting association results through latent variables better prioritized disease-treatment pairs than considering single-gene effects alone. An additional benefit is that the latent variables driving predictions represent interpretable genetic features that can be examined to infer potential mechanisms of action. Here we prioritized drugs for diseases with very different tissue etiologies, and a challenge of the approach is to select the most appropriate tissue model from TWAS to find reversed transcriptome patterns between genes and drug-induced perturbations.

Ultimately, the quality of the representations is essential to performance. Here we used a representation derived from a factorization of bulk RNA-seq data. Detailed perturbation datasets and single-cell profiling of tissues, with and without perturbagens, and at various stages of development provide an avenue to generate higher quality and more interpretable representations. On the other hand, the key to interpretability is driven by the annotation of sample metadata. New approaches to infer and annotate with structured metadata are promising and can be directly applied to existing data [98]. Rapid improvements in both areas set the stage for latent variable projections to be widely applied to disentangle the genetic basis of complex human phenotypes. By providing a new perspective for a mechanistic understanding of statistical associations from TWAS, our method can generate testable hypotheses for the post-GWAS functional characterization of complex diseases, which will likely be an area of great importance in the coming years.

## Methods and materials

PhenoPLIER is a framework that combines different computational approaches to integrate gene-trait associations and drug-induced transcriptional responses with groups of functionally-related genes (referred to as gene modules or latent variables/LVs). Gene-trait associations are computed using the PrediXcan family of methods, whereas latent variables are inferred by the MultiPLIER models applied on large gene expression compendia. PhenoPLIER provides 1) a regression model to compute an LV-trait association, 2) a consensus clustering approach applied to the latent space to learn shared and distinct transcriptomic properties between traits, and 3) an interpretable, LV-based drug repurposing framework. We provide the details of these methods below.

### The PrediXcan family of methods for gene-based associations

We used Summary-PrediXcan (S-PrediXcan) [99] and Summary-MultiXcan (S-MultiXcan) [24] as the gene-based statistical approaches, which belong to the PrediXcan family of methods [25]. We broadly refer to these approaches as TWAS (transcription-wide association studies). S-PrediXcan, the summary-based version of PrediXcan, computes the univariate association between a trait and a gene’s predicted expression in a single tissue. In contrast, S-MultiXcan, the summary-based version of MultiXcan, computes the joint association between a gene’s predicted expression in all tissues and a trait. S-PrediXcan and S-MultiXcan only need GWAS summary statistics instead of individual-level genotype and phenotype data.

Here we briefly provide the details about these TWAS methods that are necessary to explain our regression framework later (see the referenced articles for more information). In the following, we refer to **y** as a vector of traits for *n* individuals that is centered for convenience (so that no intercept is necessary); 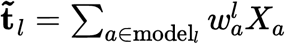 is the gene’s predicted expression for all individuals in tissue ***l*, *X_a_*** is the genotype of SNP *a* and its *w_a_* weight in the tissue prediction model ***l***; and **t_*l*_** is the standardized version of 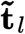 with mean equal to zero and standard deviation equal to one.

S-PrediXcan [99] is the summary version of PrediXcan [25]. PrediXcan models the trait as a linear function of the gene’s expression on a single tissue using the univariate model

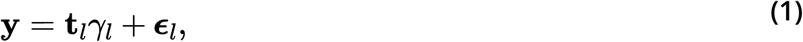

where 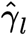 is the estimated effect size or regression coefficient, and ***ϵ_l_*** are the error terms with variance 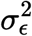. The significance of the association is assessed by computing the *z*-score 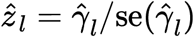 for a gene’s tissue model ***l***. PrediXcan needs individual-level data to fit this model, whereas S-PrediXcan approximates PrediXcan *z*-scores using only GWAS summary statistics with the expression

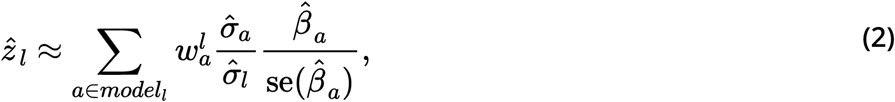

where 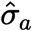 is the variance of SNP ***a***, 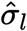 is the variance of the predicted expression of a gene in tissue ***l***, and 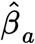 is the estimated effect size of SNP ***a*** from the GWAS. In these TWAS methods, the genotype variances and covariances are always estimated using the Genotype-Tissue Expression project (GTEx v8) [4] as the reference panel. Since S-PrediXcan provides tissue-specific direction of effects (for instance, whether a higher or lower predicted expression of a gene confers more or less disease risk), we used the *z*-scores in our drug repurposing approach (described below).

S-MultiXcan [24], on the other hand, is the summary version of MultiXcan. MultiXcan is more powerful than PrediXcan in detecting gene-trait associations, although it does not provide the direction of effects. Its main output is the ***p***-value (obtained with an F-test) of the multiple tissue model

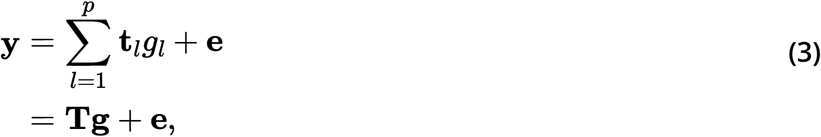

where **T** is a matrix with ***p*** columns **t***_l_*, ***ĝ_l_*** is the estimated effect size for the predicted gene expression in tissue *l* (and thus **ĝ** is a vector with ***p*** estimated effect sizes ***ĝ_l_***), and **e** are the error terms with variance 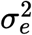. Given the high correlation between predicted expression values for a gene across different tissues, MultiXcan uses the principal components (PCs) of **T** to avoid collinearity issues. S-MultiXcan derives the joint regression estimates (effect sizes and their variances) in Equation (3) using the marginal estimates from S-PrediXcan in Equation (2). Under the null hypothesis of no association, 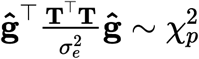, and therefore the significance of the association in S-MultiXcan is estimated with

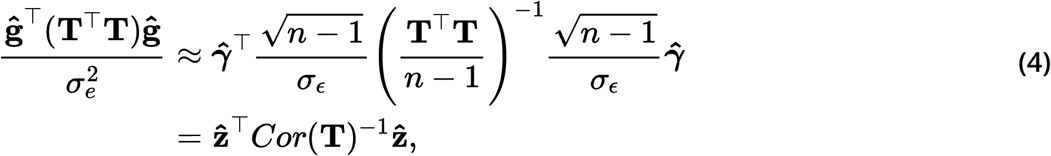

where **ẑ** is a vector with ***p*** *z*-scores (Equation (2)) for each tissue available for the gene, and ***Cor*(T)** is the autocorrelation matrix of **T**. Since **T^⊤^T** is singular for many genes, S-MultiXcan computes the pseudo-inverse ***Cor*(T)**^+^ using the ***k*** top PCs, and thus 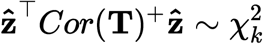. To arrive at this expression, S-MultiXcan uses the conservative approximation 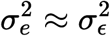, that is, the variance of the error terms in the joint regression is approximately equal to the residual variance of the marginal regressions. Another important point is that ***Cor*(T)** is estimated using a global genotype covariance matrix, whereas marginal *ẑ*_*l*_ in Equation (2) are approximated using tissue-specific genotype covariances. Although S-MultiXcan yields highly concordant estimates compared with MultiXcan, results are not perfectly correlated across genes [24]. As we explain later, these differences are important for our LV-based regression model when computing the gene-gene correlation matrix. We used S-MultiXcan results for our LV-based regression model and our cluster analyses of traits.

### TWAS resources

We used two large TWAS resources from different cohorts for discovery and replication, all obtained from European ancestries. PhenomeXcan [42], our discovery cohort, provides results on 4,091 traits across different categories. Supplemenetary File 1 has all the details about the included GWAS, sample size and disease/trait categories. In PhenomeXcan, these publicly available GWAS summary statistics were used to compute 1) gene-based associations with the PrediXcan family of methods (described before), and 2) a posterior probability of colocalization between GWAS loci and *cis*-eQTL with fastENLOC [42,96]. We refer to the matrix of *z*-scores from S-PrediXcan (Equation (2)) across *q* traits and *m* genes in tissue ***t*** as **M^*t*^** ∈ ℝ^*q*×*m*^. As explained later, matrices **M^*t*^** were used in our LV-based drug repurposing framework since they provide direction of effects. The S-MultiXcan results (22,515 gene associations across 4,091 traits) were used in our LV-based regression framework and our cluster analyses of traits. For the cluster analyses, we used the ***p***-values converted to *z*-scores: **M** = **Φ^−1^(1 − *p*/2)**, where **Φ^−1^** is the probit function. Higher *z*-scores correspond to stronger associations.

Our discovery cohort was eMERGE [46], where the same TWAS methods were run on 309 phecodes [27] across different categories (more information about traits are available in [27]). We used these results to replicate the associations found with our LV-based regression framework in PhenomeXcan.

### MultiPLIER and Pathway-level information extractor (PLIER)

MultiPLIER [44] extracts patterns of co-expressed genes from recount2 [19] (without including GTEx samples), a large gene expression dataset. The approach applies the pathway-level information extractor method (PLIER) [45], which performs unsupervised learning using prior knowledge (canonical pathways) to reduce technical noise. PLIER uses a matrix factorization approach that deconvolutes gene expression data into a set of latent variables (LV), where each LV represents a gene module. The MultiPLIER models reduced the dimensionality in recount2 to 987 LVs.

Given a gene expression dataset **Y**^*m*×*c*^ with ***m*** genes and ***c*** experimental conditions and a prior knowledge matrix **C** ∈ **{0, 1}**^*m*×*p*^ for ***p*** MSigDB pathways [100] (so that **C**_*ij*_ = 1 if gene *i* belongs to pathway *j*), PLIER finds **U, Z,** and **B** minimizing

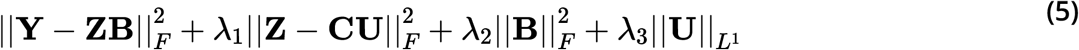

subject to **U** > 0, **Z** > 0; **Z**^*m*×*l*^ are the gene loadings with *l* latent variables, **B**^*l*×*c*^ is the latent space for *c* conditions, **U**^*p*×*l*^ specifies which of the ***p*** prior-information pathways in **C** are represented for each LV, and **λ_*i*_** are different regularization parameters used in the training step. **Z** is a low-dimensional representation of the gene space where each LV aligns as much as possible to prior knowledge, and it might represent either a known or novel gene module (i.e., a meaningful biological pattern) or noise.

For our drug repurposing and cluster analyses, we used this model to project gene-trait (from TWAS) and gene-drug associations (from LINCS L1000) into this low-dimensional gene module space. For instance, TWAS associations **M** (either from S-PrediXcan or S-MultiXcan) were projected using

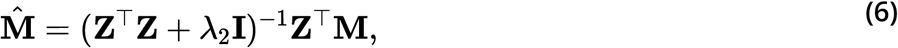

where 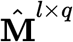 is a matrix where traits are represented by gene modules instead of single genes. As explained later, we used the same approach to project drug-induced transcriptional profiles in LINCS L1000 to obtain a representation of drugs using gene modules.

### Regression model for LV-trait associations

We adapted the gene-set analysis framework from MAGMA [101] to TWAS. We used a competitive test to predict gene-trait associations from TWAS using gene weights from an LV, testing whether top-weighted genes for an LV are more strongly associated with the phenotype than other genes with relatively small or zero weights. Thus, we fit the model

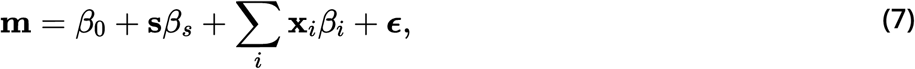

where **m** is a vector of S-MultiXcan gene ***p***-values for a trait (with a −***log*_10_** transformation); **s** is a binary indicator vector with ***s_ℓ_*** = **1** for the top 1% of genes with the largest loadings for LV *ℓ* (from **Z**_*ℓ*_) and zero otherwise; **x**_*i*_ is a gene property used as a covariate; *β* are effect sizes (with *β*_0_ as the intercept); and ***ϵ*** ∼ **MVN(0, *σ*^2^R)** is a vector of error terms with a multivariate normal distribution (MVN) where **R** is the matrix of gene correlations.

The model tests the null hypothesis *β_s_* = 0 against the one-sided hypothesis *β_s_* > 0. Therefore, *β_s_* reflects the difference in trait associations between genes that are part of LV *ℓ* and genes outside of it. Following the MAGMA framework, we used two gene properties as covariates: 1) *gene size*, defined as the number of PCs retained in S-MultiXcan, and 2) *gene density*, defined as the ratio of the number of PCs to the number of tissues available.

Since the error terms ***ϵ*** could be correlated, we cannot assume they have independent normal distributions as in a standard linear regression model. In the PrediXcan family of methods, the predicted expression of a pair of genes could be correlated if they share eQTLs or if these are in LD [95]. Therefore, we used a generalized least squares approach to account for these correlations. The gene-gene correlation matrix **R** was approximated by computing the correlations between the model sum of squares (SSM) for each pair of genes under the null hypothesis of no association. These correlations are derived from the individual-level MultiXcan model (Equation (3)), where the predicted expression matrix 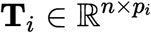 of a gene ***i*** across ***p_i_*** tissues is projected into its top ***k_i_*** PCs, resulting in matrix 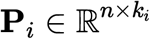. From the MAGMA framework, we know that the SSM for each gene is proportial to 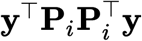. Under the null hypothesis of no association, the covariances between the SSM of genes ***i*** and ***j*** is therefore given by 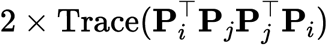. The standard deviations of each SSM are given by 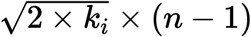. Therefore, the correlation between the SSMs for genes ***i*** and ***j*** can be written as follows:

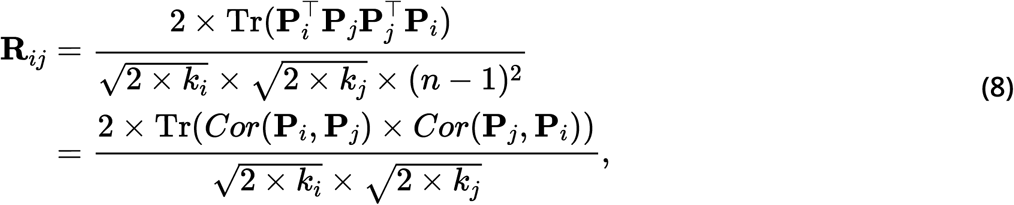

where columns **P** are standardized, **Tr** is the trace of a matrix, and the cross-correlation matrix between PCs 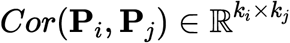 is given by

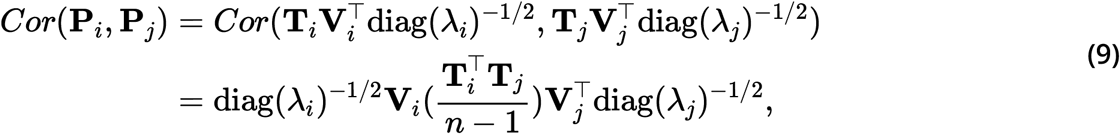

where 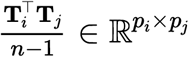 is the cross-correlation matrix between the predicted expression levels of genes ***i*** and ***j***, and columns of **V_*i*_** and scalars **λ_*i*_** are the eigenvectors and eigenvalues of **T_*i*_**, respectively. S-MultiXcan keeps only the top eigenvectors using a condition number threshold of 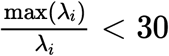. To estimate the correlation of predicted expression levels for genes ***i*** in tissue ***k*** and gene ***j*** in tissue ***l***, 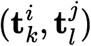 (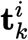 is the ***k***th column of **T_*i*_**), we used [24]

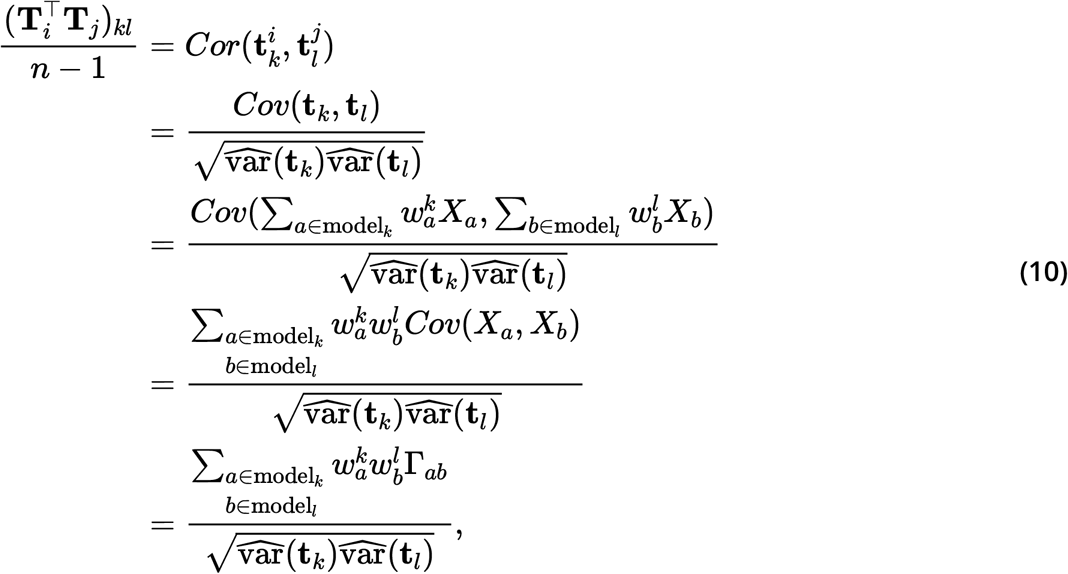

where ***X_a_*** is the genotype of SNP ***a***, 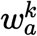 is the weight of SNP *a* for gene expression prediction in the tissue model ***k***, and 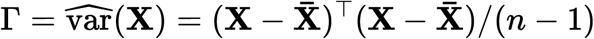 is the genotype covariance matrix using GTEx v8 as the reference panel, which is the same used in all TWAS methods described here. The variance of the predicted expression values of gene ***i*** in tissue ***k*** is estimated as [99]:

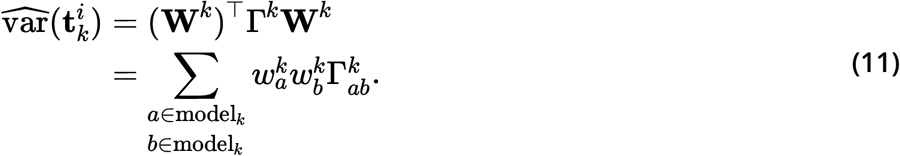

Note that, since we used the MultiXcan regression model (Equation (3)), **R** is only an approximation of gene correlations in S-MultiXcan. As explained before, S-MultiXcan approximates the joint regression parameters in MultiXcan using the marginal regression estimates from S-PrediXcan in (2) with some simplifying assumptions and different genotype covariance matrices. This complicates the derivation of an S-MultiXcan-specific solution to compute **R**. To account for this, we used a submatrix **R**_*ℓ*_ corresponding to genes that are part of LV *ℓ* only (top 1% of genes) instead of the entire matrix **R**. This simplification is conservative since correlations are accounted for top genes only. Our simulations (Supplementary Note 1) show that the model is approximately well-calibrated and can correct for LVs with adjacent and highly correlated genes at the top (e.g., Figure 9). The model can also detect LVs associated with relevant traits (Figure 2 and Table 8) that are replicated in a different cohort (Table 9).

**Table 8:**
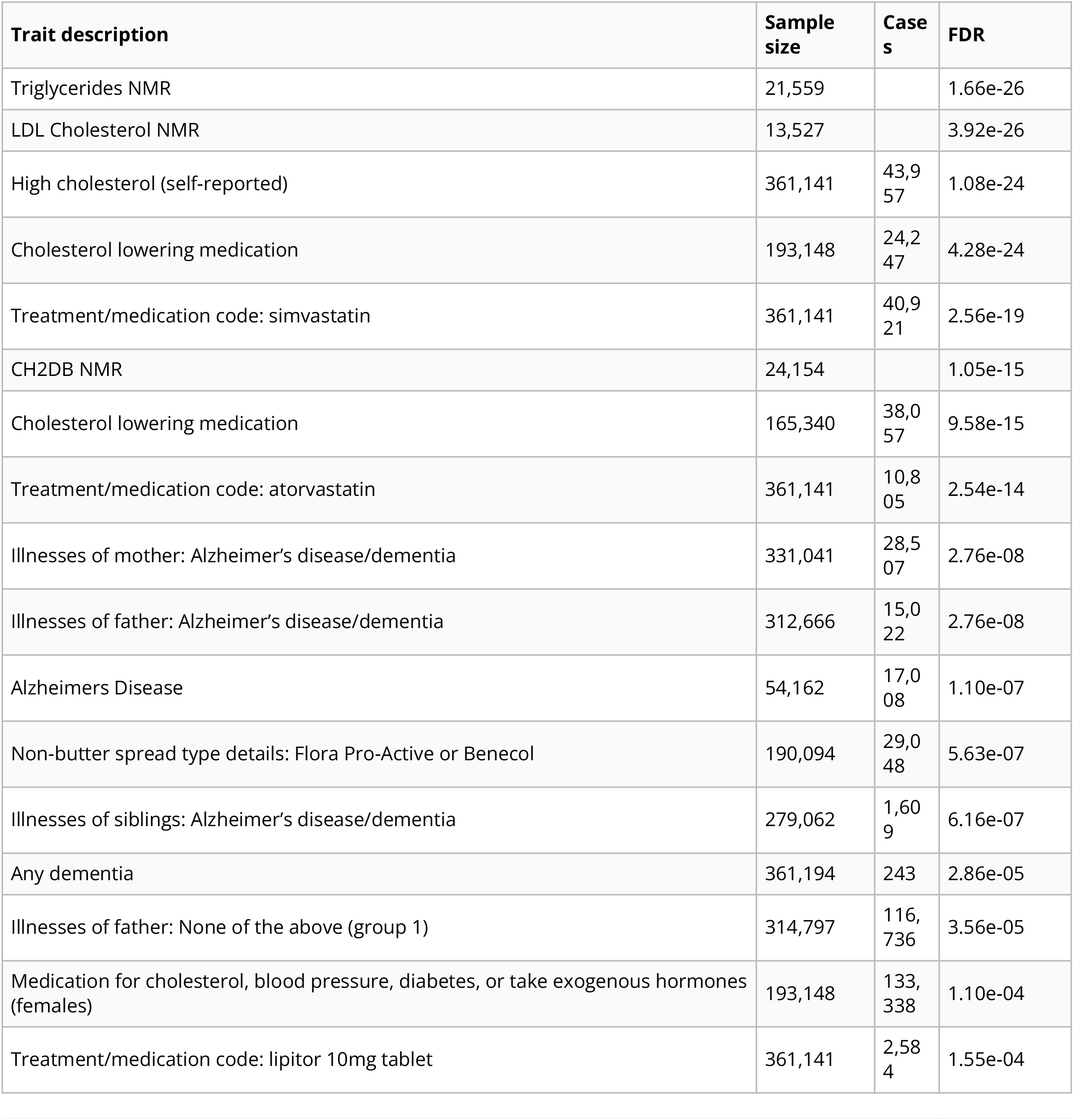
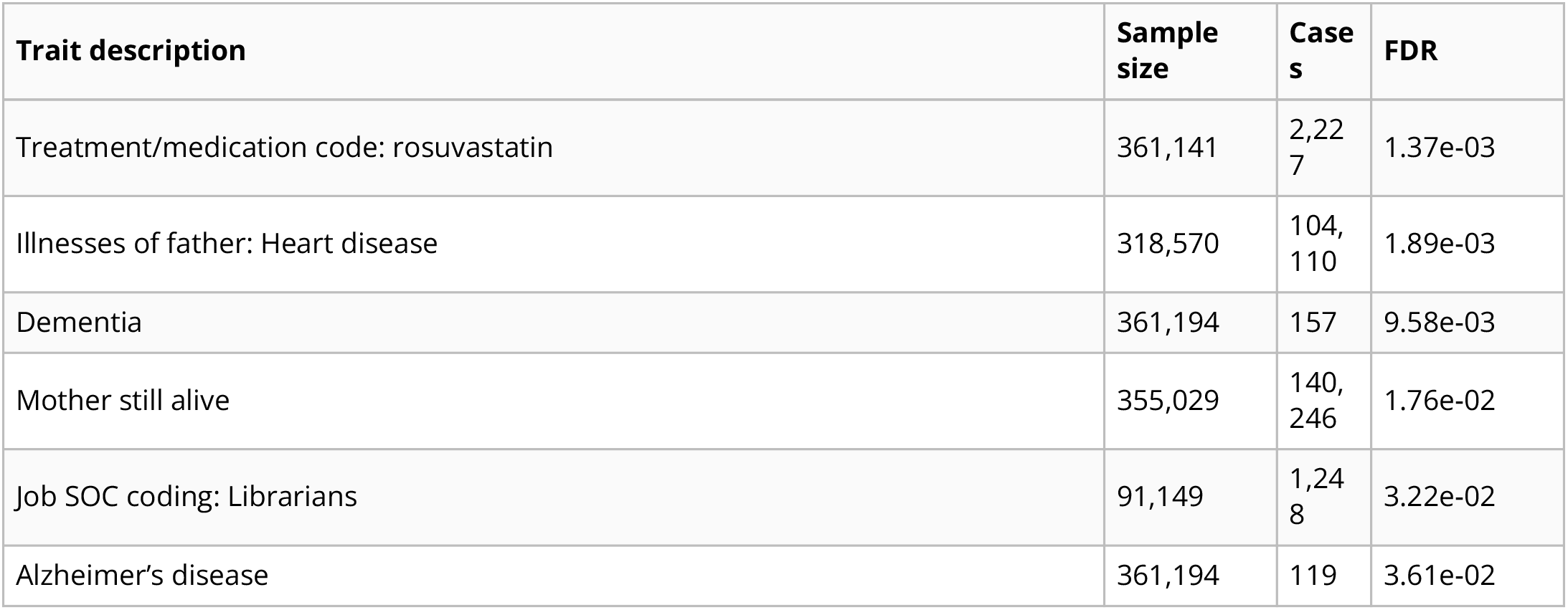
Significant trait associations of LV246 in PhenomeXcan.

**Table 9:** Significant trait associations of LV246 in eMERGE

| Phecode | Trait description | Sample size | Cases | FDR |
| --- | --- | --- | --- | --- |
| 272.11 | Hypercholesterolemia | 40,786 | 14,138 | 4.40e-09 |
| 272.1 | Hyperlipidemia | 55,843 | 29,195 | 3.57e-07 |
| 272 | Disorders of lipoid metabolism | 55,892 | 29,244 | 3.79e-07 |
| 292.3 | Memory loss | 48,785 | 2,094 | 1.80e-02 |

**Table 10:** Pathways aligned to LV116 from the MultiPLIER models.

| Pathway | AUC | FDR |
| --- | --- | --- |
| REACTOME INTERFERON SIGNALING | 0.84 | 3.48e-09 |
| SVM Macrophages M1 | 0.92 | 2.09e-05 |
| REACTOME INTERFERON ALPHA BETA SIGNALING | 0.94 | 3.36e-05 |
| REACTOME CYTOKINE SIGNALING IN IMMUNE SYSTEM | 0.67 | 1.53e-04 |
| IRIS DendriticCell-LPSstimulated | 0.65 | 1.09e-03 |
| KEGG CYTOSOLIC DNA SENSING PATHWAY | 0.84 | 3.22e-03 |
| REACTOME NEGATIVE REGULATORS OF RIG I MDA5 SIGNALING | 0.81 | 1.61e-02 |

**Table 11:**
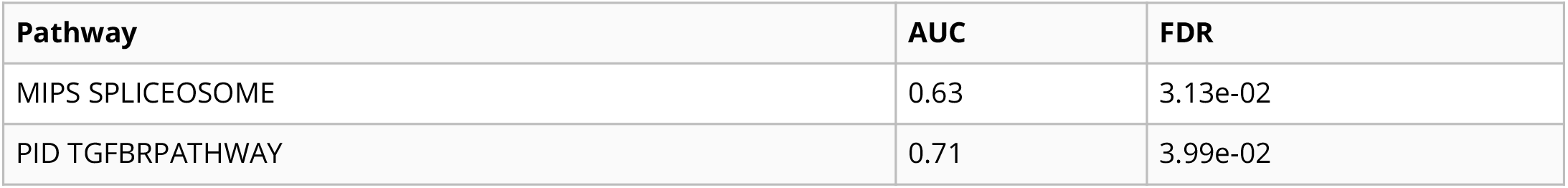
Pathways aligned to LV931 from the MultiPLIER models.

**Table 12:** Pathways aligned to LV66 from the MultiPLIER models.

| Pathway | AUC | FDR |
| --- | --- | --- |
| REACTOME METABOLISM OF LIPIDS AND LIPOPROTEINS | 0.62 | 3.12e-04 |

**Table 13:**
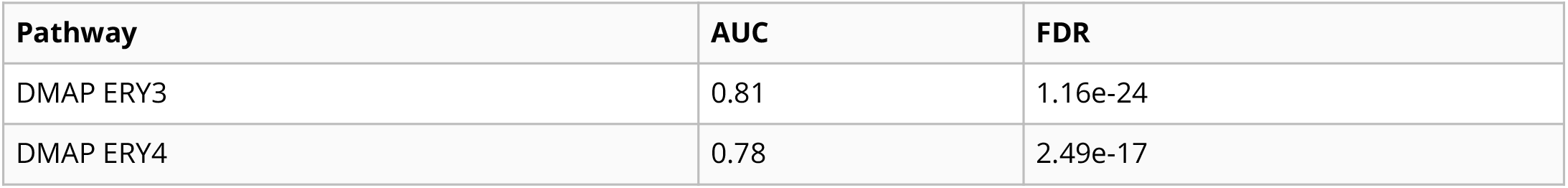
Pathways aligned to LV928 from the MultiPLIER models.

| Pathway | AUC | FDR |
| --- | --- | --- |
| DMAP ERY3 | 0.81 | 1.16e-24 |
| DMAP ERY4 | 0.78 | 2.49e-17 |

**Table 14:** Significant trait associations of LV928 in PhenomeXcan.

| Trait description | Sample size | Cases | FDR |
| --- | --- | --- | --- |
| Mean sphered cell volume | 344,729 |  | 1.60e-20 |
| Mean corpuscular haemoglobin concentration | 350,468 |  | 1.42e-17 |
| Mean reticulocyte volume | 344,728 |  | 1.77e-17 |
| Reticulocyte count | 344,729 |  | 2.28e-10 |
| Reticulocyte percentage | 344,728 |  | 1.37e-09 |
| Red blood cell (erythrocyte) distribution width | 350,473 |  | 2.90e-09 |
| Reticulocyte Count | 173,480 |  | 1.09e-07 |
| Mean corpuscular volume | 350,473 |  | 1.46e-03 |
| High light scatter reticulocyte count | 344,729 |  | 3.49e-03 |
| Age at first episode of depression | 61,033 |  | 1.33e-02 |
| High Light Scatter Reticulocyte Count | 173,480 |  | 1.48e-02 |
| Mean corpuscular haemoglobin | 350,472 |  | 4.02e-02 |

**Table 15:** Significant trait associations of LV928 in eMERGE.

| Phecode | Trait description | Sample size | Cases | FDR |
| --- | --- | --- | --- | --- |
| No significant associations |  |  |  |  |

**Table 16:** Pathways aligned to LV30 from the MultiPLIER models

| Pathway | AUC | FDR |
| --- | --- | --- |
| DMAP ERY3 | 0.95 | 5.62e-52 |
| DMAP ERY4 | 0.98 | 5.28e-51 |
| DMAP ERY5 | 0.98 | 1.96e-49 |

**Table 17:**
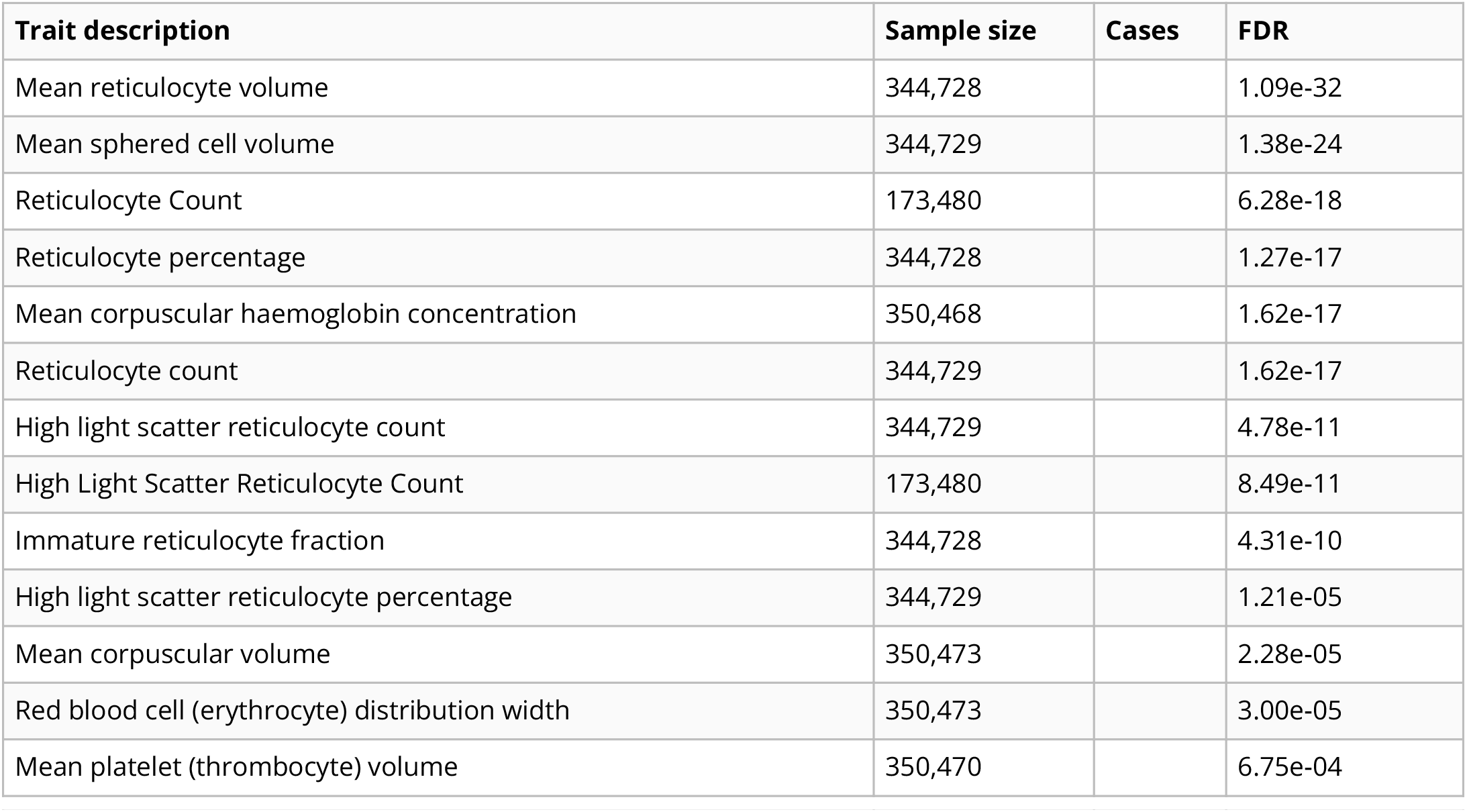
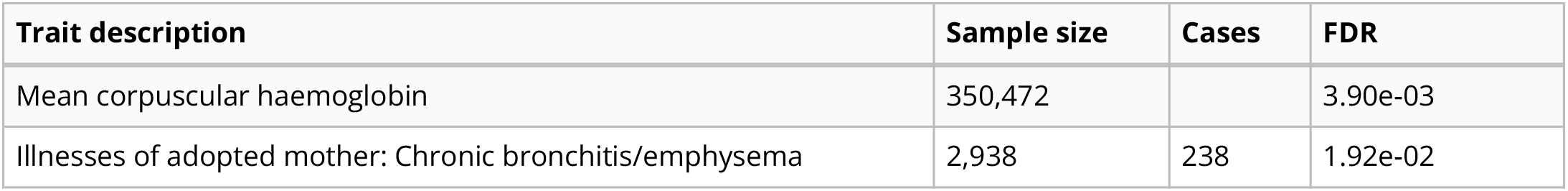
Significant trait associations of LV30 in PhenomeXcan.

**Table 18:** Significant trait associations of LV30 in eMERGE

| Phecode | Trait description | Sample size | Cases | FDR |
| --- | --- | --- | --- | --- |
| No significant associations |  |  |  |  |

**Table 19:** Pathways aligned to LV730 from the MultiPLIER models.

| Pathway | AUC | FDR |
| --- | --- | --- |
| DMAP MEGA2 | 0.82 | 2.64e-05 |

**Table 20:**
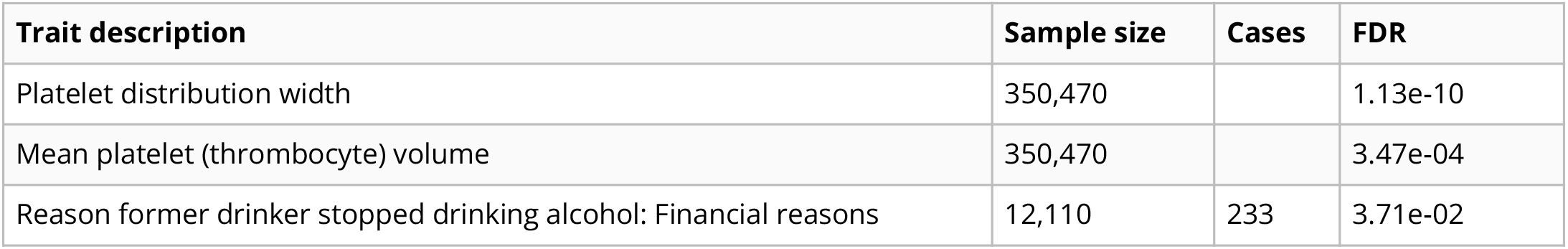
Significant trait associations of LV730 in PhenomeXcan.

**Table 21:** Significant trait associations of LV730 in eMERGE.

| Phecode | Trait description | Sample size | Cases | FDR |
| --- | --- | --- | --- | --- |
| No significant associations |  |  |  |  |

**Table 22:** Pathways aligned to LV598 from the MultiPLIER models.

| Pathway | AUC | FDR |
| --- | --- | --- |
| PID SYNDECAN 1 PATHWAY | 0.81 | 1.20e-02 |
| REACTOME COLLAGEN FORMATION | 0.77 | 1.89e-02 |

**Table 23:**
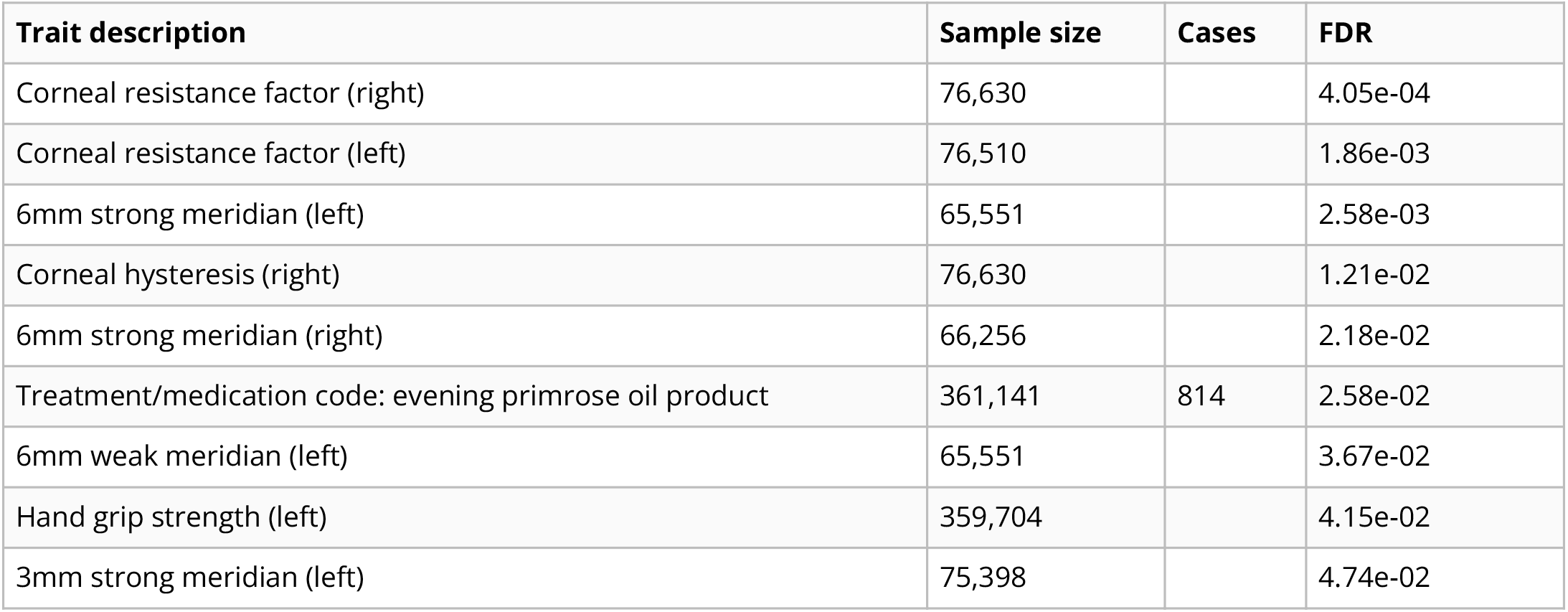
Significant trait associations of LV598 in PhenomeXcan.

**Table 24:** Significant trait associations of LV598 in eMERGE.

| Phecode | Trait description | Sample size | Cases | FDR |
| --- | --- | --- | --- | --- |
| No significant associations |  |  |  |  |

**Table 25:** Pathways aligned to LV57 from the MultiPLIER models.

| Pathway | AUC | FDR |
| --- | --- | --- |
| KEGG T CELL RECEPTOR SIGNALING PATHWAY | 0.70 | 1.26e-03 |
| SVM T cells CD4 memory activated | 0.79 | 2.59e-03 |
| IRIS CD4Tcell-Th2-restimulated12hour | 0.78 | 7.57e-03 |
| KEGG ALLOGRAFT REJECTION | 1.00 | 1.09e-02 |
| Custom Treg | 0.98 | 1.37e-02 |
| PID NFAT TFPATHWAY | 0.74 | 1.52e-02 |
| IRIS MemoryTcell-RO-activated | 0.70 | 2.87e-02 |

**Table 26:** Significant trait associations of LV57 in PhenomeXcan.

| Trait description | Sample size | Cases | FDR |
| --- | --- | --- | --- |
| Non-cancer illness code, self-reported: deep venous thrombosis (dvt) | 361,141 | 7,237 | 1.76e-13 |
| Blood clot, DVT, bronchitis, emphysema, asthma, rhinitis, eczema, allergy diagnosed by doctor: Blood clot in the leg (DVT) | 360,527 | 7,386 | 1.22e-12 |
| Diagnoses - main ICD10: I80 Phlebitis and thrombophlebitis | 361,194 | 2,289 | 7.62e-12 |
| DVT of lower extremities | 361,194 | 2,116 | 1.27e-09 |
| Venous thromboembolism | 361,194 | 4,620 | 2.28e-08 |
| DVT of lower extremities and pulmonary embolism | 361,194 | 4,319 | 4.36e-08 |
| Inflammatory Bowel Disease | 34,652 | 12,882 | 1.95e-05 |
| hypothyroidism (self-reported) | 361,141 | 17,574 | 3.84e-05 |
| Medication: levothyroxine sodium | 361,141 | 14,689 | 8.43e-05 |
| Mouth/teeth dental problems: Mouth ulcers | 359,841 | 36,831 | 1.02e-03 |
| Crohns Disease | 20,833 | 5,956 | 1.02e-02 |
| Facial ageing | 330,409 |  | 1.04e-02 |
| Ulcerative Colitis | 27,432 | 6,968 | 1.27e-02 |
| Hair colour (natural, before greying): Black | 360,270 | 15,809 | 1.99e-02 |
| Hair colour (natural, before greying): Light brown | 360,270 | 147,560 | 4.69e-02 |

**Table 27:** Significant trait associations of LV57 in eMERGE.

| Phecode | Trait description | Sample size | Cases | FDR |
| --- | --- | --- | --- | --- |
| 286 | Coagulation defects | 50,182 | 2,976 | 1.33e-11 |
| 452 | Other venous embolism and thrombosis | 40,476 | 3,816 | 1.52e-05 |
| 452.2 | Deep vein thrombosis [DVT] | 38,791 | 2,131 | 4.47e-05 |
| 244.4 | Hypothyroidism NOS | 53,968 | 9,284 | 1.12e-02 |
| 244 | Hypothyroidism | 54,404 | 9,720 | 1.42e-02 |

**Table 28:** Pathways aligned to LV844 from the MultiPLIER models.

| Pathway | AUC | FDR |
| --- | --- | --- |
| KEGG ANTIGEN PROCESSING AND PRESENTATION | 0.80 | 1.35e-03 |

**Table 29:** Significant trait associations of LV844 in PhenomeXcan

| Trait description | Sample size | Cases | FDR |
| --- | --- | --- | --- |
| Non-cancer illness code, self-reported: polymyalgia rheumatica | 361,141 | 753 | 5.22e-06 |
| Non-cancer illness code, self-reported: type 1 diabetes | 361,141 | 318 | 4.71e-05 |
| Type 1 diabetes with ketoacidosis | 361,194 | 168 | 1.03e-04 |
| Age diabetes diagnosed | 16,166 |  | 3.86e-04 |
| Milk type used: Other type of milk | 360,806 | 4,213 | 5.48e-04 |
| Non-cancer illness code, self-reported: appendicitis | 361,141 | 3,058 | 6.12e-04 |
| Diabetic ketoacidosis | 361,194 | 234 | 7.13e-04 |
| Rheumatoid Arthritis | 80,799 | 19,234 | 7.46e-04 |
| Type 1 diabetes without complications | 361,194 | 247 | 1.05e-03 |
| Started insulin within one year diagnosis of diabetes | 16,415 | 1,999 | 1.30e-03 |
| Insulin medication (males) | 165,340 | 2,248 | 3.61e-03 |
| Medication: insulin product | 361,141 | 3,545 | 5.48e-03 |
| Insulin medication (females) | 193,148 | 1,476 | 7.93e-03 |
| Type 1 diabetes | 361,194 | 583 | 1.04e-02 |
| Diagnoses - main ICD10: E10 Insulin-dependent diabetes mellitus | 361,194 | 470 | 1.08e-02 |
| Treatment/medication code: sulfasalazine | 361,141 | 710 | 1.10e-02 |
| malabsorption/coeliac disease (self-reported) | 361,141 | 1,587 | 3.12e-02 |
| Job coding: school inspector, education inspector | 89,866 | 238 | 3.71e-02 |
| Seropositive rheumatoid arthritis | 361,194 | 327 | 3.86e-02 |
| Non-cancer illness code, self-reported: rheumatoid arthritis | 361,141 | 4,017 | 4.21e-02 |
| Age hayfever or allergic rhinitis diagnosed by doctor | 20,904 |  | 4.44e-02 |
| Other/unspecified seropositiverheumatoid arthritis | 361,194 | 299 | 4.88e-02 |

**Table 30:**
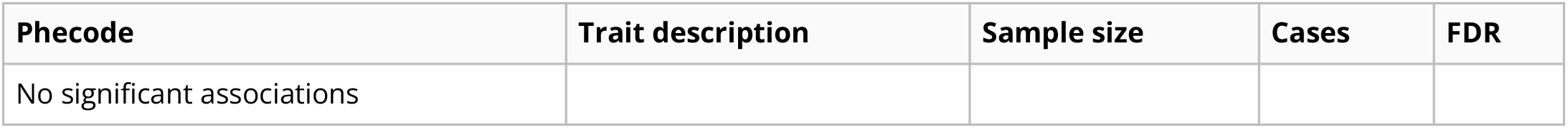
Significant trait associations of LV844 in eMERGE.

| Phecode | Trait description | Sample size | Cases | FDR |
| --- | --- | --- | --- | --- |
| No significant associations |  |  |  |  |

**Table 31:** Pathways aligned to LV136 from the MultiPLIER models.

| Pathway | AUC | FDR |
| --- | --- | --- |
| PID INTEGRIN1 PATHWAY | 0.88 | 9.35e-06 |
| KEGG ECM RECEPTOR INTERACTION | 0.80 | 7.29e-05 |
| REACTOME COLLAGEN FORMATION | 0.87 | 2.00e-04 |
| REACTOME MUSCLE CONTRACTION | 0.75 | 1.49e-02 |

**Table 32:** Significant trait associations of LV136 in PhenomeXcan.

| Trait description | Sample size | Cases | FDR |
| --- | --- | --- | --- |
| Coronary atherosclerosis | 361,194 | 14,334 | 1.84e-09 |
| Chronic ischaemic heart disease (ICD10 I25) | 361,194 | 12,769 | 3.52e-09 |
| Ischaemic heart disease (wide definition) | 361,194 | 20,857 | 3.95e-08 |
| Coronary Artery Disease | 184,305 | 60,801 | 4.18e-08 |
| 3mm strong meridian (right) | 75,410 |  | 5.54e-05 |
| 6mm strong meridian (left) | 65,551 |  | 1.35e-04 |
| Corneal resistance factor (right) | 76,630 |  | 2.02e-04 |
| 6mm strong meridian (right) | 66,256 |  | 2.58e-04 |
| Heart attack | 360,420 | 8,288 | 3.75e-04 |
| Myocardial infarction | 361,194 | 7,018 | 4.85e-04 |
| Myocardial infarction, strict | 361,194 | 7,018 | 4.85e-04 |
| 3mm strong meridian (left) | 75,398 |  | 6.65e-04 |
| heart attack/myocardial infarction (self-reported) | 361,141 | 8,239 | 1.07e-03 |
| 6mm weak meridian (left) | 65,551 |  | 1.10e-03 |
| 6mm weak meridian (right) | 66,256 |  | 1.61e-03 |
| Acute myocardial infarction (ICD10 I21) | 361,194 | 5,948 | 2.24e-03 |
| 3mm weak meridian (right) | 75,410 |  | 3.69e-03 |
| 3mm weak meridian (left) | 75,398 |  | 3.96e-03 |
| Intra-ocular pressure, Goldmann-correlated (right) | 76,630 |  | 8.64e-03 |
| 6mm asymmetry angle (right) | 41,390 |  | 1.03e-02 |
| Corneal resistance factor (left) | 76,510 |  | 1.03e-02 |
| Other specified disorders of muscle | 361,194 | 257 | 1.09e-02 |
| Major coronary heart disease event excluding revascularizations | 361,194 | 10,157 | 2.44e-02 |
| Major coronary heart disease event | 361,194 | 10,157 | 2.44e-02 |
| Non-cancer illness code, self-reported: angina | 361,141 | 11,370 | 4.53e-02 |
| Eye problems/disorders: Glaucoma | 117,890 | 5,092 | 4.94e-02 |

**Table 33:**
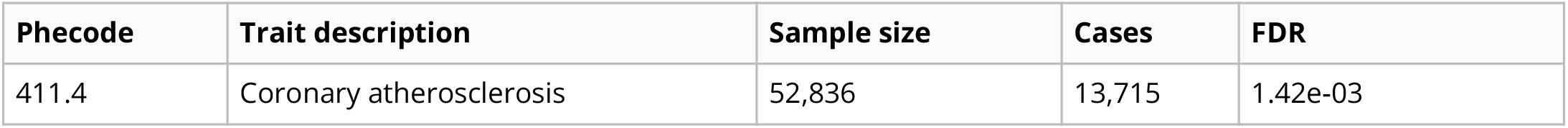
Significant trait associations of LV136 in eMERGE.

| Phecode | Trait description | Sample size | Cases | FDR |
| --- | --- | --- | --- | --- |
| 411.4 | Coronary atherosclerosis | 52,836 | 13,715 | 1.42e-03 |

**Table 34:**
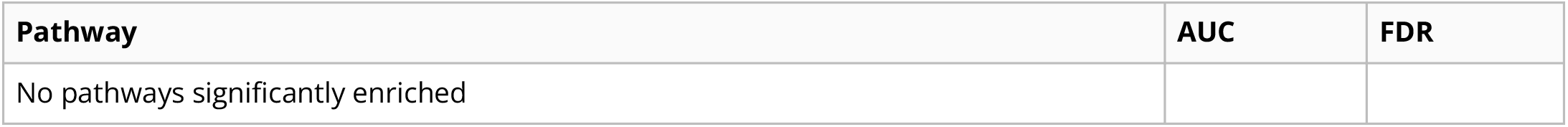
Pathways aligned to LV21 from the MultiPLIER models.

| Pathway | AUC | FDR |
| --- | --- | --- |
| No pathways significantly enriched |  |  |

**Table 35:**
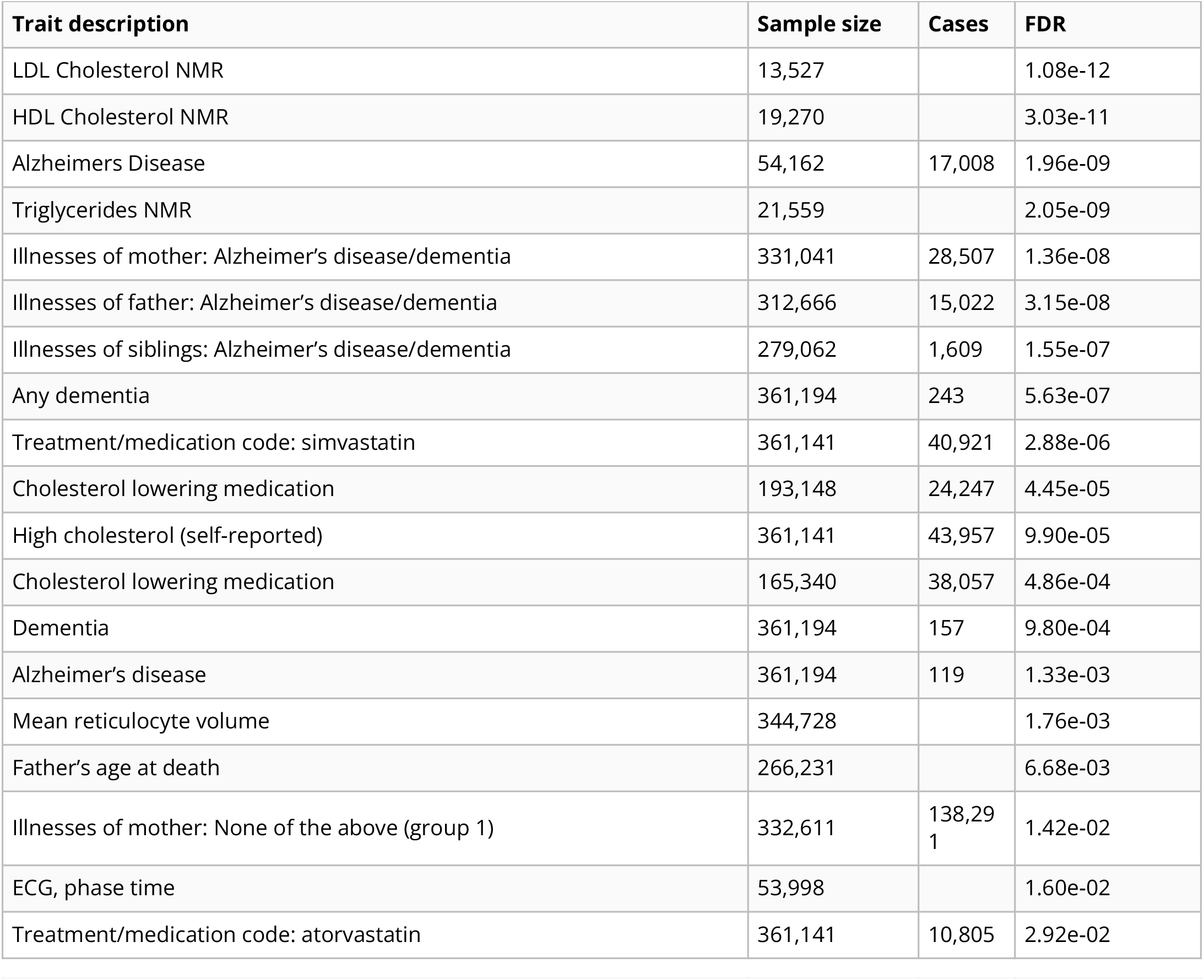
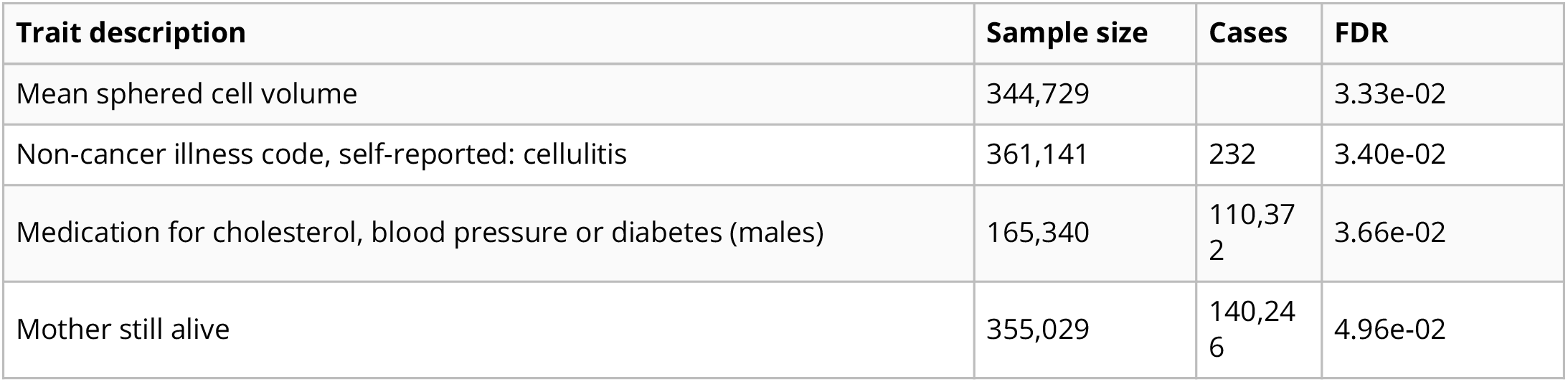
Significant trait associations of LV21 in PhenomeXcan.

**Table 36:**
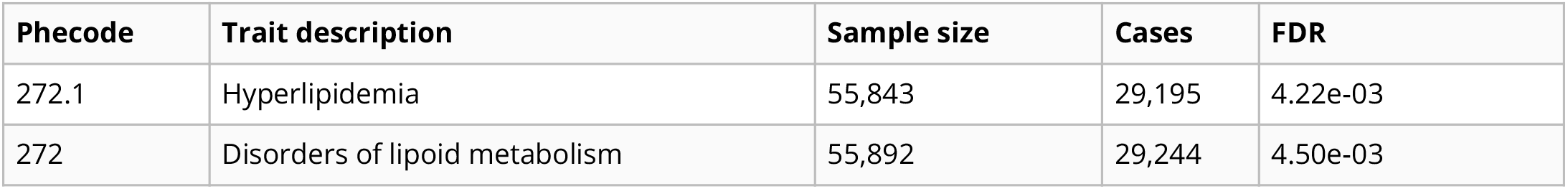
Significant trait associations of LV21 in eMERGE.

In Equation (10), for each gene, we only considered tissue models present in S-PrediXcan results, as well as SNPs present in GWAS used as input for the TWAS approaches. This is necessary to obtain more accurate correlations estimates [24]. Therefore, we computed different correlation matrices for PhenomeXcan and eMERGE. In PhenomeXcan, most of the GWAS (4,049) were obtained from the UK Biobank using the same pipeline and including the same set of SNPs, so a single correlation matrix was used for this set. For the rest, we used a single correlation matrix for each group of traits that shared the same or most of the SNPs.

We ran our regression model for all 987 LVs across the 4,091 traits in PhenomeXcan. For replication, we ran the model in the 309 phecodes in eMERGE. We adjusted the ***p***-values using the Benjamini-Hochberg procedure.

### LV-based drug repurposing approach

For the drug-disease prediction, we derived an LV-based method based on a drug repositioning framework previously used for psychiatry traits [30], where individual/single genes associated with a trait are anticorrelated with expression profiles for drugs. We compared our LV-based method with this previously published, single-gene approach. For the single-gene method, we computed a drug-disease score by multiplying each S-PrediXcan set of signed *z*-scores in tissue ***t***, **M^*t*^**, with another set of signed *z*-scores from transcriptional responses profiled in LINCS L1000 [43], **L^*c*×*m*^** (for ***c*** compounds). Here **M^*t*^** contains information about whether a higher or lower predicted expression of a gene is associated with disease risk, whereas **L** indicates whether a drug increases or decreases the expression of a gene. Therefore, these two matrices can be multiplied to compute a score for a drug-disease pair. The result of this product is **D^*t,k*^** = −**1 ⋅ M^*t,k*^L**^⊤^, where ***k*** refers to the number of most significant gene associations in **M^*t*^** for each trait. As suggested in [30], ***k*** could be either all genes or the top 50, 100, 250, and 500; then, we averaged score ranks across all ***k*** and obtained **D^*t*^**. Finally, for each drug-disease pair, we took the maximum prediction score across all tissues: 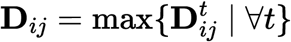.

The same procedure was used for the LV-based approach, where we projected **M^*t*^** and **L** into the gene module latent space using Equation (6), leading to 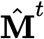 and 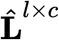, respectively. Finally, 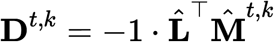, where in this case ***k*** could be all LVs or the top 5, 10, 25 and 50 (since we have an order of magnitude less LVs than genes).

Since the gold standard of drug-disease medical indications is described with Disease Ontology IDs (DOID) [102], we mapped PhenomeXcan traits to the Experimental Factor Ontology [103] using [104], and then to DOID.

### Consensus clustering of traits

We performed two preprocessing steps on the S-MultiXcan results before the cluster analysis. First, we combined results in **M** (with ***p***-values converted to *z*-scores, as described before) for traits that mapped to the same Experimental Factor Ontology (EFO) [103] term using the Stouffer’s method: 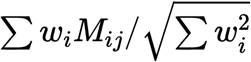, where ***w_i_*** is a weight based on the GWAS sample size for trait ***i***, and ***M_ij_*** is the *z*-score for gene ***j***. Second, we divided all *z*-scores for each trait ***i*** by their sum to reduce the effect of highly polygenic traits: ***M_ij_*/ ∑ *M_ij_***. Finally, we projected this data matrix using Equation (6), obtaining 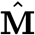 with ***n***=3,752 traits and ***l***=987 LVs as the input of our clustering pipeline.

A partitioning of 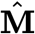 with ***n*** traits into ***k*** clusters is represented as a label vector *π* ∈ ℕ^*n*^. Consensus clustering approaches consist of two steps: 1) the generation of an ensemble **Π** with ***r*** partitions of the dataset: **Π** = **{*π*_1_, *π*_2_, … , *π_r_*}**, and 2) the combination of the ensemble into a consolidated solution defined as:

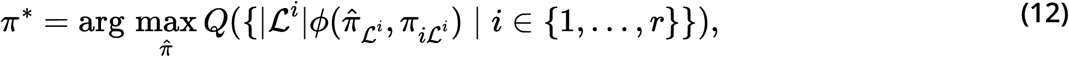

where 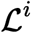 is a set of data indices with known cluster labels for partition ***i*, *ϕ*:** ℕ^*n*^ × ℕ^*n*^ → ℝ is a function that measures the similarity between two partitions, and ***Q*** is a measure of central tendency, such as the mean or median. We used the adjusted Rand index (ARI) [105] for ***ϕ*** and the median for ***Q***. To obtain ***π***\*, we define a consensus function **Γ:** ℕ^*n*×*r*^ → ℕ^*n*^ with **Π** as the input. We used consensus functions based on the evidence accumulation clustering (EAC) paradigm [79], where **Π** is first transformed into a distance matrix **D_*ij*_** = ***d_ij_*/*r***, where ***d_ij_*** is the number of times traits ***i*** and ***j*** were grouped in different clusters across all ***r*** partitions in **Π**. Then, **Γ** can be any similarity-based clustering algorithm, which is applied on **D** to derive the final partition ***π***\*.

For the ensemble generation step, we used different algorithms to create a highly diverse set of partitions (see Figure 5) since diversity is an important property for ensembles [106,107,108]. We used three data representations: the raw dataset, its projection into the top 50 principal components, and the embedding learned by UMAP [109] using 50 components. For each of these, we applied five clustering algorithms covering a wide range of different assumptions on the data structure: ***k***-means [110], spectral clustering [111], a Gaussian mixture model (GMM), hierarchical clustering, and DBSCAN [112]. For ***k***-means, spectral clustering and GMM, we specified a range of ***k*** between 2 and 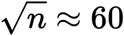, and for each ***k*** we generated five partitions using random seeds. For hierarchical clustering, for each ***k***, we generated four partitions using common linkage criteria: ward, complete, average and single. For DBSCAN, we combined different ranges for parameters *ϵ* (the maximum distance between two data points to be considered part of the same neighborhood) and *minPts* (the minimum number of data points in a neighborhood for a data point to be considered a core point), based on the procedure in [113]. Specifically, we used *minPts* values from 2 to 125. For each data representation (raw, PCA and UMAP), we determined a plausible range of *ϵ* values by observing the distribution of the mean distance of the *minPts*-nearest neighbors across all data points. Since some combinations of *minPts* and *ϵ* might not produce a meaningful partition (for instance, when all points are detected as noisy or only one cluster is found), we resampled partitions generated by DBSCAN to ensure an equal representation of this algorithm in the ensemble. This procedure generated a final ensemble of 4,428 partitions of 3,752 traits.

Finally, we used spectral clustering on **D** to derive the final consensus partitions. **D** was first transformed into a similarity matrix by applying an RBF kernel exp(−*γ***D^2^**) using four different values for *γ* that we empirically determined to work best. Therefore, for each ***k*** between 2 and 60, we derived four consensus partitions and selected the one that maximized Equation (12). We further filtered this set of 59 solutions to keep only those with an ensemble agreement larger than the 75th percentile (Supplementary Figure 15), leaving a total of 15 final consensus partitions shown in Figure 6.

The input data in our clustering pipeline undergoes several linear and nonlinear transformations, including PCA, UMAP and the ensemble transformation using the EAC paradigm (distance matrix **D**). Although consensus clustering has clear advantages for biological data [114], this set of data transformations complicates the interpretation of results. To circumvent this, we used a supervised learning approach to detect which gene modules/LVs are the most important for each cluster of traits (Figure 5b). Note that we did not use this supervised model for prediction but only to learn which features (LVs) were most discriminative for each cluster. For this, we used the highest resolution partition (***k***=29, although any could be used) to train a decision tree model using each of the clusters as labels and the projected data 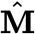 as the training samples. For each ***k***, we built a set of binary labels with the current cluster’s traits as the positive class and the rest of the traits as the negative class. Then, we selected the LV in the root node of the trained model only if its threshold was positive and larger than one standard deviation. Next, we removed this LV from 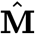 (regardless of being previously selected or not) and trained the model again. We repeated this procedure 20 times to extract the top 20 LVs that better discriminate traits in a cluster from the rest.

In Supplementary Note 2, we performed several analyses under a null hypothesis of no structure in the data to verify that the clustering results detected by this pipeline were real.

### CRISPR-Cas9 screening

#### Cell culture

HepG2 cells were obtained from ATCC (ATCC® HB-8065™), and maintained in Eagle’s Minimum Essential Medium with L-Glutamine (EMEM, Cat. 112-018-101, Quality Biology) supplemented with 10% Fetal Bovine Serum (FBS, Gibco, Cat.16000-044), and 1% Pen/Strep (Gibco, Cat.15140-122). Cells were kept at 37oC in a humidity-controlled incubator with 5% CO2, and were maintained at a density not exceeding more than 80% confluency.

#### Genome-wide lentiviral pooled CRISPR-Cas9 library

3rd lentiviral generation, Broad GPP genome-wide Human Brunello CRISPR knockout Pooled library was provided by David Root and John Doench from Addgene (Cat. 73179-LV), and was used for HepG2 cell transduction. It consists of 76,441 sgRNAs, and targets 19,114 genes in the human genome with an average of 4 sgRNAs per gene. Each 20nt sgRNA cassette was inserted into the lentiCRIS-PRv2 backbone between U6 promoter and gRNA scaffold. Through cell transduction, the lentiviral vectors which encode Cas9 were used to deliver the sgRNA cassette containing plasmids into cells during cell replication. Unsuccessful transduced cells were excluded through puromycin selection.

#### Lentiviral titer determination

No-spin lentiviral transduction was utilized for the screen. In a Collagen-I coated 6-wells plate, approximate 2.5 M cells were seeded each well in the presence of 8ug/ml polybrene (Millipore Sigma, Cat. TR-1003 G), and a different titrated virus volume (e.g., 0, 50, 100, 200, 250, and 400ul) was assigned to each well. EMEM complete media was added to make the final volume of 1.24ml. 16-18hrs post-transduction, virus/polybrene-containing media was removed from each well. Cells were washed twice with 1x DPBS and replaced with fresh EMEM. At 24h, cells in each well were trypsinized, diluted (e.g.,1:10), and seeded in pairs of wells of 6-well plates. At 60hr post-transduction, cell media in each well was replaced with fresh EMEM. 2ug/ml of puromycin (Gibco, Cat. A1113803) was added to one well out of the pair. 2-5 days after puromycin selection, or the 0 virus well treated with puromycin had no survival of cells, cells in both wells with/without puromycin were collected and counted for viability. Percentage of Infection (PI%) was obtained by comparing the cell numbers with/without puromycin selection within each pair. By means of Poisson’s distribution theory, when transduction efficiency (PI%) is between 30-50%, which corresponds to an MOI (Multiplicity of Infection) of ~0.35-0.70. At MOI equal to or close to 0.3, around 95% of infected cells are predicted to have only one copy of the virus. Therefore, a volume of virus (120ul) yielding 30-40% of transduction efficiency was chosen for further large-scale viral transduction.

#### Lentiviral Transduction in HepG2 Using Brunello CRISPR Knockout Pooled Library

In order to achieve a coverage (representation) of at least 500 cells per sgRNA, and at an MOI between 0.3-0.4 to ensure 95% of infected cells get only one viral particle per cell, ~200M cells were initiated for the screen. Transduction was carried out in a similar fashion as described above. Briefly, 2.5M cells were seeded in each well of 14 6-well plates, along with 8ug/ml of polybrene. A volume of 120ul of the virus was added to each experimental well. 18hrs post-transduction, virus/PB mix medium was removed, and cells in each well were collected, counted, and pooled into T175 flasks. At 60hr post-transduction, 2ug/ml of puromycin was added to each flask. Mediums were changed every two days with fresh EMEM, topped with 2ug/ml puromycin. Seven days after puromycin selection, cells were collected, pooled, counted, and replated.

#### Fluorescent dye staining

9 days after puromycin selection, cells were assigned to 2 groups. 20-30M cells were collected as Unsorted Control. The cell pellet was spun down at 500 x g for 5min at 4oC. The dry pellet was kept at −80oC for further genomic DNA isolation. The rest of the cells (approximately 200M) were kept in 100mm dishes and stained with a fluorescent dye (LipidSpotTM 488, Biotium, Cat. 70065-T). In Brief, LipidSpot 488 was diluted to 1:100 with DPBS. 4ml of staining solution was used for each dish and incubated at 37oC for 30min. Cell images were captured through fluorescent microscope EVOS for GFP signal detection (Figure S1).

#### Fluorescence-activated cell sorting (FACS)

Cells were immediately collected into 50ml tubes (From this point on, keep cells cold), and spun at 500 x g for 5min at 4oC. After DPBS wash, cell pellets were resuspended with FACS Sorting Buffer (1x DPBS without Ca2+/Mg2+, 2.5mM EDTA, 25mM HEPES, 1% BSA. The solution was filter sterilized, and kept at 4oC), pi-pet gently to make single cells. The cell solution was then filtered through a cell strainer (Falcon, Cat. 352235) and was kept on ice, protected from light. Collected cells were sorted on FACSJazz. 100um nozzle was used for sorting. ~20% of each GFP-High and GFP-Low (Figure S2) were collected into 15ml tubes. After sorting, cells were immediately spun down. Pellets were kept at −80oC for further genomic DNA isolation.

#### Genomic DNA isolation and verification

Three conditions of Genomic DNA (Un-Sorted Control, lentiV2 GFP-High, and lentiV2 GFP-Low) were extracted using QIAamp DNA Blood Mini Kit (Qiagen, Cat.51104), followed by UV Spectroscopy (Nanodrop) to access the quality and quantity of the gDNA. A total of 80-160ug of gDNA was isolated for each condition. sgRNA cassette and lentiviral specific transgene in isolated gDNA were verified through PCR (Figure S3).

#### Illumina libraries generation and sequencing

The fragment containing sgRNA cassette was amplified using P5 /P7 primers, as indicated in [115], and primer sequences were adapted from Broad Institute protocol (Figure S4). Stagger sequence (0-8nt) was included in P5 and 8bp uniquely barcoded sequence in P7. Primers were synthesized through Integrated DNA Technologies (IDT), and each primer was PAGE purified. 32 PCR reactions were set up for each condition. Each 100ul PCR reaction consists of roughly 5ug of gDNA, 5ul of each 10uM P5 and P7. ExTaq DNA Polymerase (TaKaRa, Cat. RR001A) was used to amplify the amplicon. PCR Thermal Cycler Parameters set as Initial at 95oC for 1min; followed by 24 cycles of Denaturation at 94oC for 30 seconds, Annealing at 52.5oC for 30 seconds, Extension at 72oC for 30 seconds. A final Elongation at 72oC for 10 minutes. 285bp-293bp PCR products were expected (Figure S5 A). PCR products within the same condition were pooled and purified using SPRIselect beads (Beckman Coulter, Cat. B23318). Purified Illumina libraries were quantitated on Qubit, and the quality of the library was analyzed on Bio-analyzer using High Sensitivity DNA Chip. A single approximate 285bp peak was expected. (Figure S5 B). Final Illumina library samples were sequenced on Nova-seq 6000. Samples were pooled and loaded on an SP flow cell, along with a 20% PhiX control v3 library spike-in.

## Code and data availability

The code and data to reproduce all the analyses in this work are available in https://github.com/greenelab/phenoplier.

## Acknowledgements

Figure 1a was created with BioRender.com.

## Supplementary material

### Regression model for LV-trait associations

#### Supplementary Note 1: mean type I error rates and calibration of LV-based regression model

We assessed our GLS model type I error rates (proportion of ***p***-values below 0.05) and calibration using a null model of random traits and genotype data from 1000 Genomes Phase III. We selected 312 individuals with European ancestry, and then analyzed 1,000 traits drawn from a standard normal distribution 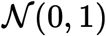. We ran all the standard procedures for the TWAS approaches (S-PrediXcan and S-MultiXcan), including: 1) a standard GWAS using linear regression under an additive genetic model, 2) different GWAS processing steps, including harmonization and imputation procedures as defined in [116], 3) S-PrediXcan and S-MultiXcan analyses. Below we provide details for each of these steps.

##### Step 1 - GWAS

We performed standard QC procedures such as filtering out variants with missing call rates eexceeding 0.01, MAF below 1% or MAC below 20, and HWE below 1e-6, and removing samples with high sex-discrepancy and high-relatedness (first and second degree). We included sex and the top 20 principal components as covariates, performing the association test on 5,923,554 variants across all 1,000 random phenotypes.

##### Step 2 - GWAS processing

These steps include harmonization of GWAS and imputation of *z*-scores, which are part of the TWAS pipeline and are needed in order to ensure an acceptable overlap with SNPs in prediction models. The scripts to run these steps are available in [117]. These procedures were run for all 1,000 random phenotypes and generated a total number of 8,325,729 variants, including those with original and imputed *z*-scores.

##### Step 3 - TWAS

We processed the imputed GWAS with S-PrediXcan using the MASHR prediction models on 49 tissues from GTEx v8. Then, S-MultiXcan was ran using the GWAS and S-PrediXcan outputs to generate gene-trait association ***p***-values.

Finally, we ran our GLS model (Equation (7)) to compute an association between each of the 987 LVs in MultiPLIER and the 1,000 S-MultiXcan results on random phenotypes. For this, we built a gene correlation matrix specifically for this cohort (see Methods). Then, we compared the GLS results with an equivalent, baseline ordinarly least squares (OLS) model assuming independence between genes. Figure 8 compares the distribution of ***p***-values of the OLS and GLS models. The GLS model has a slightly smaller mean type I error rate (0.0558, SD=0.0127) than the baseline OLS model (0.0584, SD=0.0140), and ***p***-values follow more closely the expected uniform distribution. Importantly, the GLS model is able to correct for LVs with adjacent and highly correlated genes at the top such as LV234 (Figure 9), LV847 (Figure 10), LV45 (Figure 11), or LV800 (Figure 12), among others. In contrast and as expected, the OLS model has higher mean type I errors and smaller-than-expected ***p***-values in all these cases.

**Figure 8:**
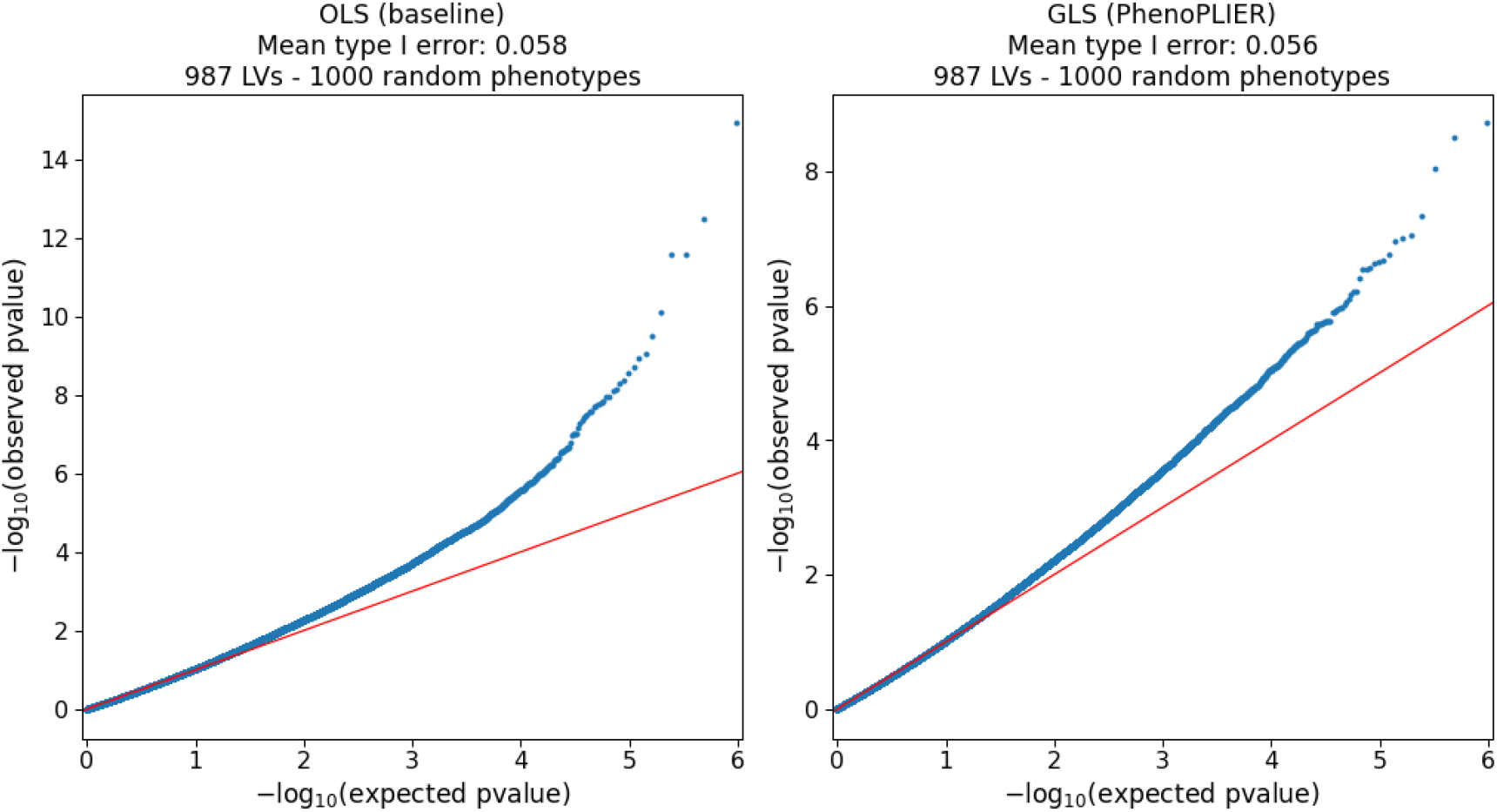
QQ-plots for OLS (baseline) and GLS (PhenoPLIER) models on random phenotypes.

**Figure 9:**
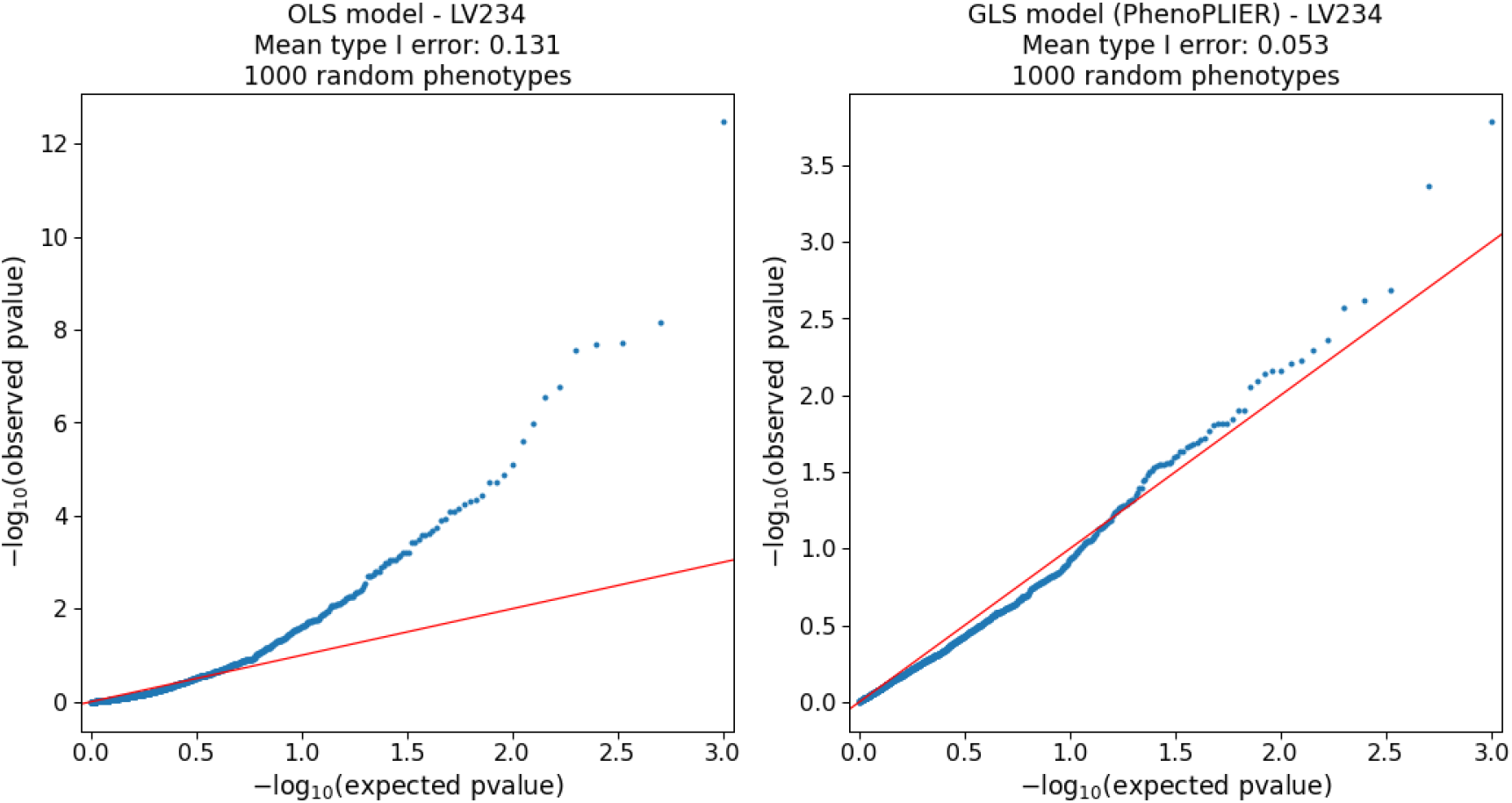
QQ-plots for LV234 on random phenotypes. Among the top 1% of genes in this LV, 17 are located in band 6p22.2, 5 in 6p22.1 and 3 in 7q11.23.

**Figure 10:**
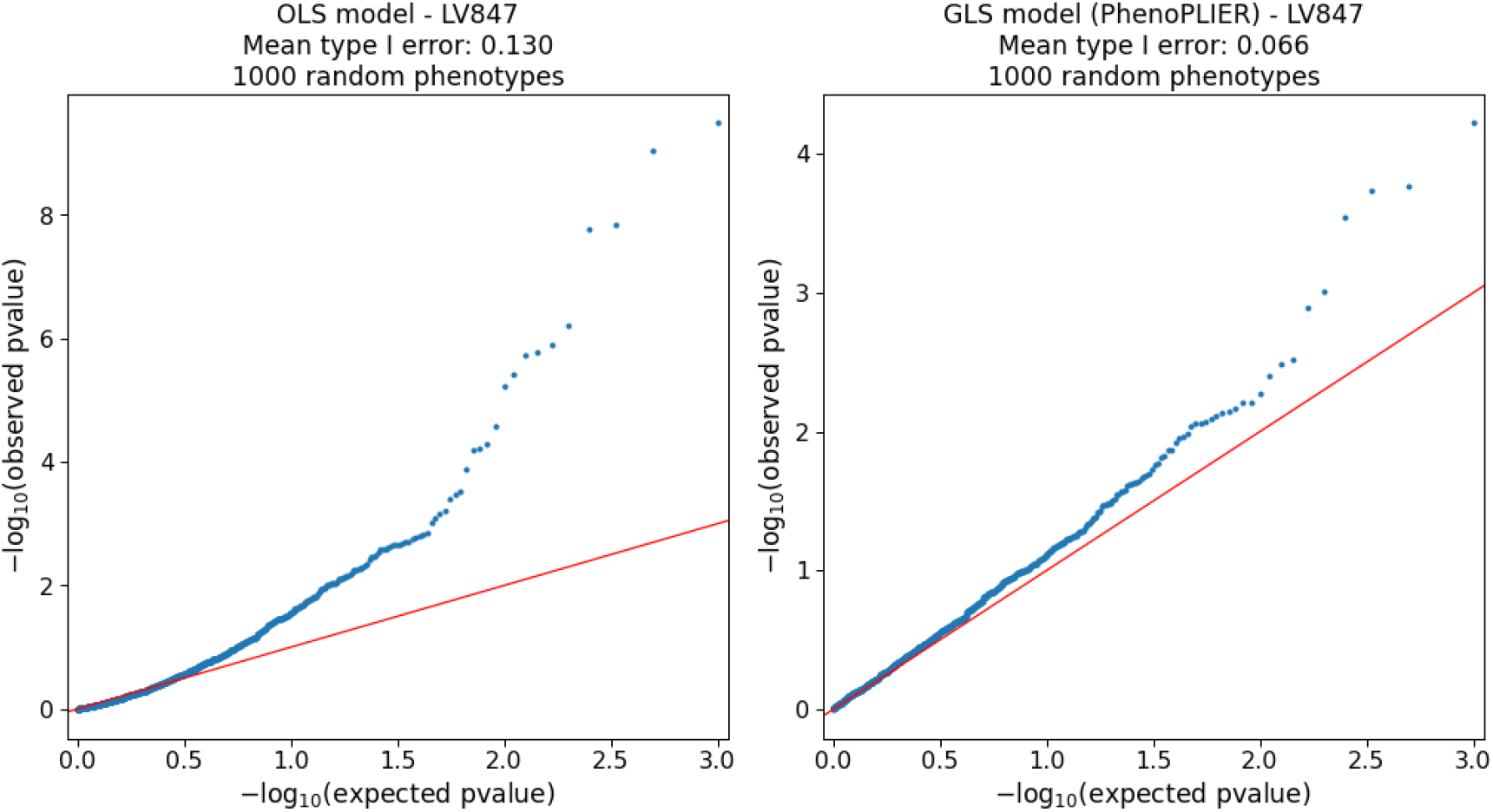
QQ-plots for LV847 on random phenotypes. Among the top 1% of genes in this LV, 15 are located in band 6p22.2, 5 in 6p22.1 and 2 in 15q26.1.

**Figure 11:**
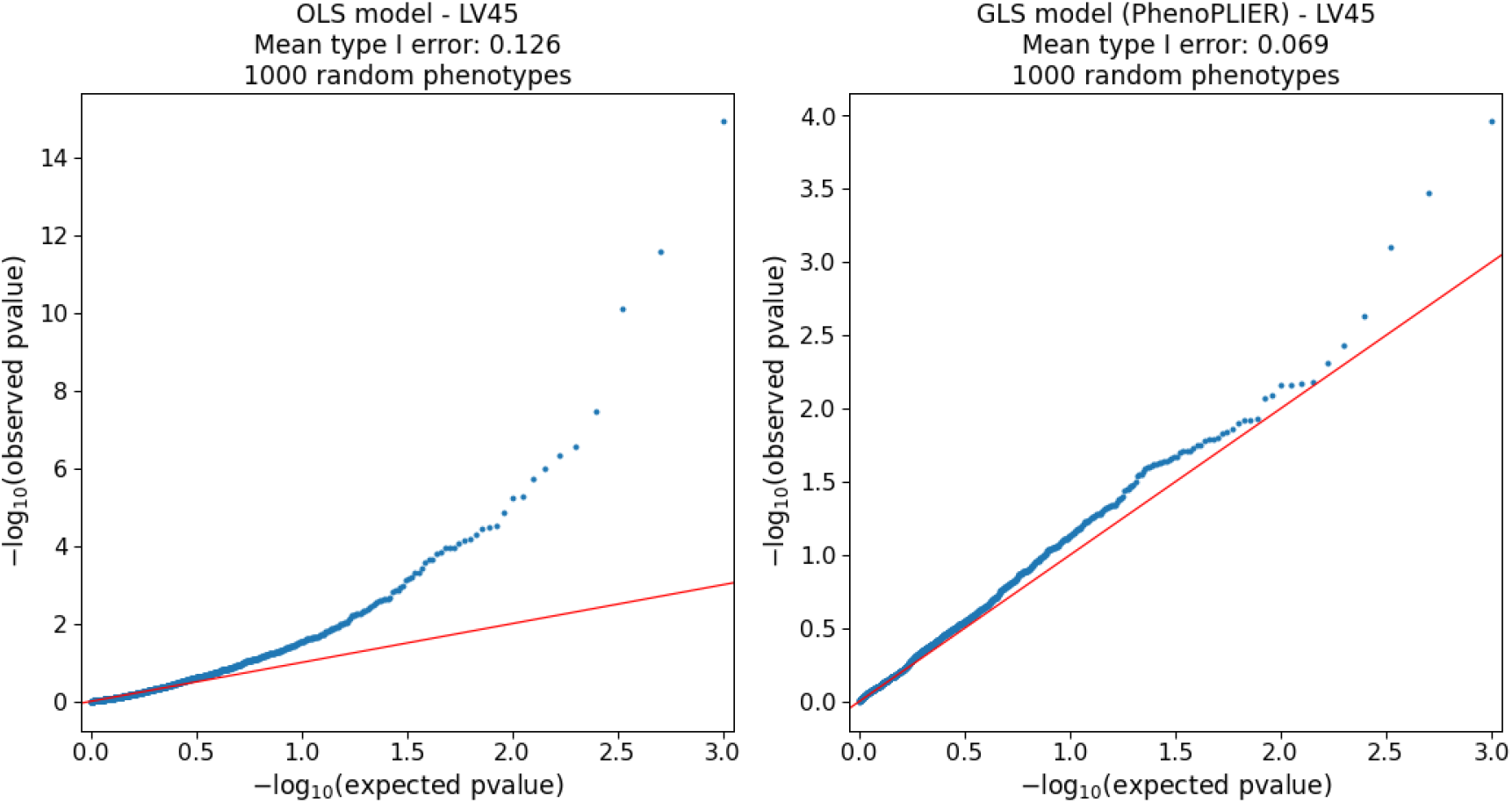
QQ-plots for LV45 on random phenotypes. Among the top 1% of genes in this LV, 12 are located in band 6p22.2, 6 in 6p22.1 and 3 in 1q23.3.

**Figure 12:**
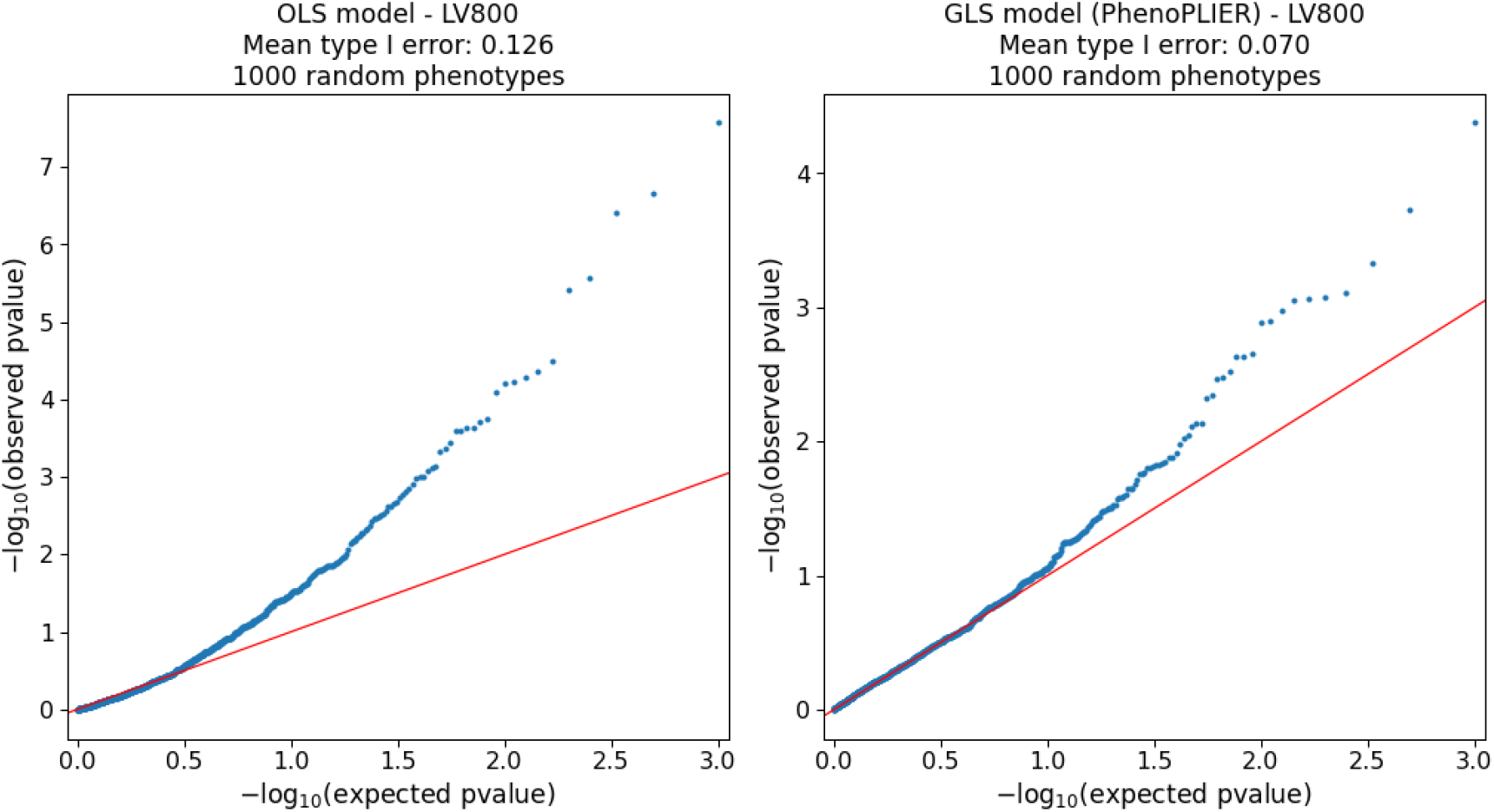
QQ-plots for LV800 on random phenotypes. Among the top 1% of genes in this LV, 16 are located in band 19q13.43, 9 in 19p13.2 and 9 in 19q13.31.

We also detected other LVs with higher-than-expected mean type I errors for both the GLS and OLS models, although they don’t have a relatively large number of adjacent genes at the top. One example is LV914, shown in Figure 13. Inflation in these LVs might be explained by inaccuracies in correlation estimates between the individual-level MultiXcan model and its summary-based version (see Methods). Therefore, we flagged those with a type I error rate larger than 0.07 (127 LVs) and excluded them from our main analyses.

**Figure 13:**
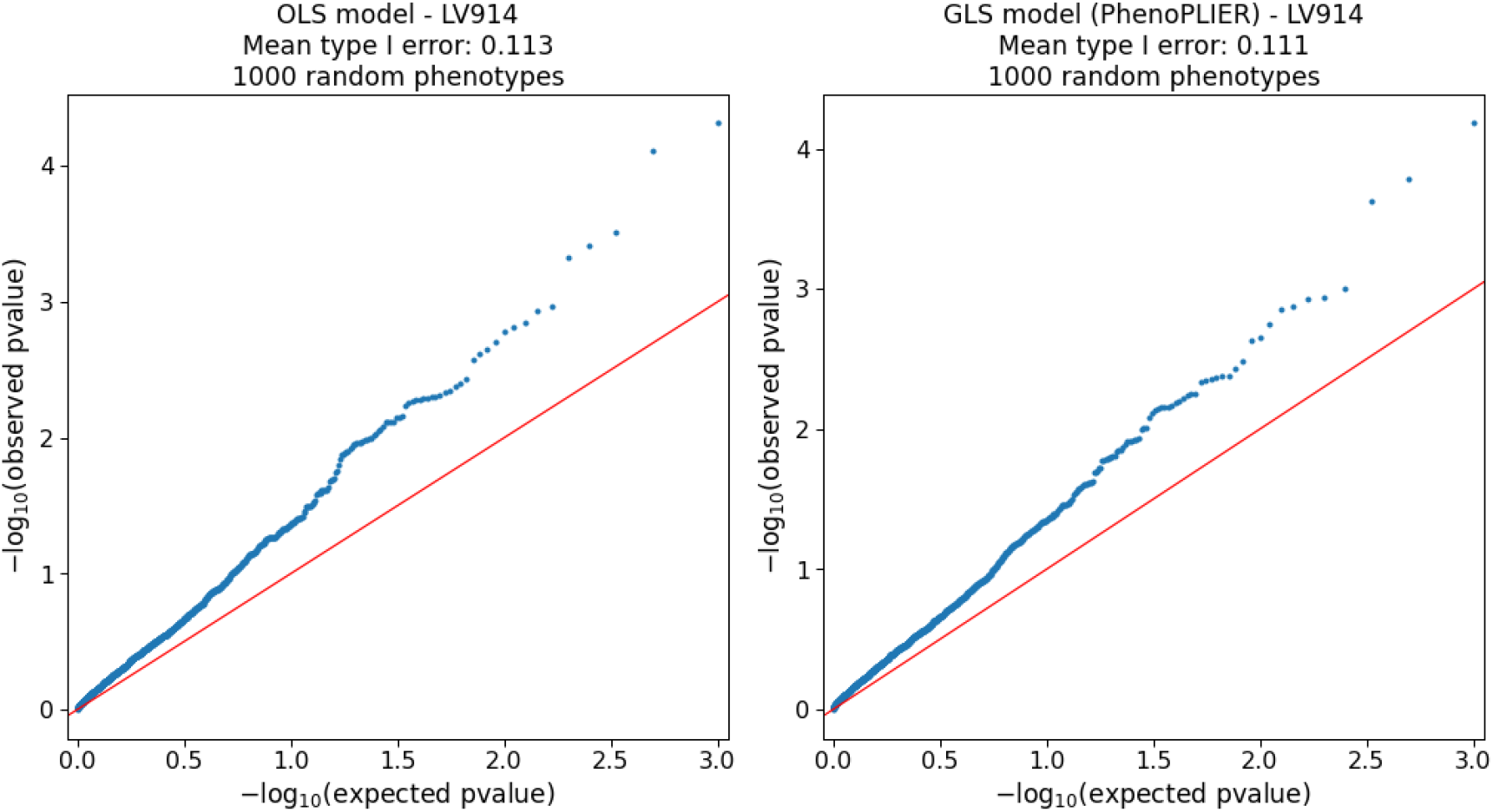
QQ-plots for LV914 on random phenotypes. Among the top 1% of genes in this LV, 2 are located in band 13q13.3, 2 in 7p15.2 and 2 in 19q13.2.

**Figure 14:**
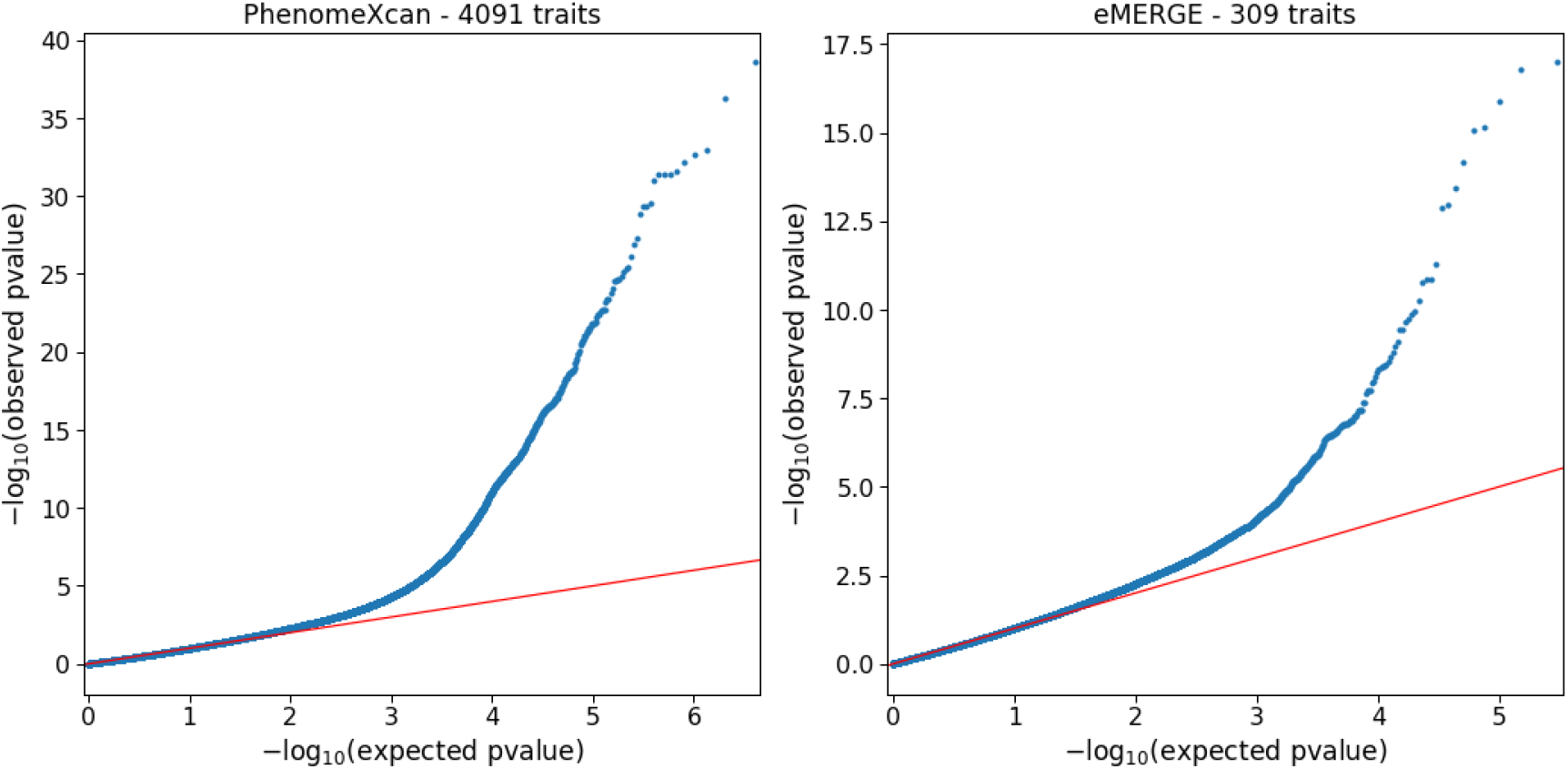
QQ-plots of LV-trait associations in real data. QQ-plot in PhenomeXcan (left, discovery cohort) across 4,091 traits and 987 LVs, and eMERGE (right, replication cohort) across 309 traits and 987 LVs.

#### LV-trait associations in real data

#### CRISPR-Cas9

##### Screening steps

**Figure S1:**
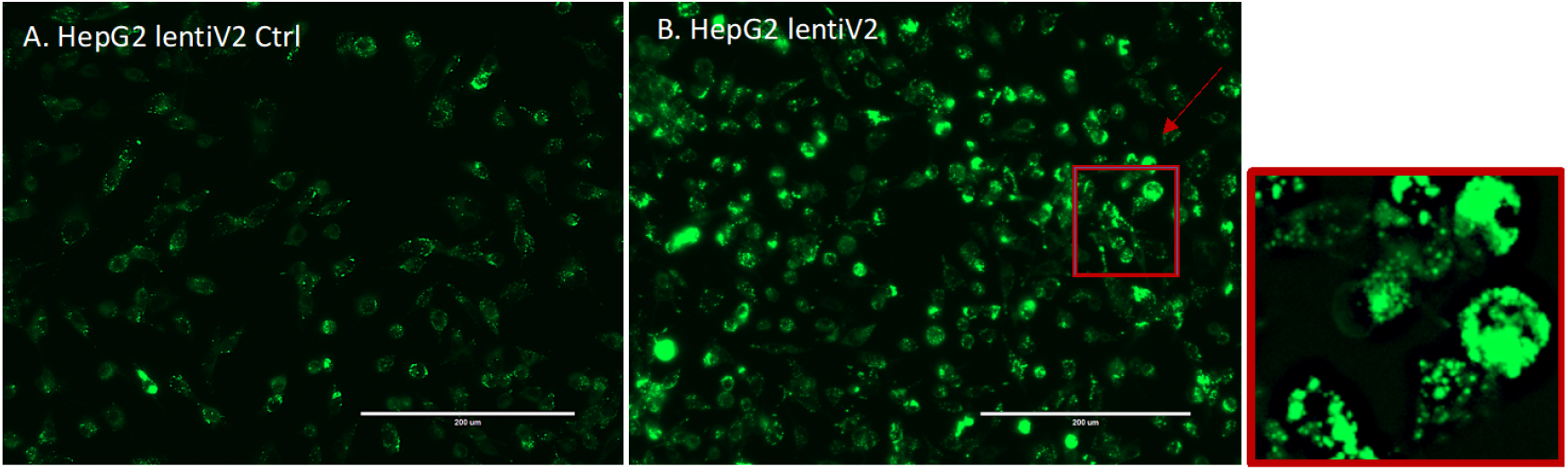
EVOS Fluorescent Microscope Image Capture. A. HepG2_lentiV2_Ctrl with no-viral transduction. B. HepG2_lentiV2 with viral transduction.

**Figure S2:**
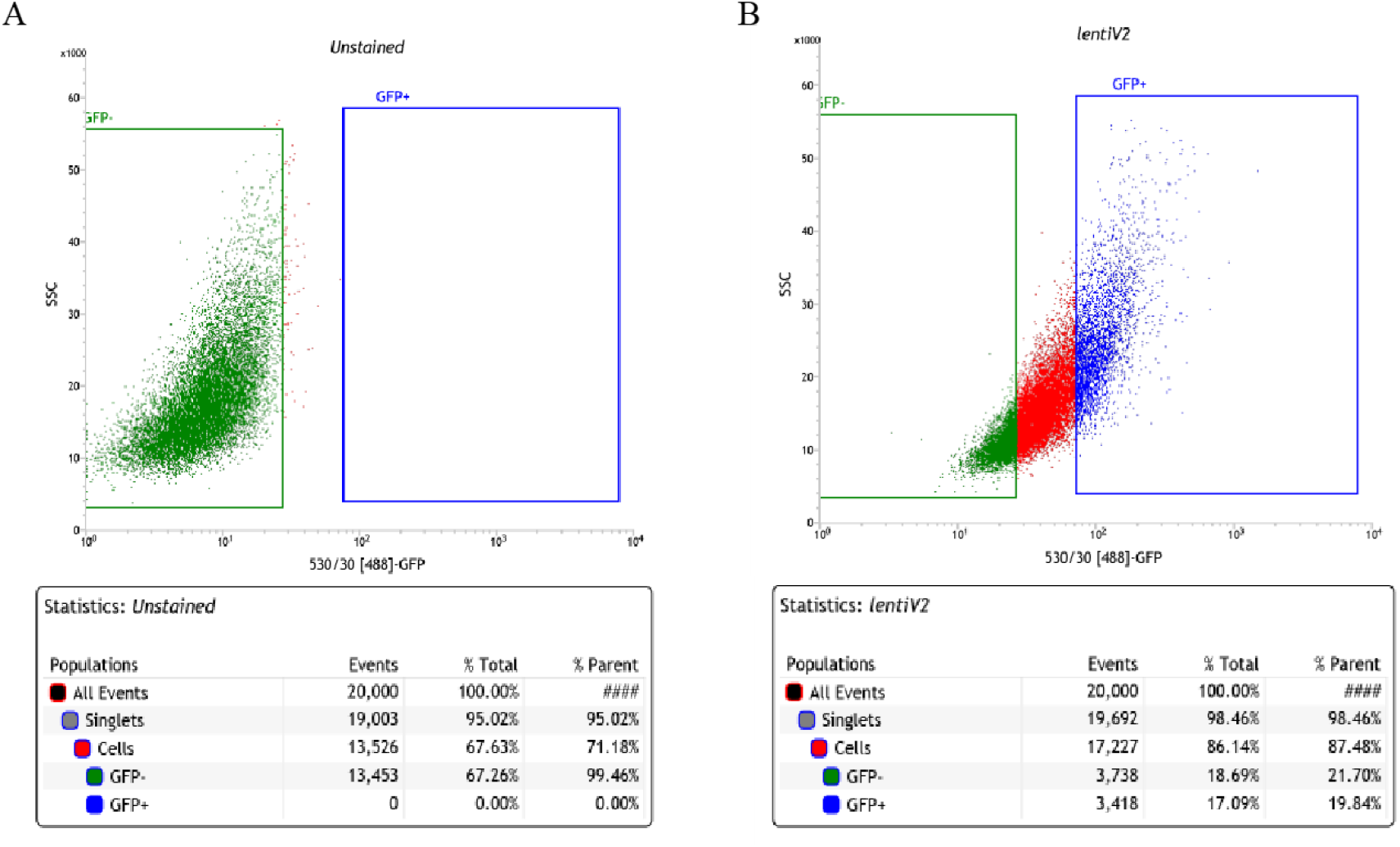
Fluorescence-Activated Cell Sorting Gate Setting. A. HepG2_UnStained WT. B. HepG2_lentiV2 with viral transduction.

**Figure S3:**
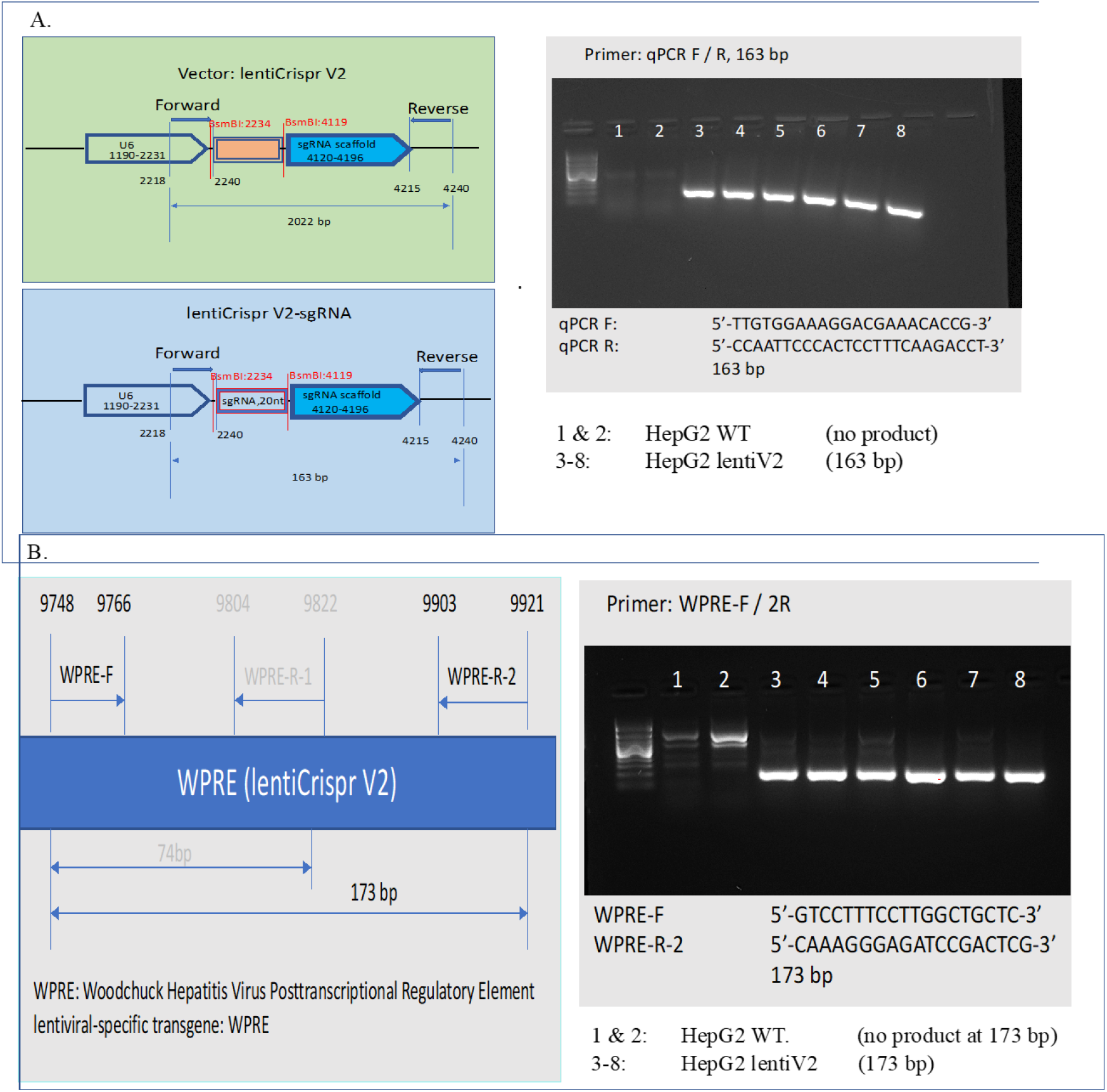
Verification of sgRNA cassette and lentiV2 transgene. A. 20nt sgRNA cassette was verified in lentiV2 transduced genomic DNA population, 163 bp PCR product obtained, while WT HepG2 didn’t possess the cassette, thus, no PCR product. B. lentiviral-specific transgene WPRE was verified in lentiV2 transduced genomic DNA population, while no transduced WT didn’t have the transgene, therefore, no 173 bp PCR product observed.

**Figure S4:**
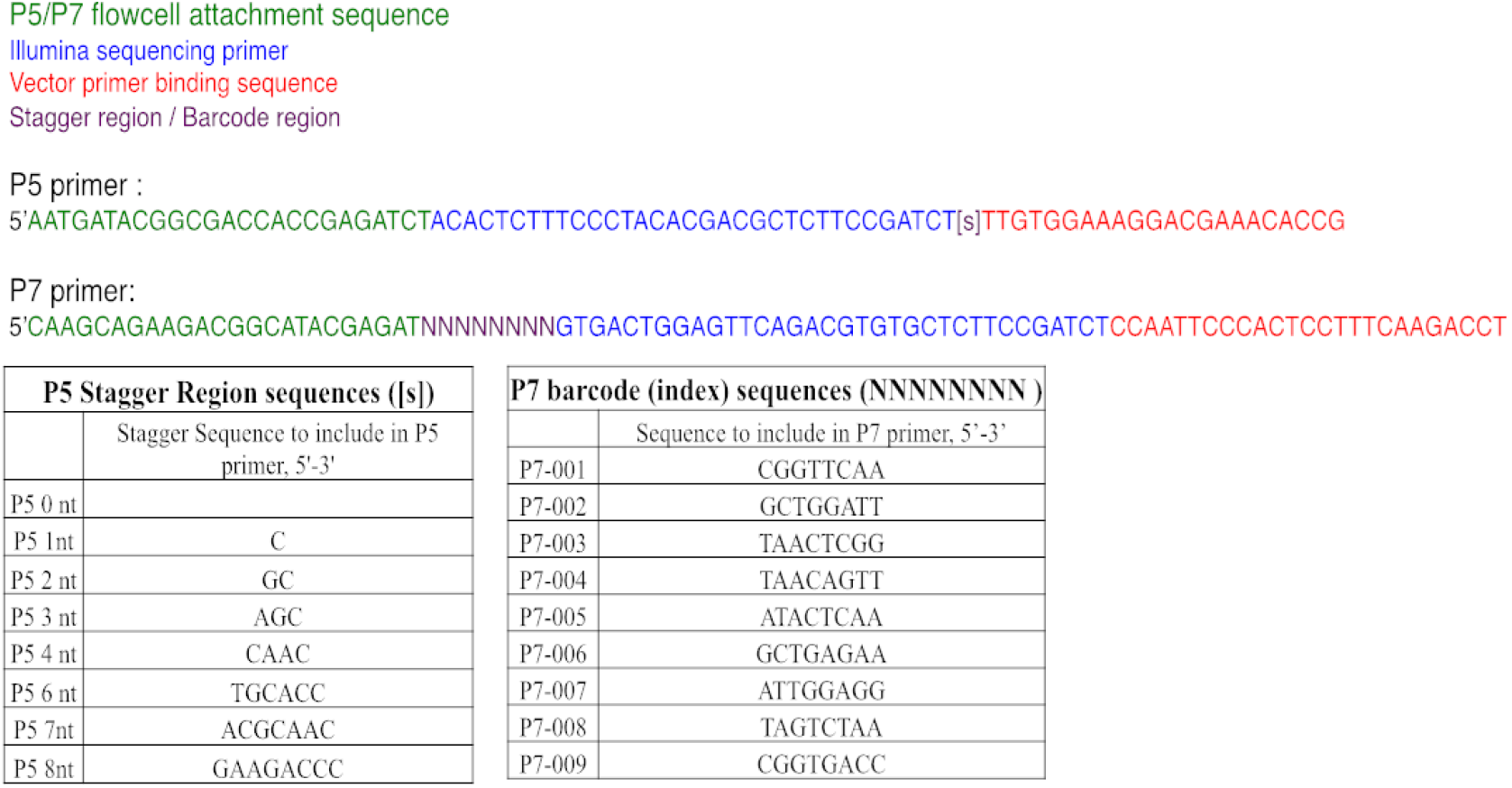

**Figure S5:**
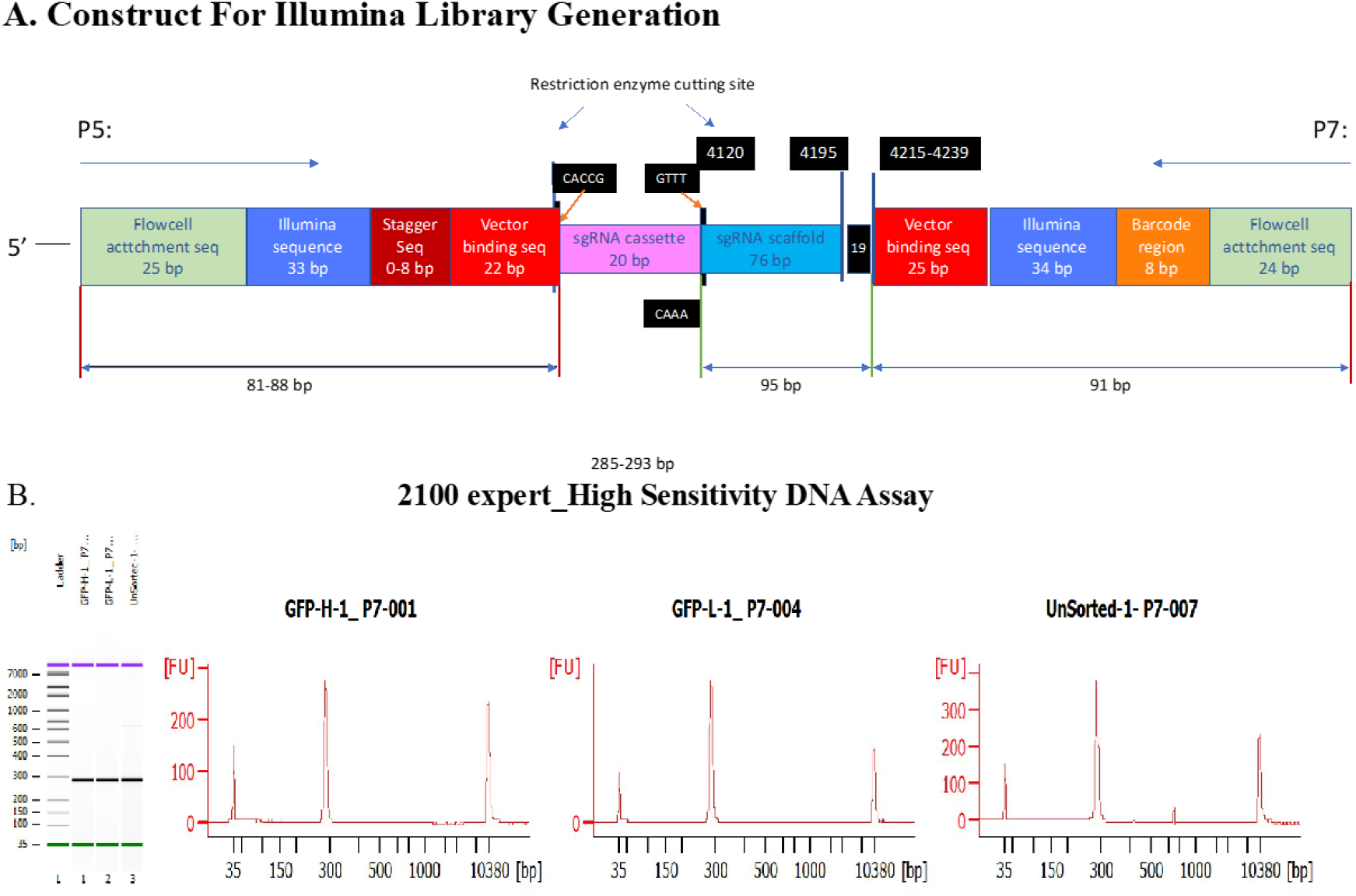
Illumina library generation. A. Construct for generating illumina libraries. B. Final illumina library from HS DNA —showed a single ~285bp peak was generated.

## Consensus clustering of traits

### Supplementary Note 2: Cluster analyses under the null hypothesis of no structure in the data

For our clustering pipeline, we simulated different escenarios where there is no structure in the input data matrix 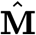 (gene-trait associations from PhenomeXcan projected into the latent gene expression representation). For this, we simulated two cases where any groupings of traits are removed: 1) the gene-trait association matrix **M** (from S-MultiXcan) does not have any meaningful structure to find groups of traits, while preserving the latent variables in **Z** from the MultiPLIER models; and 2) the latent variables in matrix **Z** does not have any meaningful structure to find groups of traits, while preserving the gene-trait association matrix **M**.

For the first scenario, we shuffled genes in **M** for each trait, and this randomized matrix was then projected into the latent space. For the second scenario, we projected matrix **M** into the latent space, and then shuffled LVs in 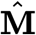 for each trait. For each of these scenarios, we ran exactly the same clustering pipeline we used for the real data (Methods), generating an ensemble of partitions that was later combined using the same consensus functions to derive the final partitions of traits. Finally, we computed 1) stability statistics on the ensemble partitions from different algorithms and 2) the agreement of the final consensus partition with the ensemble.

The results of this analysis (Figure 15) show that, under the two simulated null scenarios, the agreement of the consensus partitions with the ensemble is very close to zero. This means, as expected, that there is no consensus among ensemble partitions generated with different clustering algorithms and data representations. In contrast, using the real data, the consensus clustering approach finds trait pairs that are grouped together across the different members of the ensemble. The partitions above the 75th percentile were considered in the main analyses, and are shown in the clustering tree in Figure 6.

**Figure 15:**
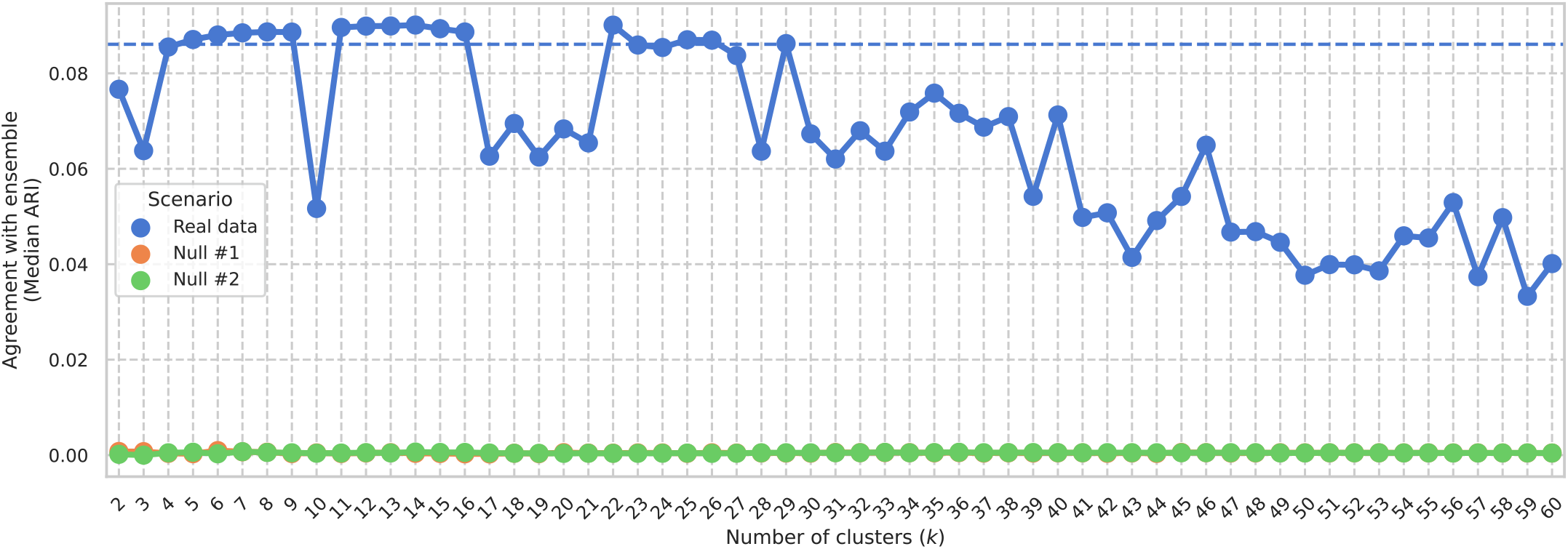
Agreement of consensus partitions with ensemble. A real and two simulated scenarios with no data structure are shown. For each scenario, one final consensus partition was derived for each ***k*** from 2 to 60 (***x***-axis) following our clustering pipeline. For each partition, the agreement with the corresponding ensemble was computed using the ARI (***y***-axis). For the real data scenario, partitions with an agreement above the 75th percentile (dashed line) were selected for follow-up analyses in the main text.

**Figure 16:**
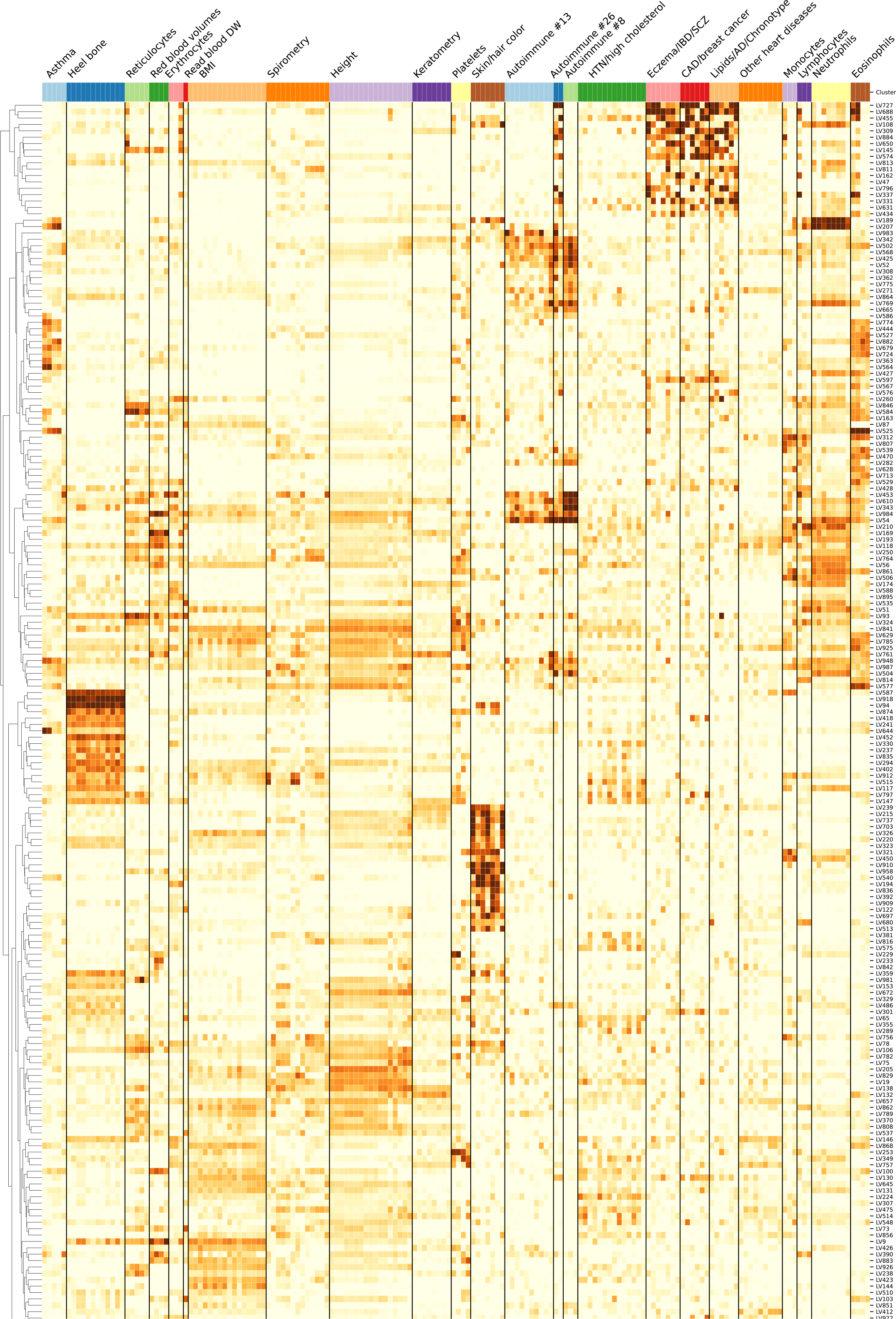
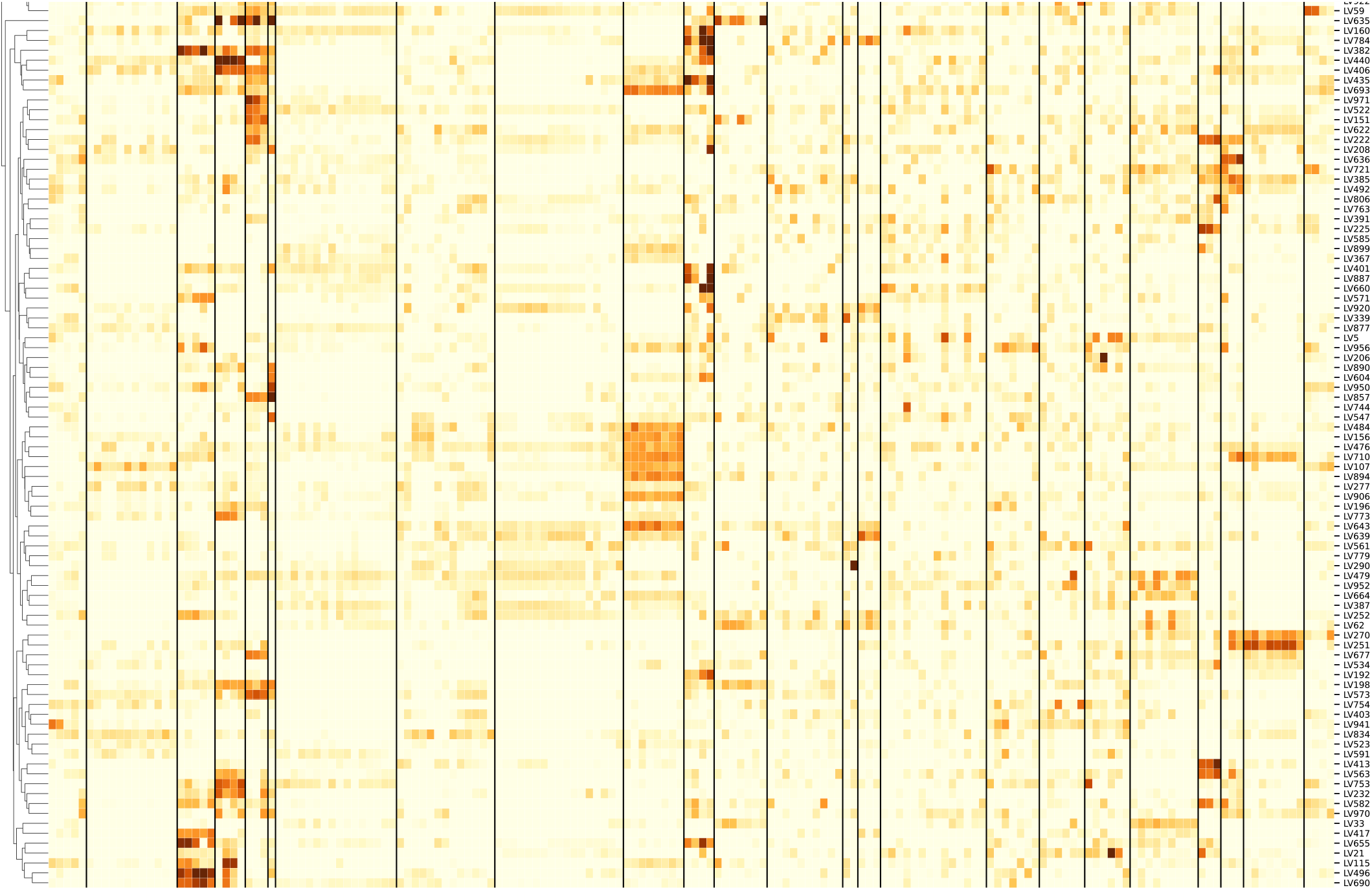
Cluster-specific and general transcriptional processes associated with disease using novel LVs. The plot shows a submatrix of 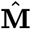 for the main trait clusters at ***k***=29, considering only LVs (rows) that are not aligned with any pathway. Standardized values from −6 (lighter color) to 21 (darker color).

**Figure 17:**
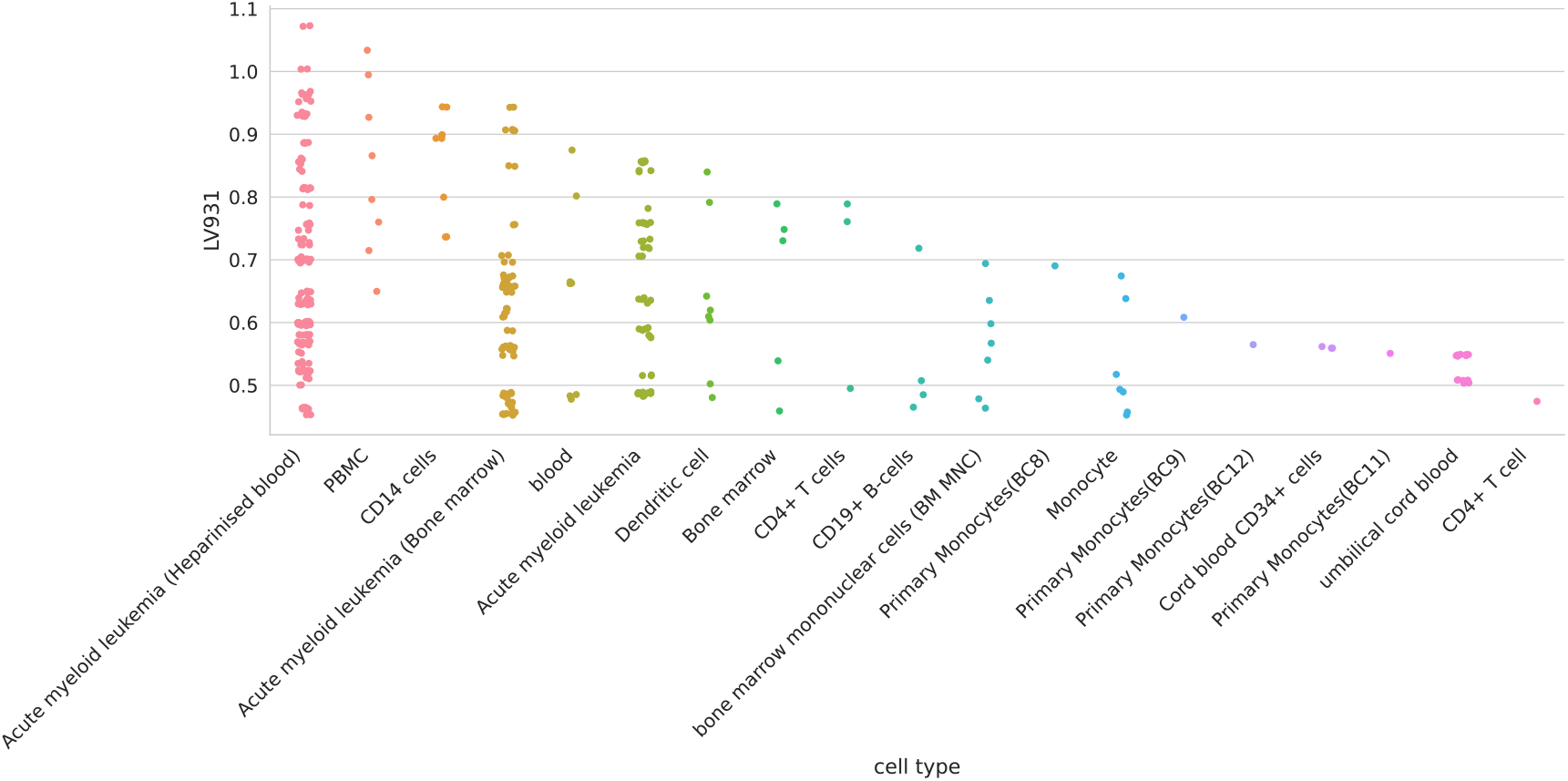
Cell types for LV931.

**Figure 18:**
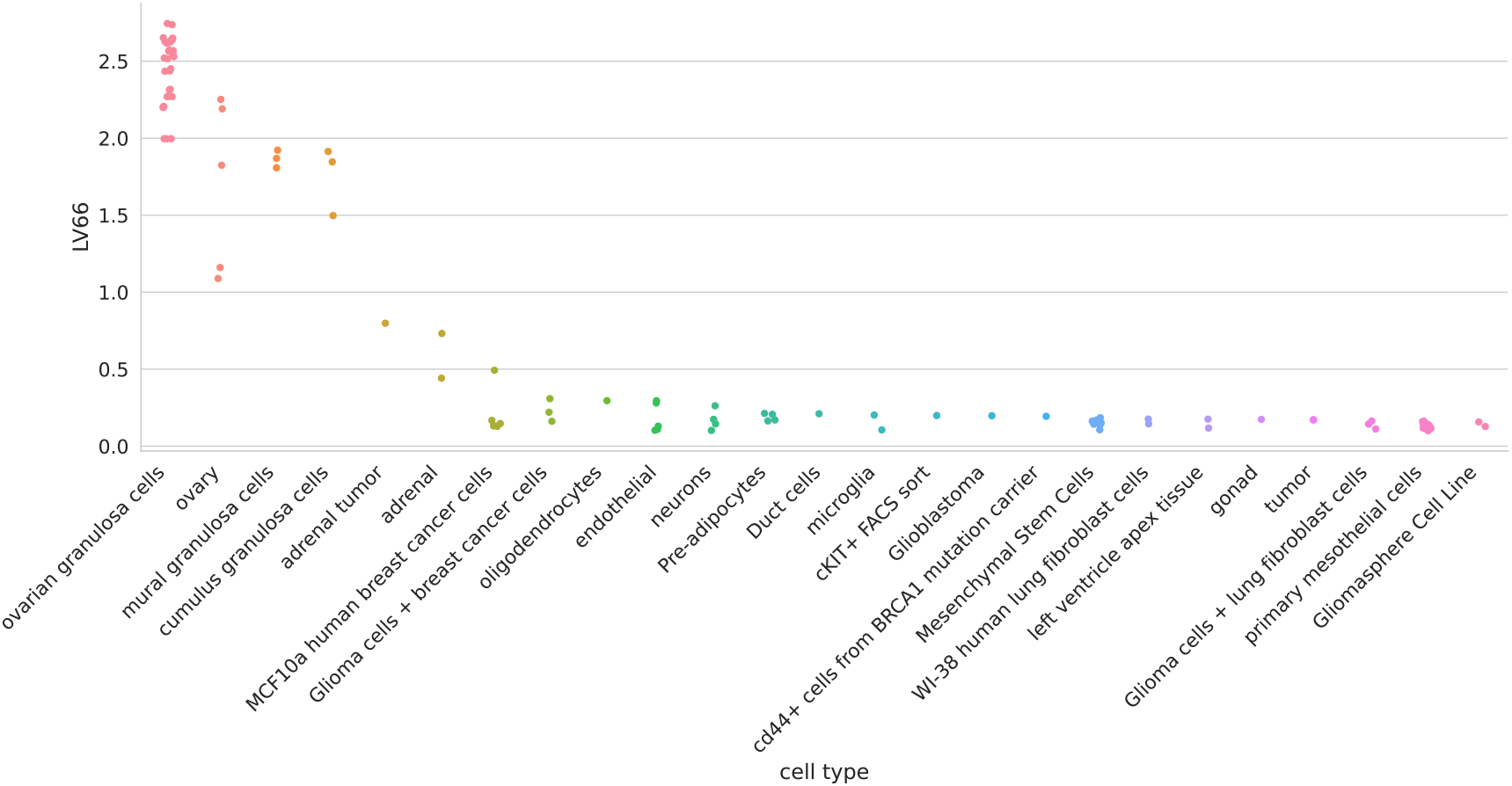
Cell types for LV66.

**Figure 19:**
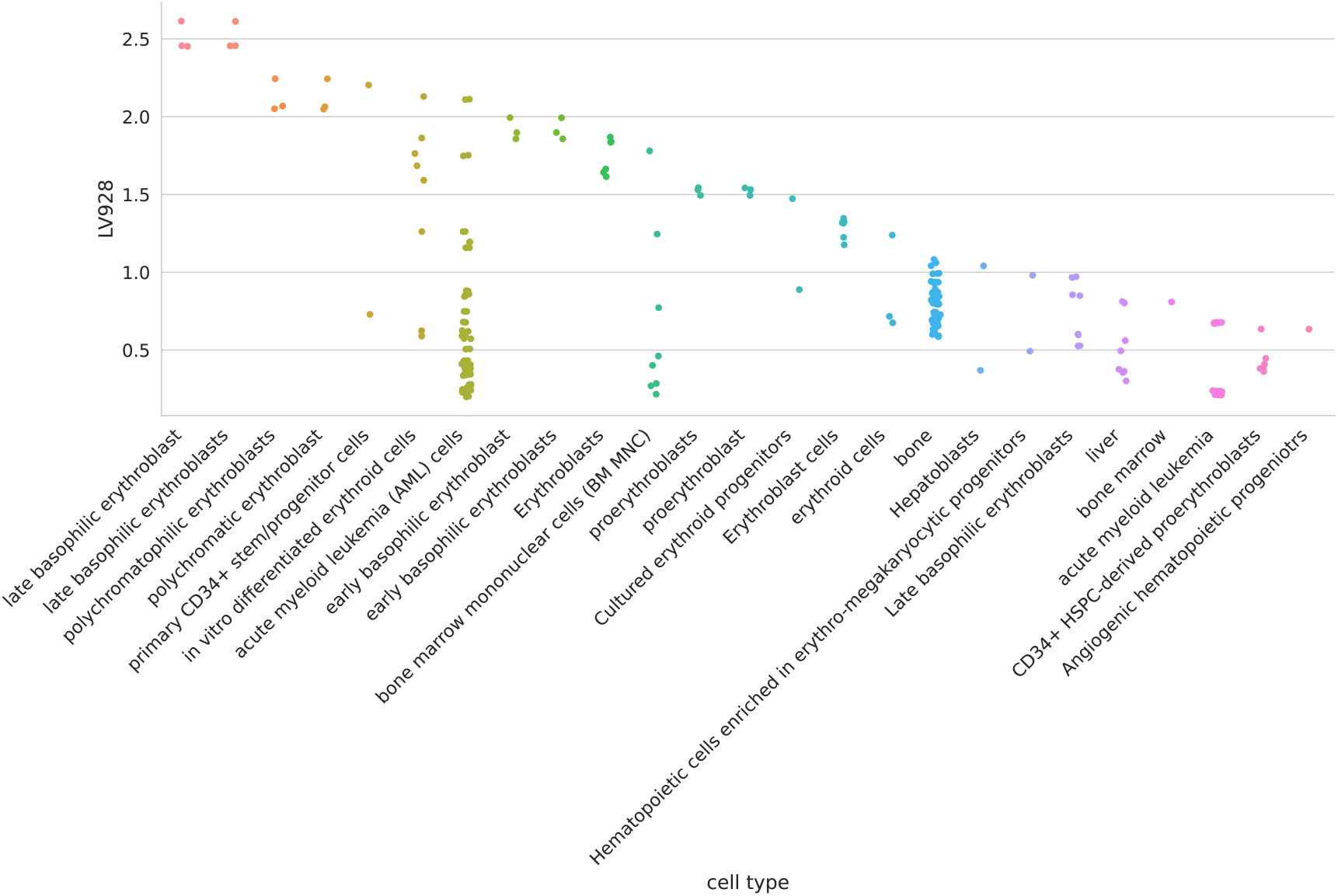
Cell types for LV928.

**Figure 20:**
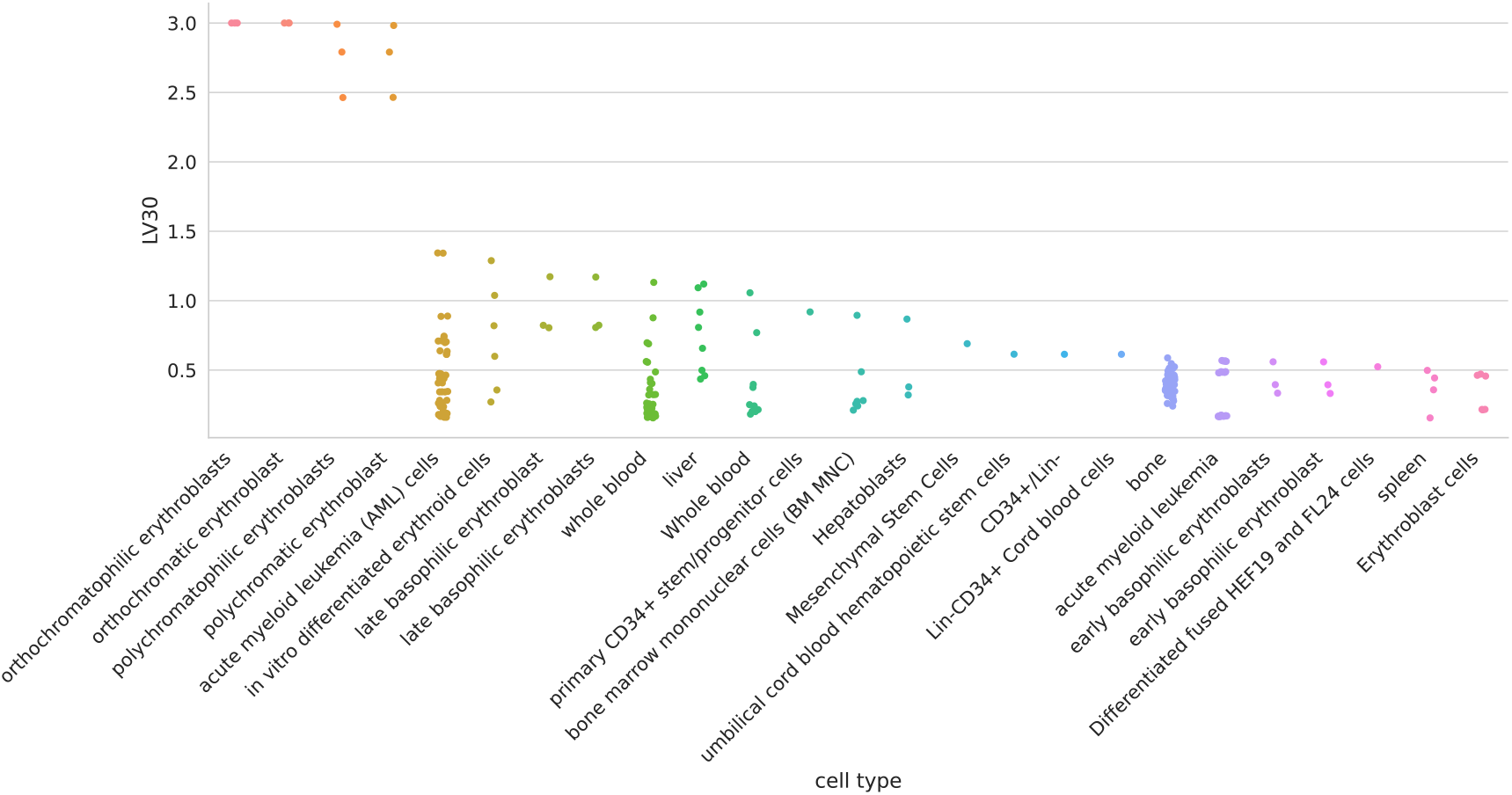
Cell types for LV30.

**Figure 21:**
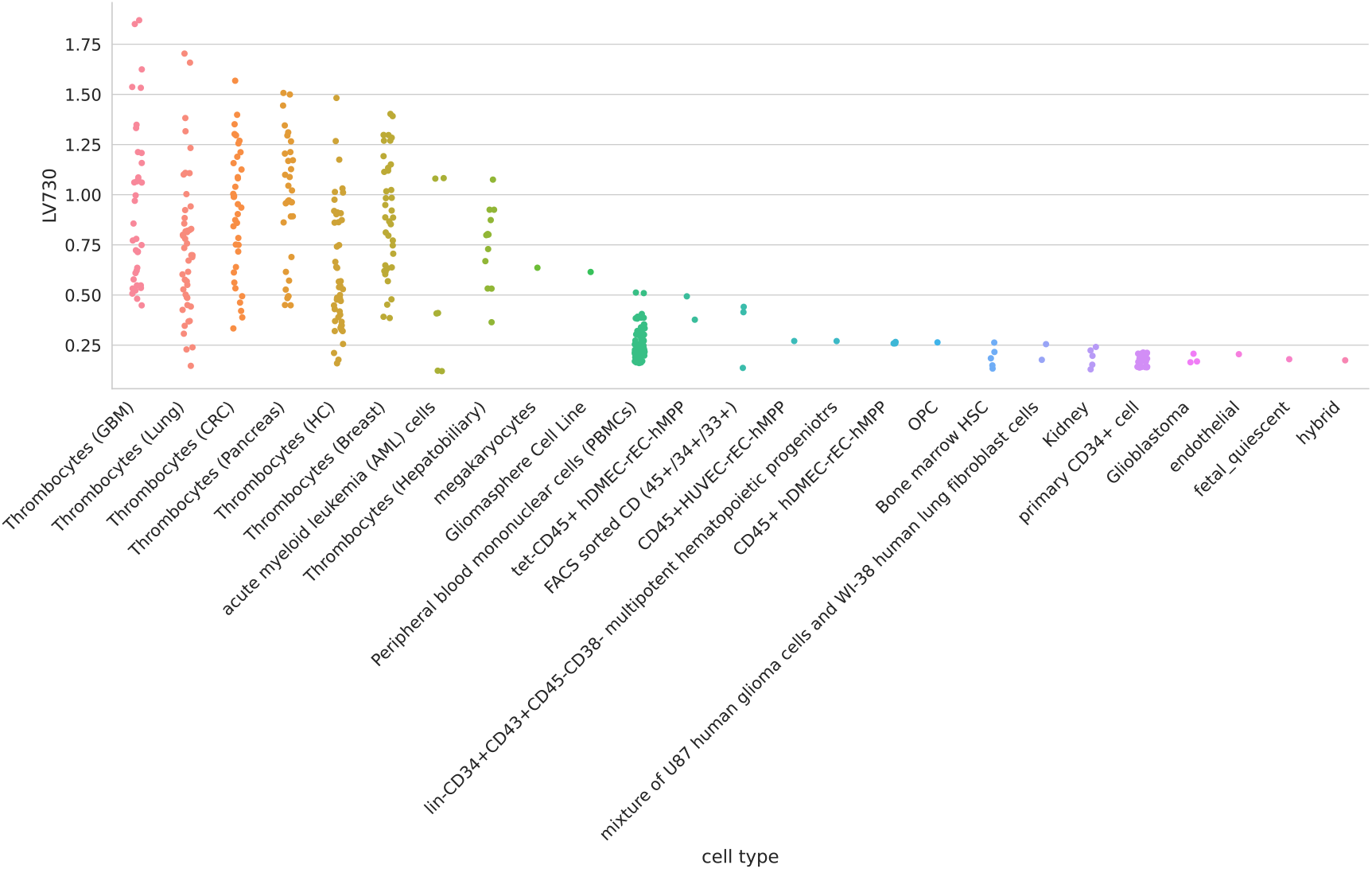
Cell types for LV730.

**Figure 22:**
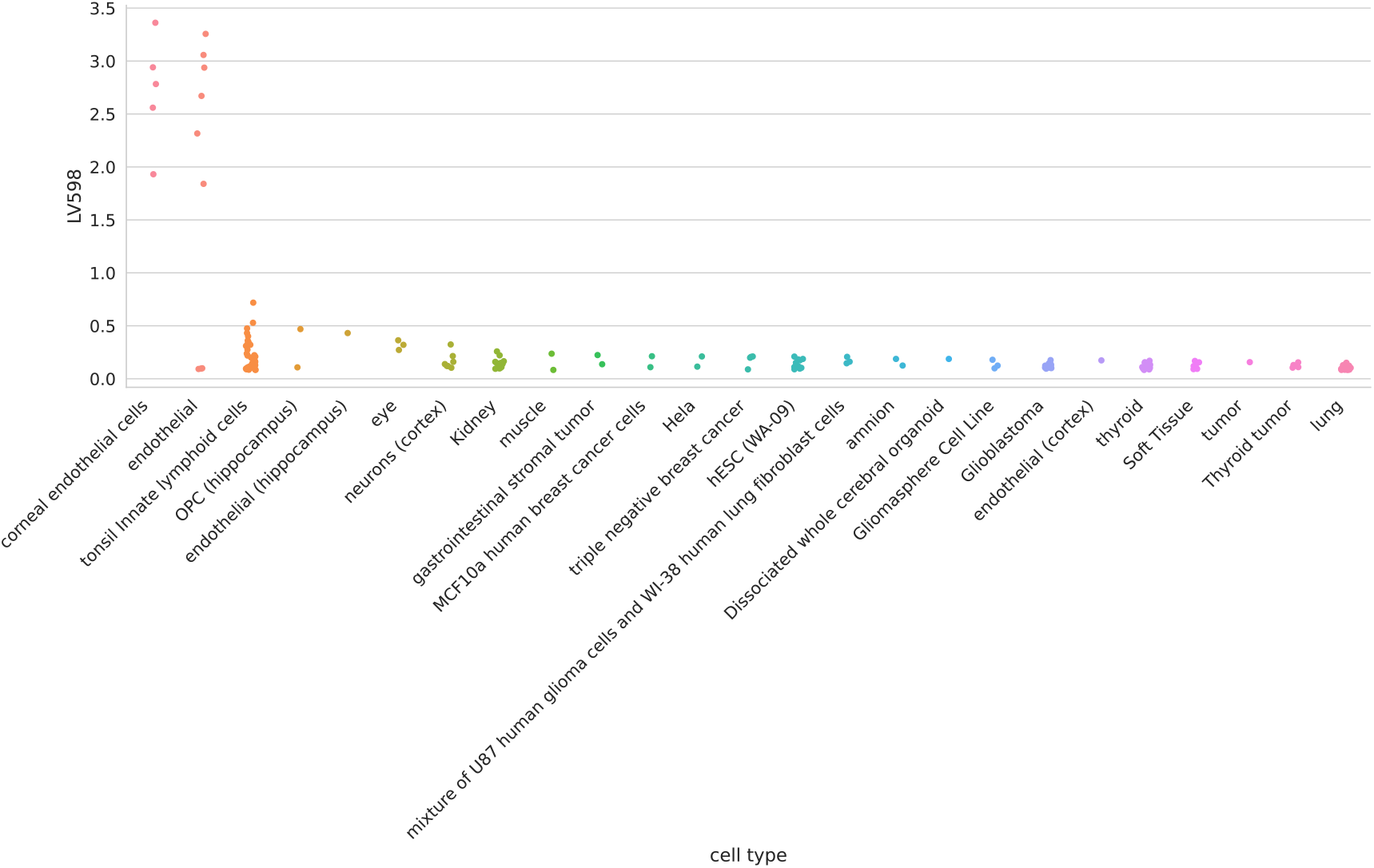
Cell types for LV598.

**Figure 23:**
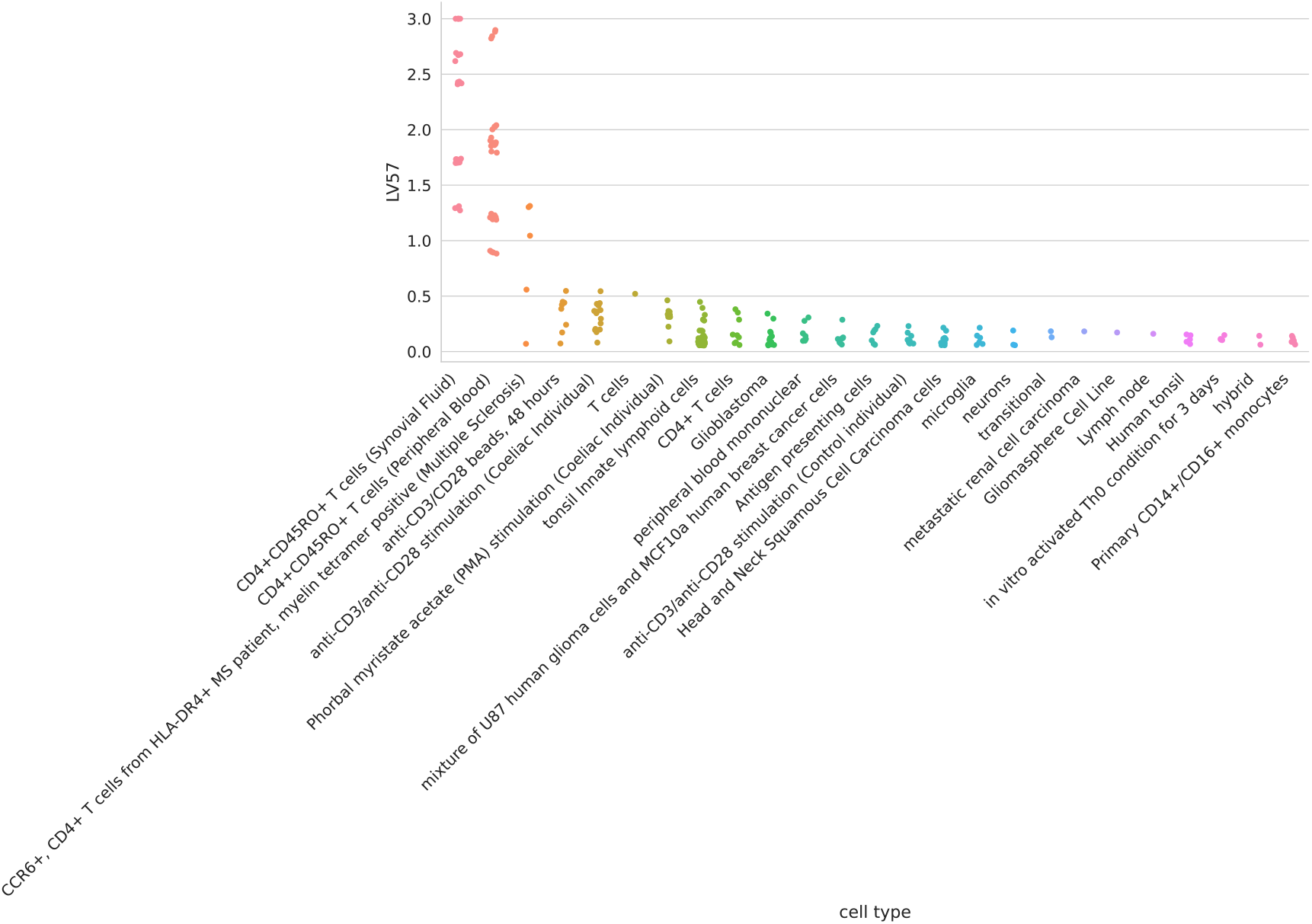
Cell types for LV57.

**Figure 24:**
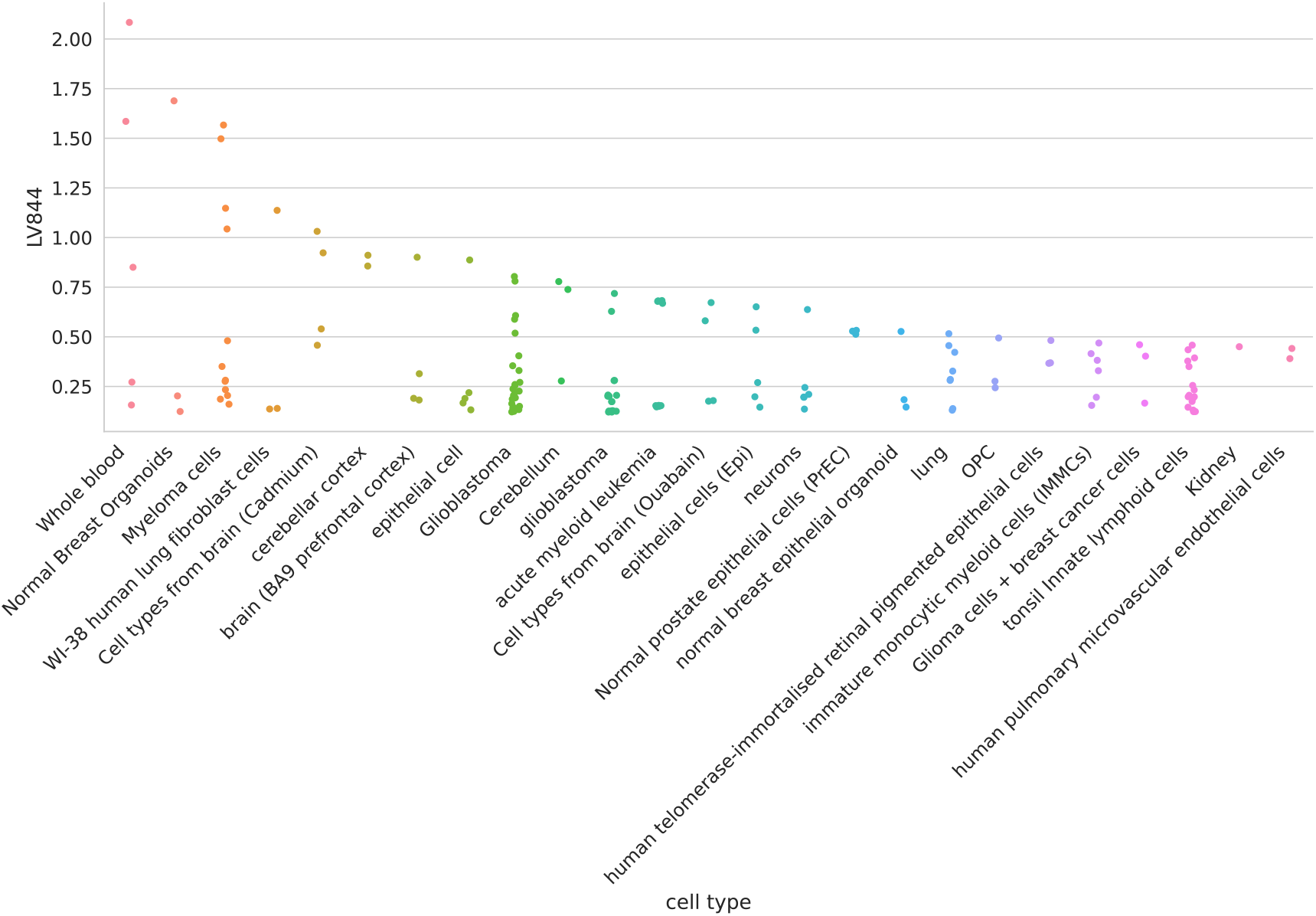
Cell types for LV844.

**Figure 25:**
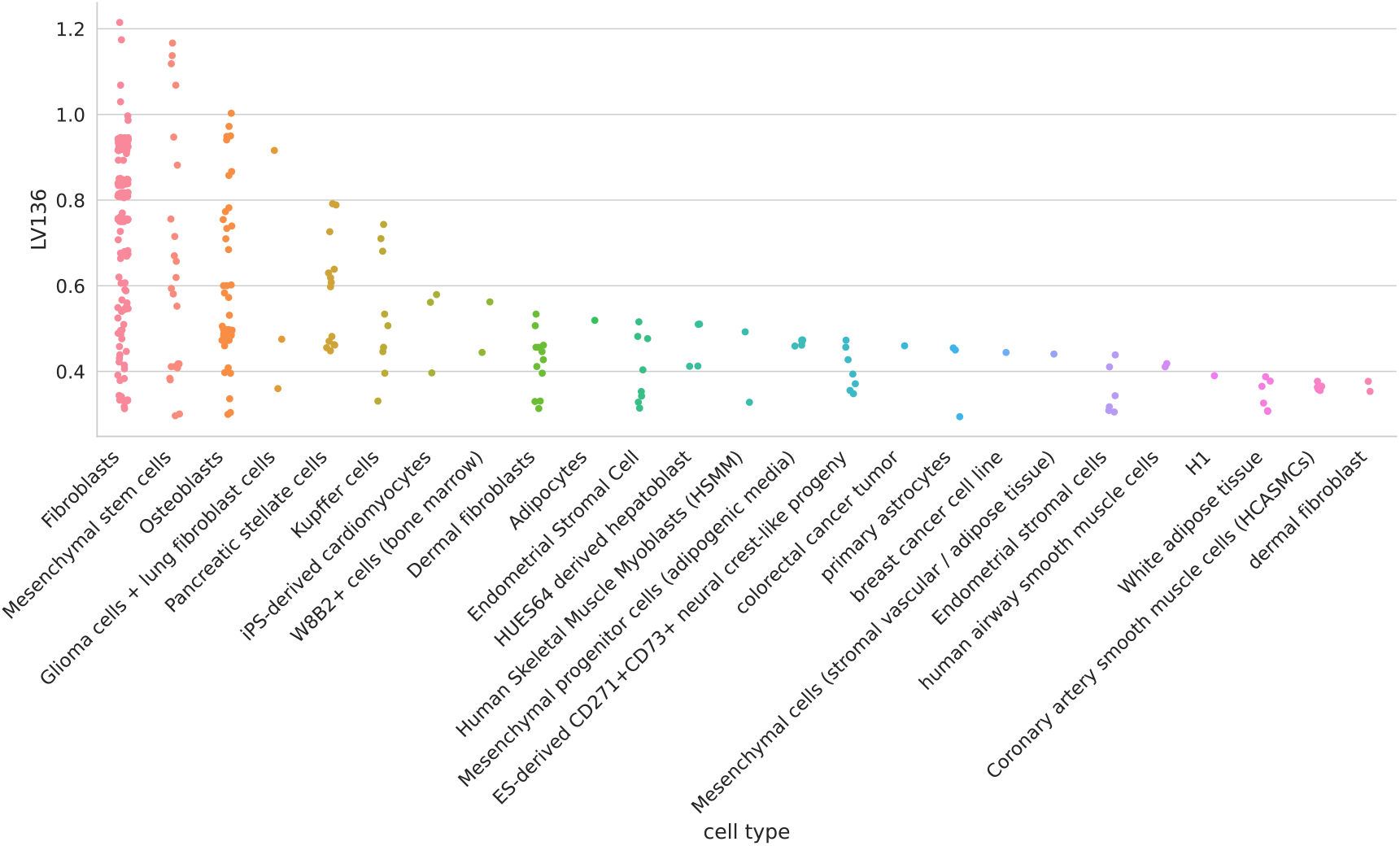
Cell types for LV136. Pulmonary microvascular endothelial cells were exposed to hypoxia for 24 hours or more [118];

**Figure 26:**
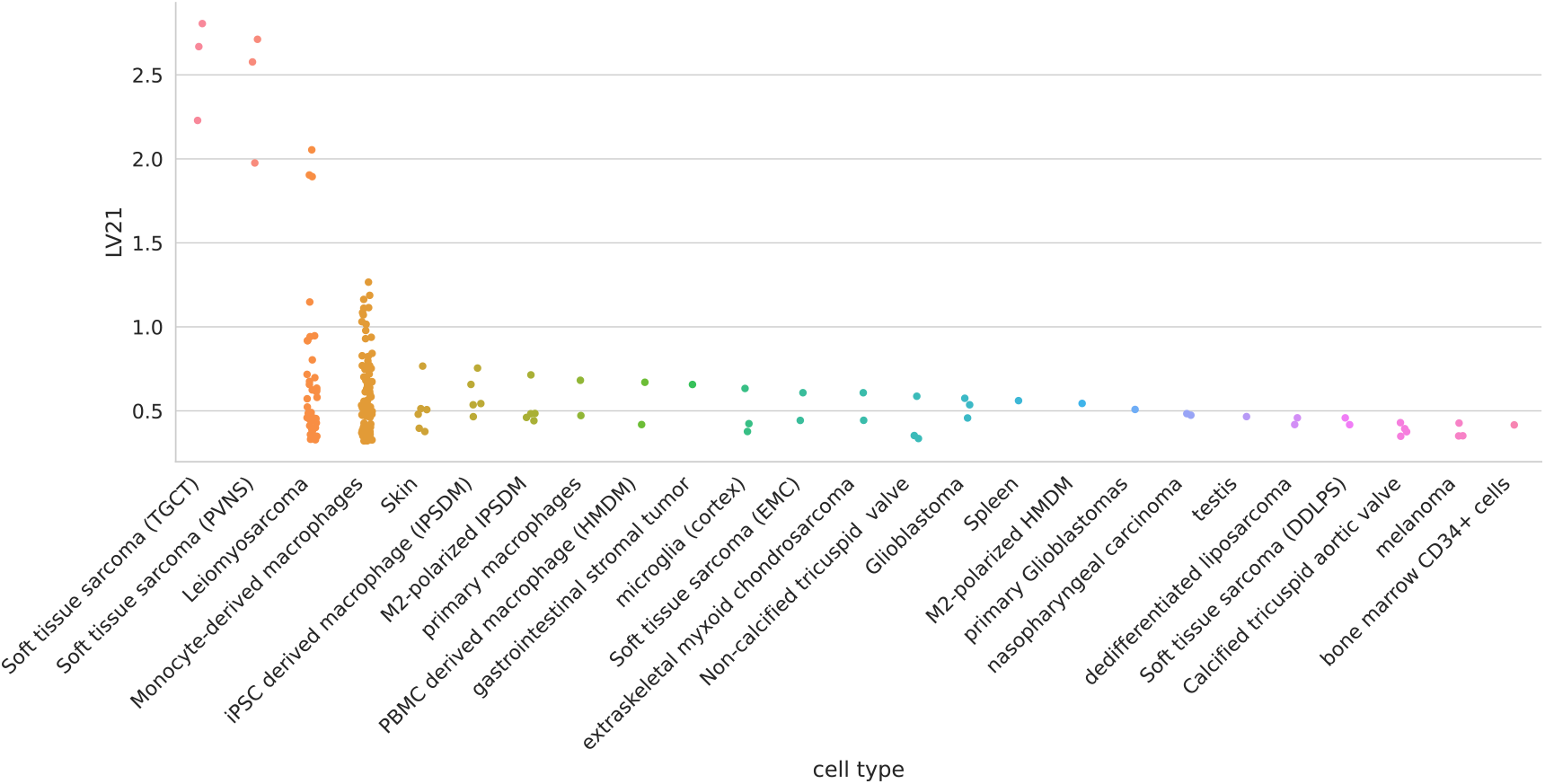
Cell types for LV21.

## References

1. A large-scale analysis of tissue-specific pathology and gene expression of human disease genes and complexes. Kasper Lage, Niclas Tue Hansen, EOlof Karlberg, Aron C Eklund, Francisco S Roque, Patricia K Donahoe, Zoltan Szallasi, Thomas Skøt Jensen, Søren Brunak Proceedings of the National Academy of Sciences of the United States of America (2008-12-22) https://www.ncbi.nlm.nih.gov/pubmed/19104045 DOI: 10.1073/pnas.0810772105 · PMID: 19104045 · PMCID: PMC2606902

2. Understanding multicellular function and disease with human tissue-specific network Casey S Greene, Arjun Krishnan, Aaron K Wong, Emanuela Ricciotti, Rene A Zelaya, Daniel S Himmelstein, Ran Zhang, Boris M Hartmann, Elena Zaslavsky, Stuart C Sealfon, … Olga G Troyanskaya Nature Genetics (2015-04-27) https://doi.org/f7dvkv DOI: 10.1038/ng.3259 · PMID: 25915600 · PMCID: PMC4828725

3. Conditional and interaction gene-set analysis reveals novel functional pathways for blood pressure Christiaan A de Leeuw, Sven Stringer, Ilona A Dekkers, Tom Heskes, Danielle Posthuma Nature Communications (2018-09-14) https://doi.org/gd6d85 DOI: 10.1038/s41467-018-06022-6 · PMID: 30218068 · PMCID: PMC6138636

4. The GTEx Consortium atlas of genetic regulatory effects across human tissues Science (2020-09-10) https://doi.org/ghbnhr DOI: 10.1126/science.aaz1776 · PMID: 32913098 · PMCID: PMC7737656

5. Index and biological spectrum of human DNase I hypersensitive sites Wouter Meuleman, Alexander Muratov, Eric Rynes, Jessica Halow, Kristen Lee, Daniel Bates, Morgan Diegel, Douglas Dunn, Fidencio Neri, Athanasios Teodosiadis, … John Stamatoyannopoulos Nature (2020-07-29) https://doi.org/gg6dhp DOI: 10.1038/s41586-020-2559-3 · PMID: 32728217 · PMCID: PMC7422677

6. Mechanisms of tissue and cell-type specificity in heritable traits and diseases Idan Hekselman, Esti Yeger-Lotem Nature Reviews Genetics (2020-01-08) https://doi.org/ggkx9v DOI: 10.1038/s41576-019-0200-9 · PMID: 31913361

7. The support of human genetic evidence for approved drug indications Matthew R Nelson, Hannah Tipney, Jeffery L Painter, Judong Shen, Paola Nicoletti, Yufeng Shen, Aris Floratos, Pak Chung Sham, Mulin Jun Li, Junwen Wang, … Philippe Sanseau Nature Genetics (2015-06-29) https://doi.org/f3mn52 DOI: 10.1038/ng.3314 · PMID: 26121088

8. Are drug targets with genetic support twice as likely to be approved? Revised estimates of the impact of genetic support for drug mechanisms on the probability of drug approval Emily A King, JWade Davis, Jacob F Degner PLOS Genetics (2019-12-12) https://doi.org/gg957r DOI: 10.1371/journal.pgen.1008489 · PMID: 31830040 · PMCID: PMC6907751

9. An integrated encyclopedia of DNA elements in the human genome Nature (2012-09) https://doi.org/bg9d DOI: 10.1038/nature11247 · PMID: 22955616 · PMCID: PMC3439153

10. Integrative analysis of 111 reference human epigenomes Anshul Kundaje, Wouter Meuleman, Jason Ernst, Misha Bilenky, Angela Yen, Alireza Heravi-Moussavi, Pouya Kheradpour, Zhizhuo Zhang, Jianrong Wang, … Manolis Kellis Nature (2015-02-18) https://doi.org/f62jpn DOI: 10.1038/nature14248 · PMID: 25693563 · PMCID: PMC4530010

11. An atlas of active enhancers across human cell types and tissues Robin Andersson, Claudia Gebhard, Irene Miguel-Escalada, Ilka Hoof, Jette Bornholdt, Mette Boyd, Yun Chen, Xiaobei Zhao, Christian Schmidl, … Albin Sandelin Nature (2014-03) https://doi.org/r35 DOI: 10.1038/nature12787 · PMID: 24670763 · PMCID: PMC5215096

12. Regulatory genomic circuitry of human disease loci by integrative epigenomics Carles A Boix, Benjamin T James, Yongjin P Park, Wouter Meuleman, Manolis Kellis Nature (2021-02-03) https://doi.org/ghzkhr DOI: 10.1038/s41586-020-03145-z · PMID: 33536621 · PMCID: PMC7875769

13. Heritability enrichment of specifically expressed genes identifies disease-relevant tissues and cell types Hilary K Finucane, Yakir A Reshef, Verneri Anttila, Kamil Slowikowski, Alexander Gusev, Andrea Byrnes, Steven Gazal, Po-Ru Loh, Caleb Lareau, Noam Shoresh, … Nature Genetics (2018-04) https://doi.org/gdfjqt DOI: 10.1038/s41588-018-0081-4 · PMID: 29632380 · PMCID: PMC5896795

14. The Post-GWAS Era: From Association to Function Michael D Gallagher, Alice S Chen-Plotkin The American Journal of Human Genetics (2018-05) https://doi.org/gdmftd DOI: 10.1016/j.ajhg.2018.04.002 · PMID: 29727686 · PMCID: PMC5986732

15. Biological interpretation of genome-wide association studies using predicted gene functions Tune H Pers, Juha M Karjalainen, Yingleong Chan, Harm-Jan Westra, Andrew R Wood, Jian Yang, Julian C Lui, Sailaja Vedantam, Stefan Gustafsson, … Lude Franke Nature Communications (2015-01-19) https://doi.org/f3mwhd DOI: 10.1038/ncomms6890 · PMID: 25597830 · PMCID: PMC4420238

16. Relaxed purifying selection and possibly high rate of adaptation in primate lineage-specific genes. James J Cai, Dmitri A Petrov Genome biology and evolution (2010-07-12) https://www.ncbi.nlm.nih.gov/pubmed/20624743 DOI: 10.1093/gbe/evq019 · PMID: 20624743 · PMCID: PMC2997544

17. Elevated rates of protein secretion, evolution, and disease among tissue-specific genes. Eitan E Winter, Leo Goodstadt, Chris P Ponting Genome research (2004-01) https://www.ncbi.nlm.nih.gov/pubmed/14707169 DOI: 10.1101/gr.1924004 · PMID: 14707169 · PMCID: PMC314278

18. A large-scale analysis of tissue-specific pathology and gene expression of human disease genes and complexes Kasper Lage, Niclas Tue Hansen, EOlof Karlberg, Aron C Eklund, Francisco S Roque, Patricia K Donahoe, Zoltan Szallasi, Thomas Skøt Jensen, Søren Brunak Proceedings of the National Academy of Sciences (2008-12-30) https://doi.org/d5qcv9 DOI: 10.1073/pnas.0810772105 · PMID: 19104045 · PMCID: PMC2606902

19. Reproducible RNA-seq analysis using recount2 Leonardo Collado-Torres, Abhinav Nellore, Kai Kammers, Shannon E Ellis, Margaret A Taub, Kasper D Hansen, Andrew E Jaffe, Ben Langmead, Jeffrey T Leek Nature Biotechnology (2017-04) https://doi.org/gf75hp DOI: 10.1038/nbt.3838 · PMID: 28398307 · PMCID: PMC6742427

20. Massive mining of publicly available RNA-seq data from human and mouse Alexander Lachmann, Denis Torre, Alexandra B Keenan, Kathleen M Jagodnik, Hoyjin J Lee, Lily Wang, Moshe C Silverstein, Avi Ma’ayan Nature Communications (2018-04-10) https://doi.org/gc92dr DOI: 10.1038/s41467-018-03751-6 · PMID: 29636450 · PMCID: PMC5893633

21. Identification of therapeutic targets from genetic association studies using hierarchical component analysis Hao-Chih Lee, Osamu Ichikawa, Benjamin S Glicksberg, Aparna A Divaraniya, Christine E Becker, Pankaj Agarwal, Joel T Dudley BioData Mining (2020-06-17) https://doi.org/gjp5pf DOI: 10.1186/s13040-020-00216-9 · PMID: 32565911 · PMCID: PMC7301559

22. Novel Variance-Component TWAS method for studying complex human diseases with applications to Alzheimer’s dementia Shizhen Tang, Aron S Buchman, Philip L De Jager, David A Bennett, Michael P Epstein, Jingjing Yang PLOS Genetics (2021-04-02) https://doi.org/gjpr3j DOI: 10.1371/journal.pgen.1009482 · PMID: 33798195 · PMCID: PMC8046351

23. Integrative approaches for large-scale transcriptome-wide association studies Alexander Gusev, Arthur Ko, Huwenbo Shi, Gaurav Bhatia, Wonil Chung, Brenda WJH Penninx, Rick Jansen, Eco JC de Geus, Dorret I Boomsma, Fred A Wright, … Bogdan Pasaniuc Nature Genetics (2016-02-08) https://doi.org/f3vf4p DOI: 10.1038/ng.3506 · PMID: 26854917 · PMCID: PMC4767558

24. Integrating predicted transcriptome from multiple tissues improves association detection Alvaro N Barbeira, Milton Pividori, Jiamao Zheng, Heather E Wheeler, Dan L Nicolae, Hae Kyung Im PLOS Genetics (2019-01-22) https://doi.org/ghs8vx DOI: 10.1371/journal.pgen.1007889 · PMID: 30668570 · PMCID: PMC6358100

25. A gene-based association method for mapping traits using reference transcriptome data Eric R Gamazon, Heather E Wheeler, Kaanan P Shah, Sahar V Mozaffari, Keston Aquino-Michaels, Robert J Carroll, Anne E Eyler, Joshua C Denny, Dan L Nicolae, Nancy J Cox, … Nature Genetics (2015-08-10) https://doi.org/f7p9zv DOI: 10.1038/ng.3367 · PMID: 26258848 · PMCID: PMC4552594

26. UTMOST, a single and cross-tissue TWAS (Transcriptome Wide Association Study), reveals new ASD (Autism Spectrum Disorder) associated genes. Cristina Rodriguez-Fontenla, Angel Carracedo Translational psychiatry (2021-04-30) https://www.ncbi.nlm.nih.gov/pubmed/33931583 DOI: 10.1038/s41398-021-01378-8 · PMID: 33931583 · PMCID: PMC8087708

27. Multi-ancestry gene-trait connection landscape using electronic health record (EHR) linked biobank data Binglan Li, Yogasudha Veturi, Anastasia Lucas, Yuki Bradford, Shefali S Verma, Anurag Verma, Joseph Park, Wei-Qi Wei, Qiping Feng, Bahram Namjou, … Marylyn D Ritchie Cold Spring Harbor Laboratory (2021-10-26) https://doi.org/gnbdnb DOI: 10.1101/2021.10.21.21265225

28. Shared and distinct genetic risk factors for childhood-onset and adult-onset asthma: genome-wide and transcriptome-wide studies. Milton Pividori, Nathan Schoettler, Dan L Nicolae, Carole Ober, Hae Kyung Im The Lancet. Respiratory medicine (2019-04-27) https://www.ncbi.nlm.nih.gov/pubmed/31036433 DOI: 10.1016/s2213-2600(19)30055-4 · PMID: 31036433 · PMCID: PMC6534440

29. Polygenic transcriptome risk scores (PTRS) can improve portability of polygenic risk scores across ancestries Yanyu Liang, Milton Pividori, Ani Manichaikul, Abraham A Palmer, Nancy J Cox, Heather E Wheeler, Hae Kyung Im Genome Biology (2022-01-13) https://doi.org/gqtdvn DOI: 10.1186/s13059-021-02591-w · PMID: 35027082 · PMCID: PMC8759285

30. Analysis of genome-wide association data highlights candidates for drug repositioning in psychiatry Hon-Cheong So, Carlos Kwan-Long Chau, Wan-To Chiu, Kin-Sang Ho, Cho-Pong Lo, Stephanie Ho-Yue Yim, Pak-Chung Sham Nature Neuroscience (2017-08-14) https://doi.org/gbrssh DOI: 10.1038/nn.4618 · PMID: 28805813

31. An Expanded View of Complex Traits: From Polygenic to Omnigenic Evan A Boyle, Yang I Li, Jonathan K Pritchard Cell (2017-06) https://doi.org/gcpgdz DOI: 10.1016/j.cell.2017.05.038 · PMID: 28622505 · PMCID: PMC5536862

32. A global overview of pleiotropy and genetic architecture in complex traits Kyoko Watanabe, Sven Stringer, Oleksandr Frei, Maša Umićević Mirkov, Christiaan de Leeuw, Tinca JC Polderman, Sophie van der Sluis, Ole A Andreassen, Benjamin M Neale, Danielle Posthuma Nature Genetics (2019-08-19) https://doi.org/ggr84r DOI: 10.1038/s41588-019-0481-0 · PMID: 31427789

33. Detection and interpretation of shared genetic inluences on 42 human traits Joseph K Pickrell, Tomaz Berisa, Jimmy Z Liu, Laure Ségurel, Joyce Y Tung, David A Hinds Nature Genetics (2016-05-16) https://doi.org/f8ssw4 DOI: 10.1038/ng.3570 · PMID: 27182965 · PMCID: PMC5207801

34. Trans Effects on Gene Expression Can Drive Omnigenic Inheritance Xuanyao Liu, Yang I Li, Jonathan K Pritchard Cell (2019-05) https://doi.org/gfz8bj DOI: 10.1016/j.cell.2019.04.014 · PMID: 31051098 · PMCID: PMC6553491

35. Architecture of the human regulatory network derived from ENCODE data. Mark B Gerstein, Anshul Kundaje, Manoj Hariharan, Stephen G Landt, Koon-Kiu Yan, Chao Cheng, Xinmeng Jasmine Mu, Ekta Khurana, Joel Rozowsky, Roger Alexander, … Michael Snyder Nature (2012-09-06) https://www.ncbi.nlm.nih.gov/pubmed/22955619 DOI: 10.1038/nature11245 · PMID: 22955619 · PMCID: PMC4154057

36. A narrow repertoire of transcriptional modules responsive to pyogenic bacteria is impaired in patients carrying loss-of-function mutations in MYD88 or IRAK4. Laia Alsina, Elisabeth Israelsson, Matthew C Altman, Kristen K Dang, Pegah Ghandil, Laura Israel, Horst von Bernuth, Nicole Baldwin, Huanying Qin, Zongbo Jin, … Damien Chaussabel Nature immunology (2014-10-26) https://www.ncbi.nlm.nih.gov/pubmed/25344726 DOI: 10.1038/ni.3028 · PMID: 25344726 · PMCID: PMC4281021

37. Cluster analysis and display of genome-wide expression patterns. MB Eisen, PT Spellman, PO Brown, D Botstein Proceedings of the National Academy of Sciences of the United States of America (1998-12-08) https://www.ncbi.nlm.nih.gov/pubmed/9843981 DOI: 10.1073/pnas.95.25.14863 · PMID: 9843981 · PMCID: PMC24541

38. Democratizing systems immunology with modular transcriptional repertoire analyses. Damien Chaussabel, Nicole Baldwin Nature reviews. Immunology (2014-04) https://www.ncbi.nlm.nih.gov/pubmed/24662387 DOI: 10.1038/nri3642 · PMID: 24662387 · PMCID: PMC4118927

39. How does gene expression clustering work? Patrik D’haeseleer Nature biotechnology (2005-12) https://www.ncbi.nlm.nih.gov/pubmed/16333293 DOI: 10.1038/nbt1205-1499 · PMID: 16333293

40. A comprehensive evaluation of module detection methods for gene expression data Wouter Saelens, Robrecht Cannoodt, Yvan Saeys Nature Communications (2018-03-15) https://doi.org/gc9x36 DOI: 10.1038/s41467-018-03424-4 · PMID: 29545622 · PMCID: PMC5854612

41. A modular analysis framework for blood genomics studies: application to systemic lupus erythematosus. Damien Chaussabel, Charles Quinn, Jing Shen, Pinakeen Patel, Casey Glaser, Nicole Baldwin, Dorothee Stichweh, Derek Blankenship, Lei Li, Indira Munagala, … Virginia Pascual Immunity (2008-07-18) https://www.ncbi.nlm.nih.gov/pubmed/18631455 DOI: 10.1016/j.immuni.2008.05.012 · PMID: 18631455 · PMCID: PMC2727981

42. PhenomeXcan: Mapping the genome to the phenome through the transcriptome Milton Pividori, Padma S Rajagopal, Alvaro Barbeira, Yanyu Liang, Owen Melia, Lisa Bastarache, YoSon Park, GTEx Consortium, Xiaoquan Wen, Hae K Im Science Advances (2020-09-11) https://doi.org/ghbvbf DOI: 10.1126/sciadv.aba2083 · PMID: 32917697

43. A Next Generation Connectivity Map: L1000 Platform and the First 1,000,000 Profiles Aravind Subramanian, Rajiv Narayan, Steven M Corsello, David D Peck, Ted E Natoli, Xiaodong Lu, Joshua Gould, John F Davis, Andrew A Tubelli, Jacob K Asiedu, … Todd R Golub Cell (2017-11) https://doi.org/cgwt DOI: 10.1016/j.cell.2017.10.049 · PMID: 29195078 · PMCID: PMC5990023

44. MultiPLIER: A Transfer Learning Framework for Transcriptomics Reveals Systemic Features of Rare Disease Jaclyn N Taroni, Peter C Grayson, Qiwen Hu, Sean Eddy, Matthias Kretzler, Peter A Merkel, Casey S Greene Cell Systems (2019-05) https://doi.org/gf75g5 DOI: 10.1016/j.cels.2019.04.003 · PMID: 31121115 · PMCID: PMC6538307

45. Pathway-level information extractor (PLIER) for gene expression data Weiguang Mao, Elena Zaslavsky, Boris M Hartmann, Stuart C Sealfon, Maria Chikina Nature Methods (2019-06-27) https://doi.org/gf75g6 DOI: 10.1038/s41592-019-0456-1 · PMID: 31249421 · PMCID: PMC7262669

46. The Electronic Medical Records and Genomics (eMERGE) Network: past, present, and future Omri Gottesman, Helena Kuivaniemi, Gerard Tromp, WAndrew Faucett, Rongling Li, Teri A Manolio, Saskia C Sanderson, Joseph Kannry, Randi Zinberg, Melissa A Basford, … Marc S Williams Genetics in Medicine (2013-10) https://doi.org/f5dwbt DOI: 10.1038/gim.2013.72 · PMID: 23743551 · PMCID: PMC3795928

47. The UK Biobank resource with deep phenotyping and genomic data Clare Bycroft, Colin Freeman, Desislava Petkova, Gavin Band, Lloyd T Elliott, Kevin Sharp, Allan Motyer, Damjan Vukcevic, Olivier Delaneau, Jared O’Connell, … Jonathan Marchini Nature (2018-10) https://doi.org/gfb7h2 DOI: 10.1038/s41586-018-0579-z · PMID: 30305743 · PMCID: PMC6786975

48. Finding function: evaluation methods for functional genomic data Chad L Myers, Daniel R Barrett, Matthew A Hibbs, Curtis Huttenhower, Olga G Troyanskaya BMC Genomics (2006-07-25) https://doi.org/fg6wnk DOI: 10.1186/1471-2164-7-187 · PMID: 16869964 · PMCID: PMC1560386

49. The CAFA challenge reports improved protein function prediction and new functional annotations for hundreds of genes through experimental screens Naihui Zhou, Yuxiang Jiang, Timothy R Bergquist, Alexandra J Lee, Balint Z Kacsoh, Alex W Crocker, Kimberley A Lewis, George Georghiou, Huy N Nguyen, Md Nafiz Hamid, … Iddo Friedberg Genome Biology (2019-11-19) https://doi.org/ggnxpz DOI: 10.1186/s13059-019-1835-8 · PMID: 31744546 · PMCID: PMC6864930

50. Estimating the population abundance of tissue-infiltrating immune and stromal cell populations using gene expression Etienne Becht, Nicolas A Giraldo, Laetitia Lacroix, Bénédicte Buttard, Nabila Elarouci, Florent Petitprez, Janick Selves, Pierre Laurent-Puig, Catherine Sautès-Fridman, Wolf H Fridman, Aurélien de Reyniès Genome Biology (2016-10-20) https://doi.org/f87sgf DOI: 10.1186/s13059-016-1070-5 · PMID: 27765066 · PMCID: PMC5073889

51. Fast gene set enrichment analysis Gennady Korotkevich, Vladimir Sukhov, Nikolay Budin, Boris Shpak, Maxim N Artyomov, Alexey Sergushichev Cold Spring Harbor Laboratory (2016-06-20) https://doi.org/gfpqhm DOI: 10.1101/060012

52. DrugBank 4.0: shedding new light on drug metabolism Vivian Law, Craig Knox, Yannick Djoumbou, Tim Jewison, An Chi Guo, Yifeng Liu, Adam Maciejewski, David Arndt, Michael Wilson, Vanessa Neveu, … David S Wishart Nucleic Acids Research (2013-11-06) https://doi.org/f3mn6d DOI: 10.1093/nar/gkt1068 · PMID: 24203711 · PMCID: PMC3965102

53. Systematic integration of biomedical knowledge prioritizes drugs for repurposing Daniel Scott Himmelstein, Antoine Lizee, Christine Hessler, Leo Brueggeman, Sabrina L Chen, Dexter Hadley, Ari Green, Pouya Khankhanian, Sergio E Baranzini eLife (2017-09-22) https://doi.org/cdfk DOI: 10.7554/elife.26726 · PMID: 28936969 · PMCID: PMC5640425

54. Dhimmel/Lincs V2.0: Refined Consensus Signatures From Lincs L1000 Daniel Himmelstein, Leo Brueggeman, Sergio Baranzini Zenodo (2016-03-08) https://doi.org/f3mqvr DOI: 10.5281/zenodo.47223

55. Computational Repositioning of the Anticonvulsant Topiramate for Inlammatory Bowel Disease Joel T Dudley, Marina Sirota, Mohan Shenoy, Reetesh K Pai, Silke Roedder, Annie P Chiang, Alex A Morgan, Minnie M Sarwal, Pankaj Jay Pasricha, Atul J Butte Science Translational Medicine (2011-08-17) https://doi.org/bmh5ts DOI: 10.1126/scitranslmed.3002648 · PMID: 21849664 · PMCID: PMC3479650

56. Discovery and Preclinical Validation of Drug Indications Using Compendia of Public Gene Expression Data Marina Sirota, Joel T Dudley, Jeewon Kim, Annie P Chiang, Alex A Morgan, Alejandro Sweet-Cordero, Julien Sage, Atul J Butte Science Translational Medicine (2011-08-17) https://doi.org/c3fwxv DOI: 10.1126/scitranslmed.3001318 · PMID: 21849665 · PMCID: PMC3502016

57. Dhimmel/Indications V1.0. Pharmacotherapydb: The Open Catalog Of Drug Therapies For Disease Daniel S Himmelstein, Pouya Khankhanian, Christine S Hessler, Ari J Green, Sergio E Baranzini Zenodo (2016-03-15) https://doi.org/f3mqwb DOI: 10.5281/zenodo.47664

58. Niacin in patients with low HDL cholesterol levels receiving intensive statin therapy., William E Boden, Jeffrey L Probstfield, Todd Anderson, Bernard R Chaitman, Patrice Desvignes-Nickens, Kent Koprowicz, Ruth McBride, Koon Teo, William Weintraub The New England journal of medicine (2011-11-15) https://www.ncbi.nlm.nih.gov/pubmed/22085343 DOI: 10.1056/nejmoa1107579 · PMID: 22085343

59. Effects of extended-release niacin with laropiprant in high-risk patients., Martin J Landray, Richard Haynes, Jemma C Hopewell, Sarah Parish, Theingi Aung, Joseph Tomson, Karl Wallendszus, Martin Craig, Lixin Jiang, … Jane Armitage The New England journal of medicine (2014-07-17) https://www.ncbi.nlm.nih.gov/pubmed/25014686 DOI: 10.1056/nejmoa1300955 · PMID: 25014686

60. Assessment of the Role of Niacin in Managing Cardiovascular Disease Outcomes: A Systematic Review and Meta-analysis. Elvira D’Andrea, Spencer P Hey, Cherie L Ramirez, Aaron S Kesselheim JAMA network open (2019-04-05) https://www.ncbi.nlm.nih.gov/pubmed/30977858 DOI: 10.1001/jamanetworkopen.2019.2224 · PMID: 30977858 · PMCID: PMC6481429

61. Mechanism of Action of Niacin Vaijinath S Kamanna, Moti L Kashyap The American Journal of Cardiology (2008-04) https://doi.org/c8zwdt DOI: 10.1016/j.amjcard.2008.02.029 · PMID: 18375237

62. Niacin: an old lipid drug in a new NAD+ dress Mario Romani, Dina Carina Hofer, Elena Katsyuba, Johan Auwerx Journal of Lipid Research (2019-04) https://doi.org/gjpjft DOI: 10.1194/jlr.s092007 · PMID: 30782960 · PMCID: PMC6446705

63. The therapeutic role of niacin in dyslipidemia management. William E Boden, Mandeep S Sidhu, Peter P Toth Journal of cardiovascular pharmacology and therapeutics (2013-12-20) https://www.ncbi.nlm.nih.gov/pubmed/24363242 DOI: 10.1177/1074248413514481 · PMID: 24363242

64. High-density lipoproteins in the prevention of cardiovascular disease: changing the paradigm. S Tuteja, DJ Rader Clinical pharmacology and therapeutics (2014-04-08) https://www.ncbi.nlm.nih.gov/pubmed/24713591 DOI: 10.1038/clpt.2014.79 · PMID: 24713591

65. The nicotinic acid receptor GPR109A (HM74A or PUMA-G) as a new therapeutic target S Offermanns Trends in Pharmacological Sciences (2006-07) https://doi.org/fgb4tr DOI: 10.1016/j.tips.2006.05.008 · PMID: 16766048

66. Langerhans Cells Release Prostaglandin D2 in Response to Nicotinic Acid Dominique Maciejewski-Lenoir, Jeremy G Richman, Yaron Hakak, Ibragim Gaidarov, Dominic P Behan, Daniel T Connolly Journal of Investigative Dermatology (2006-12) https://doi.org/dgxg75 DOI: 10.1038/sj.jid.5700586 · PMID: 17008871

67. Nicotinic acid inhibits progression of atherosclerosis in mice through its receptor GPR109A expressed by immune cells Martina Lukasova, Camille Malaval, Andreas Gille, Jukka Kero, Stefan Offermanns Journal of Clinical Investigation (2011-03-01) https://doi.org/cqftcq DOI: 10.1172/jci41651 · PMID: 21317532 · PMCID: PMC3048854

68. Role of HDL, ABCA1, and ABCG1 Transporters in Cholesterol Elux and Immune Responses Laurent Yvan-Charvet, Nan Wang, Alan R Tall Arteriosclerosis, Thrombosis, and Vascular Biology (2010-02) https://doi.org/ds23w6 DOI: 10.1161/atvbaha.108.179283 · PMID: 19797709 · PMCID: PMC2812788

69. Shared and organism-specific host responses to childhood diarrheal diseases revealed by whole blood transcript profiling Hannah A DeBerg, Mussaret B Zaidi, Matthew C Altman, Prasong Khaenam, Vivian H Gersuk, Freddy D Campos, Iza Perez-Martinez, Mario Meza-Segura, Damien Chaussabel, Jacques Banchereau, … Peter S Linsley PLOS ONE (2018-01-29) https://doi.org/gcwgcr DOI: 10.1371/journal.pone.0192082 · PMID: 29377961 · PMCID: PMC5788382

70. Copy Number Loss of the Interferon Gene Cluster in Melanomas Is Linked to Reduced T Cell Infiltrate and Poor Patient Prognosis Peter S Linsley, Cate Speake, Elizabeth Whalen, Damien Chaussabel PLoS ONE (2014-10-14) https://doi.org/gk9k8s DOI: 10.1371/journal.pone.0109760 · PMID: 25314013 · PMCID: PMC4196925

71. The Ro60 autoantigen binds endogenous retroelements and regulates inlammatory gene expression T Hung, GA Pratt, B Sundararaman, MJ Townsend, C Chaivorapol, T Bhangale, RR Graham, W Ortmann, LA Criswell, GW Yeo, TW Behrens Science (2015-10-23) https://doi.org/f7vs67 DOI: 10.1126/science.aac7442 · PMID: 26382853 · PMCID: PMC4691329

72. Homo sapiens (ID 258384) - BioProject - NCBI https://www.ncbi.nlm.nih.gov/bioproject/PRJNA258384

73. Identification of Genes Critical for Resistance to Infection by West Nile Virus Using RNA-Seq Analysis Feng Qian, Lisa Chung, Wei Zheng, Vincent Bruno, Roger Alexander, Zhong Wang, Xiaomei Wang, Sebastian Kurscheid, Hongyu Zhao, Erol Fikrig, … Ruth Montgomery Viruses (2013-07-08) https://doi.org/f49d7g DOI: 10.3390/v5071664 · PMID: 23881275 · PMCID: PMC3738954

74. Mycobacterial infection induces a specific human innate immune response John D Blischak, Ludovic Tailleux, Amy Mitrano, Luis B Barreiro, Yoav Gilad Scientific Reports (2015-11-20) https://doi.org/f7zk5c DOI: 10.1038/srep16882 · PMID: 26586179 · PMCID: PMC4653619

75. Niacin Inhibits Apoptosis and Rescues Premature Ovarian Failure Shufang Wang, Min Sun, Ling Yu, Yixuan Wang, Yuanqing Yao, Deqing Wang Cellular Physiology and Biochemistry (2018) https://doi.org/gfqvcq DOI: 10.1159/000495051 · PMID: 30415247

76. Chronic niacin administration ameliorates ovulation, histological changes in the ovary and adiponectin concentrations in a rat model of polycystic ovary syndrome Negin Asadi, Mahin Izadi, Ali Aflatounian, Mansour Esmaeili-Dehaj, Mohammad Ebrahim Rezvani, Zeinab Hafizi Reproduction, Fertility and Development (2021) https://doi.org/gjpjkt DOI: 10.1071/rd20306 · PMID: 33751926

77. Avoiding common pitfalls when clustering biological data Tom Ronan, Zhijie Qi, Kristen M Naegle Science Signaling (2016-06-14) https://doi.org/gcvjr6 DOI: 10.1126/scisignal.aad1932 · PMID: 27303057

78. Cluster Ensembles – A Knowledge Reuse Framework for Combining Multiple Partitions Alexander Strehl, Ghosh Joydeep Journal of Machine Learning Research (2002) https://www.jmlr.org/papers/v3/strehl02a.html

79. Combining multiple clusterings using evidence accumulation Ana LN Fred, Anil K Jain IEEE Transactions on Pattern Analysis and Machine Intelligence (2005-06) https://doi.org/bsknv6 DOI: 10.1109/tpami.2005.113 · PMID: 15943417

80. Clustering trees: a visualization for evaluating clusterings at multiple resolutions Luke Zappia, Alicia Oshlack GigaScience (2018-07-01) https://doi.org/gfzqf5 DOI: 10.1093/gigascience/giy083 · PMID: 30010766 · PMCID: PMC6057528

81. Prognostic value of grip strength: findings from the Prospective Urban Rural Epidemiology (PURE) study. Darryl P Leong, Koon K Teo, Sumathy Rangarajan, Patricio Lopez-Jaramillo, Alvaro Avezum, Andres Orlandini, Pamela Seron, Suad H Ahmed, Annika Rosengren, Roya Kelishadi, … Lancet (London, England) (2015-05-13) https://www.ncbi.nlm.nih.gov/pubmed/25982160 DOI: 10.1016/s0140-6736(14)62000-6 · PMID: 25982160

82. Densely Interconnected Transcriptional Circuits Control Cell States in Human Hematopoiesis Noa Novershtern, Aravind Subramanian, Lee N Lawton, Raymond H Mak, WNicholas Haining, Marie E McConkey, Naomi Habib, Nir Yosef, Cindy Y Chang, Tal Shay, … Benjamin L Ebert Cell (2011-01) https://doi.org/cf5k92 DOI: 10.1016/j.cell.2011.01.004 · PMID: 21241896 · PMCID: PMC3049864

83. Depression as a predictor for coronary heart disease. a review and meta-analysis. Reiner Rugulies American journal of preventive medicine (2002-07) https://www.ncbi.nlm.nih.gov/pubmed/12093424 DOI: 10.1016/s0749-3797(02)00439-7 · PMID: 12093424

84. Mental Disorders Across the Adult Life Course and Future Coronary Heart Disease Catharine R Gale, GDavid Batty, David PJ Osborn, Per Tynelius, Finn Rasmussen Circulation (2014-01-14) https://doi.org/qm4 DOI: 10.1161/circulationaha.113.002065 · PMID: 24190959 · PMCID: PMC4107269

85. Mortality gap for people with bipolar disorder and schizophrenia: UK-based cohort study 2000–2014 Joseph F Hayes, Louise Marston, Kate Walters, Michael B King, David PJ Osborn British Journal of Psychiatry (2017-09) https://doi.org/gbwcjx DOI: 10.1192/bjp.bp.117.202606 · PMID: 28684403 · PMCID: PMC5579328

86. Getting to the Heart of Alzheimer Disease Joshua M Tublin, Jeremy M Adelstein, Federica del Monte, Colin K Combs, Loren E Wold Circulation Research (2019-01-04) https://doi.org/gjzjgq DOI: 10.1161/circresaha.118.313563 · PMID: 30605407 · PMCID: PMC6319653

87. The overlap between vascular disease and Alzheimer’s disease - lessons from pathology Johannes Attems, Kurt A Jellinger BMC Medicine (2014-11-11) https://doi.org/f6pjd4 DOI: 10.1186/s12916-014-0206-2 · PMID: 25385447 · PMCID: PMC4226890

88. Cardiovascular Risk Factors for Alzheimer’s Disease Clive Rosendorff, Michal S Beeri, Jeremy M Silverman The American Journal of Geriatric Cardiology (2007-03) https://doi.org/bpfw5d DOI: 10.1111/j.1076-7460.2007.06696.x · PMID: 17483665

89. Reverse cholesterol transport and cholesterol elux in atherosclerosis R Ohashi, H Mu, X Wang, Q Yao, C Chen QJM: An International Journal of Medicine (2005-10-28) https://doi.org/dn2fgt DOI: 10.1093/qjmed/hci136 · PMID: 16258026

90. Lipid and Lipoprotein Metabolism in Microglia Bailey A Loving, Kimberley D Bruce Frontiers in Physiology (2020-04-28) https://doi.org/gk92xd DOI: 10.3389/fphys.2020.00393 · PMID: 32411016 · PMCID: PMC7198855

91. Shared genetic origin of asthma, hay fever and eczema elucidates allergic disease biology Manuel A Ferreira, Judith M Vonk, Hansjörg Baurecht, Ingo Marenholz, Chao Tian, Joshua D Hoffman, Quinta Helmer, Annika Tillander, Vilhelmina Ullemar, … Nature Genetics (2017-10-30) https://doi.org/gchg62 DOI: 10.1038/ng.3985 · PMID: 29083406 · PMCID: PMC5989923

92. A genome-wide cross-trait analysis from UK Biobank highlights the shared genetic architecture of asthma and allergic diseases Zhaozhong Zhu, Phil H Lee, Mark D Chain, Wonil Chung, Po-Ru Loh, Quan Lu, David C Christiani, Liming Liang Nature Genetics (2018-05-21) https://doi.org/gdpmtn DOI: 10.1038/s41588-018-0121-0 · PMID: 29785011 · PMCID: PMC5980765

93. recount3: summaries and queries for large-scale RNA-seq expression and splicing Christopher Wilks, Shijie C Zheng, Feng Yong Chen, Rone Charles, Brad Solomon, Jonathan P Ling, Eddie Luidy Imada, David Zhang, Lance Joseph, Jeffrey T Leek, … Ben Langmead Cold Spring Harbor Laboratory (2021-05-23) https://doi.org/gj7cmq DOI: 10.1101/2021.05.21.445138

94. GenomicSuperSignature facilitates interpretation of RNA-seq experiments through robust, eficient comparison to public databases Sehyun Oh, Ludwig Geistlinger, Marcel Ramos, Daniel Blankenberg, Marius van den Beek, Jaclyn N Taroni, Vincent J Carey, Casey S Greene, Levi Waldron, Sean Davis Nature Communications (2022-06-27) https://doi.org/gqd7hm DOI: 10.1038/s41467-022-31411-3 · PMID: 35760813 · PMCID: PMC9237024

95. Opportunities and challenges for transcriptome-wide association studies Michael Wainberg, Nasa Sinnott-Armstrong, Nicholas Mancuso, Alvaro N Barbeira, David A Knowles, David Golan, Raili Ermel, Arno Ruusalepp, Thomas Quertermous, Ke Hao, … Anshul Kundaje Nature Genetics (2019-03-29) https://doi.org/gf3hmr DOI: 10.1038/s41588-019-0385-z · PMID: 30926968 · PMCID: PMC6777347

96. Probabilistic colocalization of genetic variants from complex and molecular traits: promise and limitations Abhay Hukku, Milton Pividori, Francesca Luca, Roger Pique-Regi, Hae Kyung Im, Xiaoquan Wen The American Journal of Human Genetics (2021-01) https://doi.org/gj58gg DOI: 10.1016/j.ajhg.2020.11.012 · PMID: 33308443 · PMCID: PMC7820626

97. Transcriptome-wide association study of schizophrenia and chromatin activity yields mechanistic disease insights Alexander Gusev, Nicholas Mancuso, Hyejung Won, Maria Kousi, Hilary K Finucane, Yakir Reshef, Lingyun Song, Alexias Safi, Steven McCarroll, Benjamin M Neale, … Nature Genetics (2018-04) https://doi.org/gdfdf2 DOI: 10.1038/s41588-018-0092-1 · PMID: 29632383 · PMCID: PMC5942893

98. Systematic tissue annotations of –omics samples by modeling unstructured metadata Nathaniel T Hawkins, Marc Maldaver, Anna Yannakopoulos, Lindsay A Guare, Arjun Krishnan Cold Spring Harbor Laboratory (2021-05-11) https://doi.org/gj2pkc DOI: 10.1101/2021.05.10.443525

99. Exploring the phenotypic consequences of tissue specific gene expression variation inferred from GWAS summary statistics Alvaro N Barbeira, Scott P Dickinson, Rodrigo Bonazzola, Jiamao Zheng, Heather E Wheeler, Jason M Torres, Eric S Torstenson, Kaanan P Shah, Tzintzuni Garcia, … Hae Kyung Im Nature Communications (2018-05-08) https://doi.org/gdjvp5 DOI: 10.1038/s41467-018-03621-1 · PMID: 29739930 · PMCID: PMC5940825

100. The Molecular Signatures Database Hallmark Gene Set Collection Arthur Liberzon, Chet Birger, Helga Thorvaldsdóttir, Mahmoud Ghandi, Jill P Mesirov, Pablo Tamayo Cell Systems (2015-12) https://doi.org/gf78hq DOI: 10.1016/j.cels.2015.12.004 · PMID: 26771021 · PMCID: PMC4707969

101. MAGMA: Generalized Gene-Set Analysis of GWAS Data Christiaan A de Leeuw, Joris M Mooij, Tom Heskes, Danielle Posthuma PLOS Computational Biology (2015-04-17) https://doi.org/gf92gp DOI: 10.1371/journal.pcbi.1004219 · PMID: 25885710 · PMCID: PMC4401657

102. Human Disease Ontology 2018 update: classification, content and worklow expansion Lynn M Schriml, Elvira Mitraka, James Munro, Becky Tauber, Mike Schor, Lance Nickle, Victor Felix, Linda Jeng, Cynthia Bearer, Richard Lichenstein, … Carol Greene Nucleic Acids Research (2018-11-08) https://doi.org/ggx9wp DOI: 10.1093/nar/gky1032 · PMID: 30407550 · PMCID: PMC6323977

103. Modeling sample variables with an Experimental Factor Ontology James Malone, Ele Holloway, Tomasz Adamusiak, Misha Kapushesky, Jie Zheng, Nikolay Kolesnikov, Anna Zhukova, Alvis Brazma, Helen Parkinson Bioinformatics (2010-03-03) https://doi.org/dsb6vt DOI: 10.1093/bioinformatics/btq099 · PMID: 20200009 · PMCID: PMC2853691

104. GitHub - EBISPOT/EFO-UKB-mappings GitHub https://github.com/EBISPOT/EFO-UKB-mappings

105. Comparing partitions Lawrence Hubert, Phipps Arabie Journal of Classification (1985-12) https://doi.org/bphmzh DOI: 10.1007/bf01908075

106. Diversity control for improving the analysis of consensus clustering Milton Pividori, Georgina Stegmayer, Diego H Milone Information Sciences (2016-09) https://doi.org/ghtqbk DOI: 10.1016/j.ins.2016.04.027

107. A Link-Based Approach to the Cluster Ensemble Problem Natthakan Iam-On, Tossapon Boongoen, Simon Garrett, Chris Price IEEE Transactions on Pattern Analysis and Machine Intelligence (2011-12) https://doi.org/cqgkh3 DOI: 10.1109/tpami.2011.84 · PMID: 21576752

108. Hybrid clustering solution selection strategy Zhiwen Yu, Le Li, Yunjun Gao, Jane You, Jiming Liu, Hau-San Wong, Guoqiang Han Pattern Recognition (2014-10) https://doi.org/ghtzwt DOI: 10.1016/j.patcog.2014.04.005

109. UMAP: Uniform Manifold Approximation and Projection for Dimension Reduction Leland McInnes, John Healy, James Melville arXiv (2020-09-21) https://arxiv.org/abs/1802.03426

110. k-means++: the advantages of careful seeding David Arthur, Sergei Vassilvitskii Proceedings of the Eighteenth Annual ACM-SIAM Symposium on Discrete Algorithms (2007) http://ilpubs.stanford.edu:8090/778/1/2006-13.pdf

111. On Spectral Clustering: Analysis and an algorithm Andrew Ng, Michael Jordan, Yair Weiss Advances in Neural Information Processing Systems (2001) https://ai.stanford.edu/~ang/papers/nips01-spectral.pdf

112. A Density-Based Algorithm for Discovering Clusters in Large Spatial Databases with Noise Martin Ester, Hans-Peter Kriegel, Jörg Sander, Xiaowei Xu Proceedings of the Second International Conference on Knowledge Discovery and Data Mining (1996) https://www.aaai.org/Papers/KDD/1996/KDD96-037.pdf

113. Determination of Optimal Epsilon (Eps) Value on DBSCAN Algorithm to Clustering Data on Peatland Hotspots in Sumatra Nadia Rahmah, Imas Sukaesih Sitanggang IOP Conference Series: Earth and Environmental Science (2016-01) https://doi.org/gqr7z2 DOI: 10.1088/1755-1315/31/1/012012

114. Avoiding common pitfalls when clustering biological data. Tom Ronan, Zhijie Qi, Kristen M Naegle Science signaling (2016-06-14) https://www.ncbi.nlm.nih.gov/pubmed/27303057 DOI: 10.1126/scisignal.aad1932 · PMID: 27303057

115. Optimized sgRNA design to maximize activity and minimize off-target effects of CRISPR-Cas9. John G Doench, Nicolo Fusi, Meagan Sullender, Mudra Hegde, Emma W Vaimberg, Katherine F Donovan, Ian Smith, Zuzana Tothova, Craig Wilen, Robert Orchard, … David E Root Nature biotechnology (2016-01-18) https://www.ncbi.nlm.nih.gov/pubmed/26780180 DOI: 10.1038/nbt.3437 · PMID: 26780180 · PMCID: PMC4744125

116. Fine-mapping and QTL tissue-sharing information improves the reliability of causal gene identification Alvaro N Barbeira, Owen J Melia, Yanyu Liang, Rodrigo Bonazzola, Gao Wang, Heather E Wheeler, François Aguet, Kristin G Ardlie, Xiaoquan Wen, Hae K Im Genetic Epidemiology (2020-09-10) https://doi.org/gqsvf7 DOI: 10.1002/gepi.22346 · PMID: 32964524 · PMCID: PMC7693040

117. GitHub - hakyimlab/summary-gwas-imputation: harmonization, liftover, and imputation of summary statistics from GWAS GitHub https://github.com/hakyimlab/summary-gwas-imputation

118. Homo sapiens (ID 232177) - BioProject - NCBI https://www.ncbi.nlm.nih.gov/bioproject/PRJNA232177

